# TransPi – a comprehensive TRanscriptome ANalysiS PIpeline for *de novo* transcriptome assembly

**DOI:** 10.1101/2021.02.18.431773

**Authors:** R.E. Rivera-Vicéns, C.A. Garcia-Escudero, N. Conci, M. Eitel, G. Wörheide

## Abstract

The use of RNA-Seq data and the generation of *de novo* transcriptome assemblies have been pivotal for studies in ecology and evolution. This is distinctly true for non-model organisms, where no genome information is available. Nevertheless, studies of differential gene expression, DNA enrichment baits design, and phylogenetics can all be accomplished with the data gathered at the transcriptomic level. Multiple tools are available for transcriptome assembly, however, no single tool can provide the best assembly for all datasets. Therefore, a multi assembler approach, followed by a reduction step, is often sought to generate an improved representation of the assembly. To reduce errors in these complex analyses while at the same time attaining reproducibility and scalability, automated workflows have been essential in the analysis of RNA-Seq data. However, most of these tools are designed for species where genome data is used as reference for the assembly process, limiting their use in non-model organisms. We present TransPi, a comprehensive pipeline for *de novo* transcriptome assembly, with minimum user input but without losing the ability of a thorough analysis. A combination of different model organisms, k-mer sets, read lengths, and read quantities were used for assessing the tool. Furthermore, a total of 49 non-model organisms, spanning different phyla, were also analyzed. Compared to approaches using single assemblers only, TransPi produces higher BUSCO completeness percentages, and a concurrent significant reduction in duplication rates. TransPi is easy to configure and can be deployed seamlessly using Conda, Docker and Singularity.

## 1. Introduction

In the last decades, technology improvements have rendered next generation sequencing (NGS) a robust and cost effective technique of wide applicability in research fields that require large-scale DNA sequencing. Among the different NGS-based approaches, RNA sequencing (RNA-Seq) allows the generation of the so-called transcriptomes *de novo* (i.e., without the need for a reference genome). Transcriptomes are applicable for several downstream applications, including the analysis of differential gene expression (Pita et al., 2018), gene model prediction (Chan et al., 2017), DNA enrichment baits design (Quek et al., 2020), genome annotation (Testa et al., 2015; Holt and Yandell, 2011), detection of whole-genome duplication (Yang et al., 2019), and phylogenetics (Cheon et al., 2020; Lozano-Fernandez et al., 2019).

Various software have been developed for the generation of *de novo* transcriptome assembly. Commonly used tools include Trinity (Grabherr et al., 2011), rnaSPADES (Bushmanova et al., 2019), Trans-ABySS (Robertson et al., 2010), and SOAPdenovo-Trans (Xie et al., 2014). However, a recent study compared ten assemblers with nine datasets (i.e. different species and samples) and demonstrated that the performance of each tool varies by dataset; no single tool was able to generate optimal assemblies for all datasets (Hölzer & Marz 2019). The assemblers performance measurement was based on a combination of biological based measures (e.g. number of Benchmarking Universal Single-Copy Orthologs - BUSCO), and reference-free measures (e.g. TransRate’s optimal assembly score; Smith-Unna et al., 2016). Therefore, combining multiple assemblers likely represents a valuable approach to increase the quality of reference assemblies (Lu et al., 2013). Additionally, factors such as reads length and number also play important roles in the assembly process (Grabherr et al., 2011; Schulz et al., 2012; Francis et al., 2013).

Transcriptome *de novo* assemblies tend to produce thousands to hundreds of thousands of different transcripts of which a significant amount can be misassembled (Bushmanova et al., 2019; Schulz et al., 2012). To reduce the complexity within a transcriptome and to identify true transcripts and isoforms, one common approach is to remove duplicated and misassembled sequences. Clustering methods are often employed for this, where similar transcripts are combined into groups. One of the tools commonly used for clustering transcripts is CD-HIT-EST (Fu et al., 2012), which tends to keep the longest transcripts only. However, clustering and selecting for the longest transcripts is not always the best strategy (Gilbert, 2013) since they often result from misassemblies (i.e. not real transcripts) and may include frameshift errors. On the other hand, tools such as EvidentialGene (Gilbert, 2013, 2019) use a combination of clustering and classification methods (i.e, sequence features like coding sequence (CDS) content and length) to generate a non-redundant consensus assembly. The latter approach is more accurate for the cost of longer computing time and higher computation demands (e.g. higher memory usage). Combining multiple assemblers with a thorough reduction of each assembly individually thus increases the complexity of the analyses.

The ideal path to optimal reference transcriptomes should, therefore, include the use of multiple assemblers, followed by thorough filtering of each assembly individually. Generating, combining and filtering all resulting assemblies step by step individually (cf. MacManes, 2018; Cerveau, & Jackson, 2016) is impractical, of limited reproducibility, and can be prone to human error. Consequently, the design of streamlined RNA-Seq analysis pipelines have gained popularity over the recent years. However, most of these pipelines require a reference genome for the transcriptome assembly (i.e., reference-guided assembly) and are, consequently, not suitable for *de novo* approaches (Cornwell et al., 2018; D’Antonio et al., 2015; Wang, D. 2018; Zhang, X., & Jonassen, I. 2020; Kohen et al., 2019; Martin et al., 2010). This represents a major limitation for transcriptomics in non-model organisms, where genome reference data is usually lacking.

To address these shortcomings, we developed TransPi, a comprehensive TRanscriptome ANalysiS Pipeline for *de novo* transcriptome assembly. TransPi is implemented using the scientific workflow manager Nextflow (Di Tommaso et al., 2017), which provides a user-friendly environment, easy deployment, scalability and reproducibility. TransPi performs all steps of standard RNA-Seq analysis workflows, from raw reads quality control up to annotation against multiple databases (e.g. SwissProt, PFAM). To reduce possible biases, duplication and misassemblies, TransPi utilizes various assemblers and k-mers (i.e. k length sequences used for the assembly) to generate an over assembled transcriptome that is then reduced to a non-redundant consensus transcriptome with the software EvidentialGene (Gilbert, 2013, 2019). Here we show that, when compared to approaches using single assemblers only, TransPi produces higher BUSCO completeness percentages, and a concurrent significant reduction in duplication rates (i.e. higher single-copy genes). Higher BUSCO scores in the complete and single-copy categories indicates a less erroneous consensus assembly (Waterhouse et al., 2011; Simão, et al., 2015).

In sum, TransPi provides a useful resource for the generation of *de novo* transcriptome assemblies, with minimum user input but without losing the ability of a thorough analysis. TransPi and all documentation is available at https://github.com/palmuc/TransPi.git.

## 2. Methods

### 2.1 Pipeline implementation and configuration

TransPi is based on the scientific workflow manager Nextflow (Di Tommaso et al., 2017). The pipeline is easy to configure and can be deployed using the package management system Conda, Docker, Singularity or cloud environments (e.g., AWS). Real-time monitoring of the pipeline can be performed by using Nextflow Tower with no modification needed to the TransPi script. Deployment of TransPi in computing clusters is accomplished by the native communication of Nextflow with scheduling managers such as SLURM, PBS and Torque. TransPi can deploy hundreds of jobs depending on user configurations and needs. Multiple datasets can be run in parallel given that enough computing resources are available. Running time of the pipeline is dependent on factors such as number of datasets, reads quantity, k-mer selection, complexity of the transcriptome being assembled, and user-specified additional options selected (e.g. filtration, signalP, etc.). TransPi consists of two major components: a precheck script to install dependencies, and the main script to run the assemblers, perform the reduction and transcriptome annotation.

### 2.2 Precheck script

TransPi integrates several programs and external databases (e.g. SwissProt, Boeckmann et al., 2003; PFAM, El-Gebali et al., 2019) for the generation and annotation of the reference transcriptome. To facilitate the setup of all necessary dependencies, TransPi includes an installation script. This will first install, if necessary, the Conda package management system, all dependencies, and download and configure the required databases. The script is designed to recognize when a previous run of the script was performed, thus preventing the repetition of previous steps. Another advantage of the precheck script is that it will automatically create the configuration file needed by Nextflow to execute the pipeline with all the necessary information. As a result, the user will only have to make some minor changes to the file (e.g. number of allocated threads, the amount of working memory, scheduling manager, node names and queue) before running the pipeline. Essentially, the precheck has to be run entirely only one time for the dependencies and database installation. Subsequent pipeline runs can be done with the same configuration file. Auxiliary scripts for the automated update of the databases such as Pfam and SwissProt are also provided.

### 2.3 Main script

A diagram of the TransPi v1.0.0 is shown in Figure 1. First, reads are checked for adapter presence and/or low quality bases with FastQC v0.11.9 (Andrews, 2010). Filtration of the reads (by default reads with an average phred quality >25 are kept) and trimming of adapters (if present) is performed with fastp v0.20.1 (Chen et al., 2018). Optionally, removal of ribosomal RNA (rRNA) is performed with SortMeRNA v4.2.0 (Kopylova et al., 2012). Filtered reads are subsequently normalized before being assembled (Grabherr et al., 2011). The assembly step combines five different assemblers and uses multiple k-mer lengths. The assemblers used by TransPi are rnaSPADES v3.14.0 (Bushmanova et al., 2019), Trans-ABySS v2.0.1 (Robertson et al., 2010), SOAPdenovo-Trans v1.03 (Xie et al., 2014), Trinity v2.9.1 (Grabherr et al., 2011) and Velvet v1.2.12/Oases v0.2.09 (Zerbino & Birney, 2008; Schulz et al., 2012). After the assembly stage, the combined transcriptomes are reduced with EvidentialGene v2019.05.14 (Gilbert, 2013, 2019). Briefly, EvidentialGene will merge perfect duplicates, cluster protein sequences, and perform local similarities searches between the transcripts using BLAST v2.2.31 (Altschul et al., 1997) (for more details see Gilbert, 2019).

**Figure 1.**
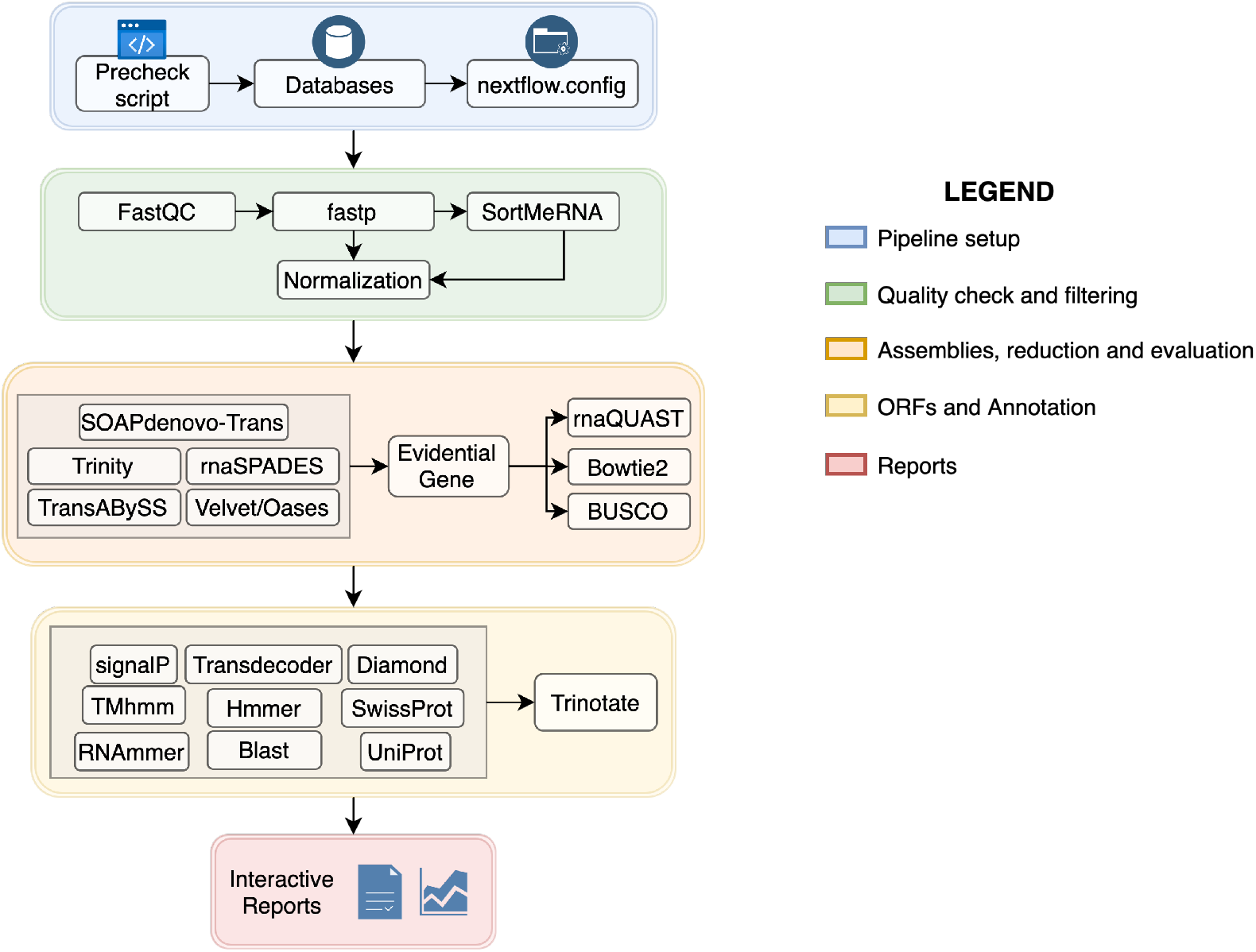
TransPi v1.0.0 flowchart showing the various steps and analyses it can performed. For simplicity, this diagram does not show all the connections between the processes. Also, it omits other additional options like the BUSCO distribution and transcriptome filtering with psytrans (see Section 2.6). ORFs=Open reading Frames

Next, TransPi uses the non-redundant reference transcriptome to run several downstream analyses commonly applied to *de novo* transcriptomes projects: 1) rnaQUAST v2.0.1 for quality assessment (Bushmanova et al., 2016), 2) Bowtie2 v2.3.5.1 to map the reads against the transcriptome (Langmead & Salzberg, 2012), 3) BUSCO (Simao et al., 2015; v3 and v4) to quantitatively assess the completeness in terms of expected universal single copy gene content, 4) TransDecoder v5.5.0 (https://transdecoder.github.io) to identify open reading frames (ORFs), with the option to perform homology searches of all ORFs to known proteins via BLAST, in order to retain ORFs that may have functional significance but don’t pass the coding likelihood scores, and 5) Trinotate v3.2.0 (Bryant et al., 2017) to provide automatic functional annotation.

By using Diamond v0.9.30 (Buchfink et al., 2015), the similarity searches of the transcripts used for the annotation step against the SwissProt and UniProt databases (chosen by the user) are accelerated. RNAmmer v1.2 (Lagesen et al., 2007), TMhmm v2.0 (Krogh et al., 2001), SignalP 4.1 (Petersen et al., 2011) are used to search for ribosomal RNA, signal peptide proteins, and transmembrane domain prediction, respectively. Protein domain searches are performed with HMMER v3.3 (Finn et al., 2011) against the last version of the PFAM database. All this information is combined into an annotation report which includes: 1) information on Gene Ontology (GO); 2) evolutionary genealogy of genes: Non-supervised Orthologous Groups (eggNOG), and; 3) Kyoto Encyclopedia of Genes and Genomes (KEGG). It also contains the similarity search against SwissProt and the user-specified UniProt database. TransPi will also produce a custom Hypertext Markup Language (HTML) report that summarizes the steps and provides interactive plots for straightforward exploration of the data. Plots from the interactive report can also be saved in SVG format. Other plots are also saved automatically (PDF and SVG) in the results directory generated by the pipeline. Altogether, TransPi provides the user with the ability to assess and evaluate the final assembly and to compare it to other commonly used methods for reference transcriptome generation (e.g., a Trinity only assembly).

### 2.4 K-mer selection, Reads length effect and Chimera detection

To test the performance of the pipeline and the effect of k-mer selection (Prjibelski et al., 2020), read quantities, and read lengths, datasets from the model organism *Caenorhabditis elegans, Drosophila melanogaster*, and *Mus musculus* were used. These species were selected given the vast amount of transcriptomic data available with various read length and quantities (Supplementary Table 1). Three k-mer sets (A, B, C) depending on read length were designed, since the selection of this parameter will modify how the assembly graph is constructed. For the read length test, data consisting of paired-end reads of 50 base-pairs (bp), 75 bp, 100 bp, 150 bp (Supplementary Table 1) were analysed. All statistical analyses, such as ANOVA and Kruskal-Wallis test, were performed in R (v3.6.2).

To measure the percentage of chimeric transcripts and transcript accuracy a similar approach to Kerkvliet et al. 2019 was used. First, gene sets for model organisms *Caenorhabditis elegans* (i.e. c_elegans.PRJNA13758.WS279.mRNA_transcripts.fa from Wormbase), *Drosophila melanogaster* (i.e. Dmel-all-transcript-r6.39.fasta from Flybase), and *Mus musculus* (i.e. GCF_000001635.27_GRCm39_rna_from_genomic.fna from NCBI) were downloaded. Then a BLASTN search was performed using the transcriptomes from TransPi and Trinity against each corresponding gene set. Parameters used were as specified by Kerkvliet et al., 2019 (-perc_identity .90 -evalue .001). BLASTN output was filtered using a minimum length of 300bp for each match. Non-chimeric transcripts were identified as transcripts with one match per gene. Transcripts with two or more matches were classified as chimeras.

### 2.6 Additional options

Various additional options were implemented in TransPi to obtain more insight into the transcriptomes being assembled. One of these options is filtering symbionts and/or contaminants from the assembly using the Parasite & Symbiont Transcriptome Separation software (psytrans) (https://github.com/sylvainforet/psytrans). The filtration step was tested with the dataset of the coral *Porites pukoensis* (accession SRR8491966) using sequences of it symbiont *Symbiodinium microadriaticum* (Uniprot Taxon Identifier: 2951) and sequences of the order Scleractinia (Uniprot Taxon Identifier: 6125) as host. Another option of TransPi examines the presence and absence of BUSCO genes in all the generated assemblies and creates a heatmap of genes distribution. This option was tested with the epadomorph barnacle *Octolasmis warwickii* dataset (SRR10527303) given the difference between TransPi and Trinity BUSCO scores for the missing category (see Results).

## 3 Results

### 3.1 K-mer selection, Reads length effect and Chimera detection

K-mer test carried out on the model organisms used here (Supplementary Table 1) showed that differences in BUSCO percentages between k-mer sets (i.e. A, B, C) were not significant (Supplementary Table 2). However, slightly higher single-copy and lower duplication BUSCO percentages were observed with k-mer set C (Figure 2; Supplementary Table 2; Supplementary File 1-4). This pattern was observed in all three model organisms: *Caenorhabditis elegans* (worm), *Drosophila melanogaster* (fly), and *Mus musculus* (mouse) (Supplementary File 1-4). The read length test (i.e. 50 bp, 75 bp, 100 bp, and 150 bp) also showed no significant difference in complete BUSCO percentages (Figure 2; Supplementary File 1-4). However, it should be noted that *D. melanogaster* 50 bp paired-end reads produced low complete BUSCO percentages for TransPi and Trinity (Complete BUSCO mean <45%). On the contrary, *D. melanogaster* libraries with paired-end reads of 75 bp, 100 bp and 150 bp length showed high BUSCO percentages for both, TransPi and Trinity, where Trinity surpasses TransPi by 1.0% (Figure 2; Supplementary File 1-4). A similar pattern of a marginal difference between TransPi and Trinity (with 1.0% higher complete BUSCO percentage in Trinity) was also observed for the *C. elegans* and *M. musculus* datasets (Supplementary File 1-4).

**Figure 2.**
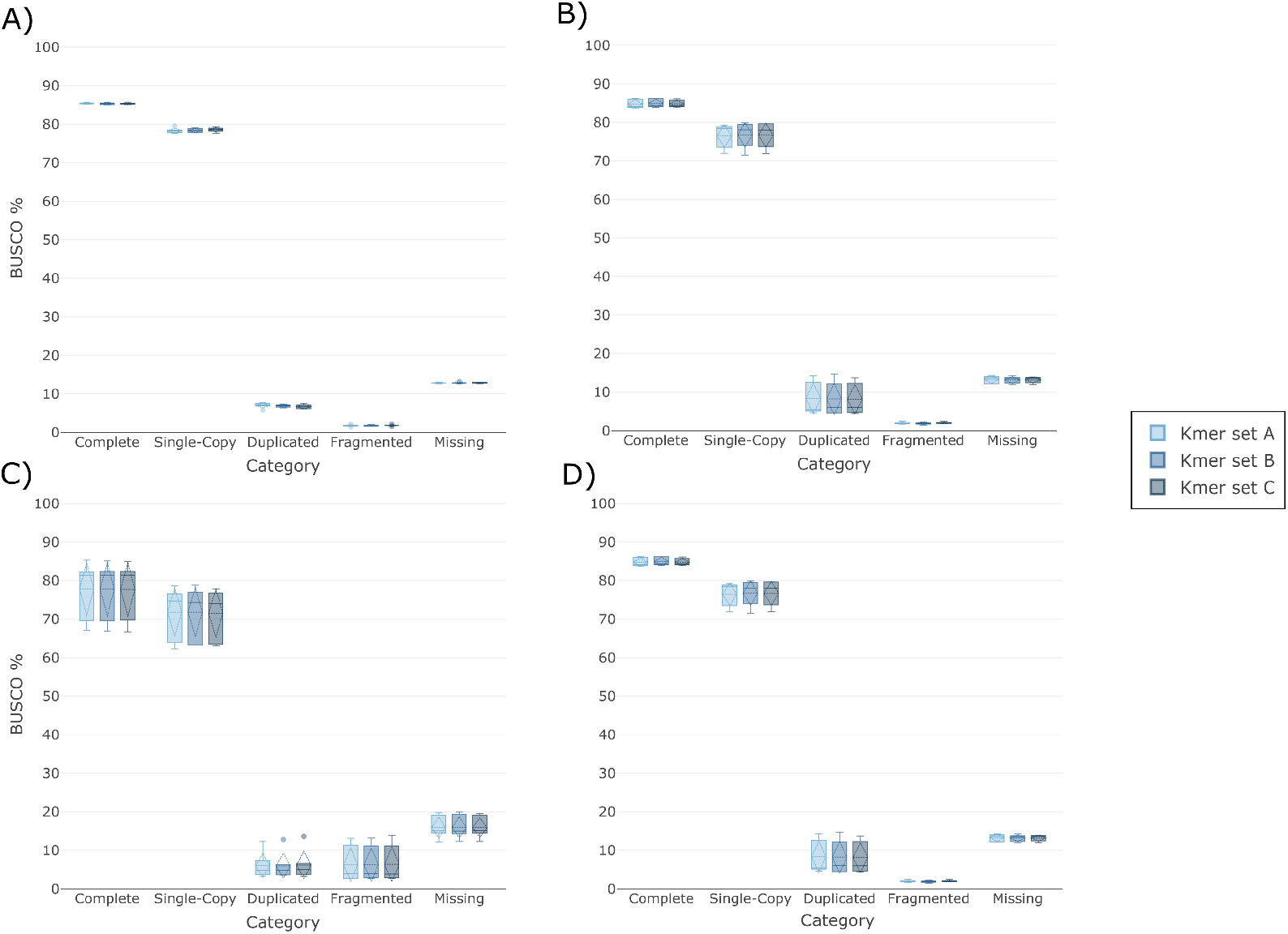
K-mers selection tests on the model organisms *C. elegans*,. Shown are the TransPi results for three different k-mer settings for read lengths of 50bp (A), 75bp (B), 100bp (C), and 150bp (D). For the k-mer test performed with *M. musculus* and *D. melanogaster* see Supplementary Files 1-4.

The major difference between TransPi and Trinity in the model organisms was observed in the single-copy BUSCO category. This difference was more significant in the *D. melanogaster* and *M. musculus* datasets. For the *M. musculus* 150 bp reads, the difference between the single-copy BUSCO percentages was over 37% (Supplementary Table 2). In terms of fragmented and missing BUSCO genes, TransPi scores were slightly higher (<1.0%) than for Trinity alone in most cases (Supplementary Table 2; Supplementary File 1-4). The increase of read length showed no clear effect on producing better BUSCO percentages on the majority of the model organisms datasets (Supplementary File 1-4). The same was observed for the increase of the read quantities in the datasets (Supplementary File 1-4). Only for *D. melanogaster* 50 bp reads, an increase was observed in complete BUSCO percentages when incrementing reads quantity from 10M to 26M. The other model organisms datasets did not show significant differences with respect to read quantities (Supplementary File 1-4).

Results for the chimera detection test are presented in Table 2. A similar trend was observed in all model species (i.e. *C. elegans, D. melanogaster*, and *M. musculus*). The number of non-chimeric transcripts (i.e. % of unique BLASTN matches) in TransPi (i.e. lowest: 3.07% - highest: 39.13%) was higher than in Trinity alone (i.e. lowest: 3.66% - highest: 38.32%). Only in one sample (i.e. *M. musculus* SRR8329326) the percentage of non-chimeric transcripts of Trinity was higher than TransPi. However, the Trinity assembly had over 215,000 more transcripts than the TransPi transcriptome. Nevertheless, the percentage difference was only 0.59%. BUSCO scores followed the same pattern as explained above.

### 3.2 TransPi on non-model organisms

A similar trend as seen in the model organisms was observed in the non-model organisms datasets (Figure 3; Supplementary Table 3). However, there were some key differences. First, results of complete BUSCO percentages were higher for TransPi in 41 of the 49 datasets tested in the study. The mean of complete BUSCO percentages was 79.57%±18.60 (median: 85%) for TransPi and 78.14%±19.30 (median: 84.2%) for the Trinity assemblies. Of all datasets, 21 had complete BUSCO percentages higher than 90% with TransPi and 17 with Trinity (Figure 4). Eleven and 13 datasets resulted in 80-90% identified complete BUSCO genes with TransPi and Trinity, respectively. However, Kruskal-Wallis test showed no significant differences between TransPi and Trinity (Table 3).

**Figure 3.**
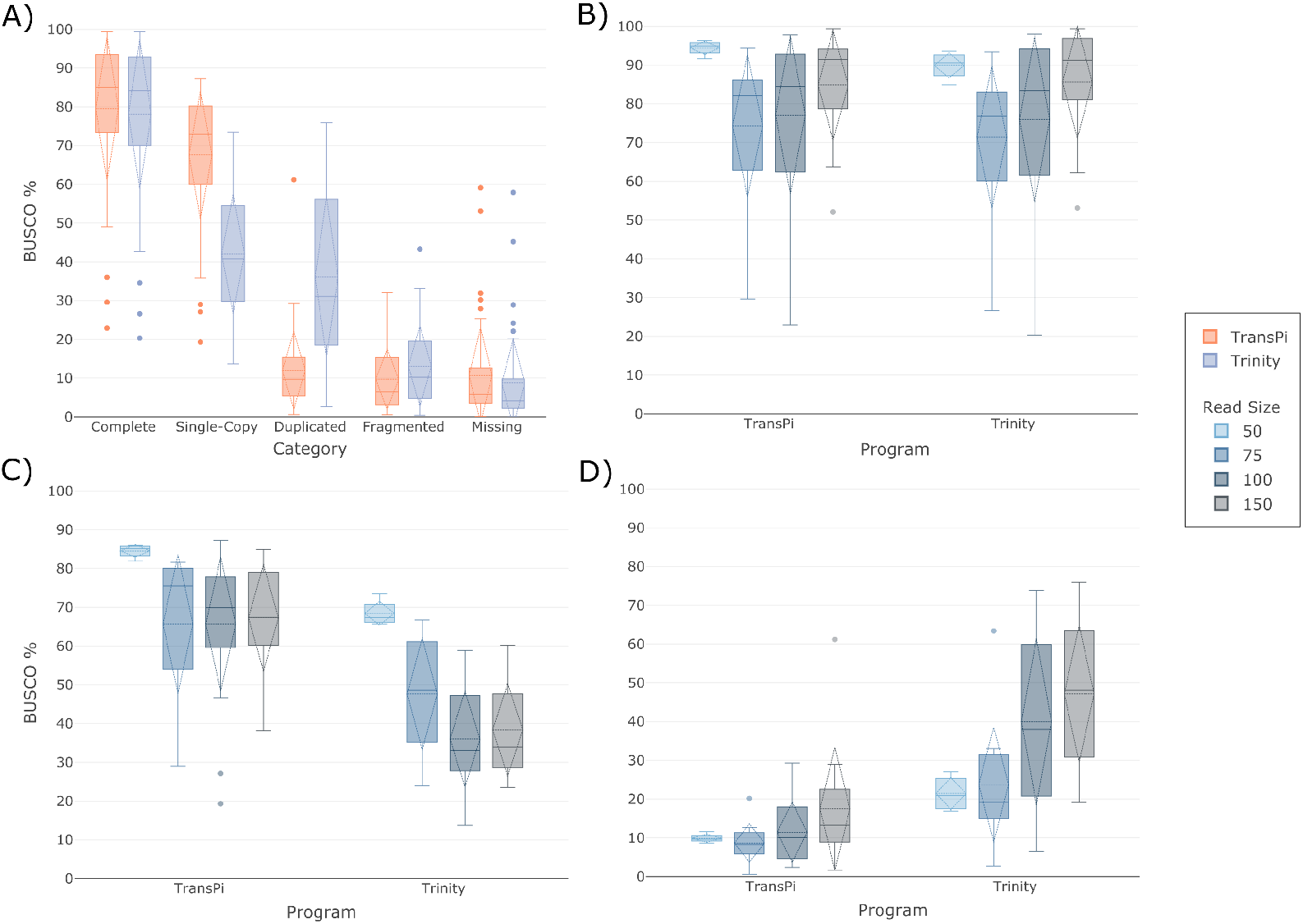
BUSCO results of non-model organisms (n=49). A full list of analysed taxa see table 2. A) BUSCO percentages comparison for TransPi and Trinity for all datasets. Comparisons of scores by read length for complete (B), single-copy (C), and duplicated (D) BUSCO genes. Significant differences (Kruskal-Wallis test p <0.05) were obtained for (B) and (C).

**Figure 4.**
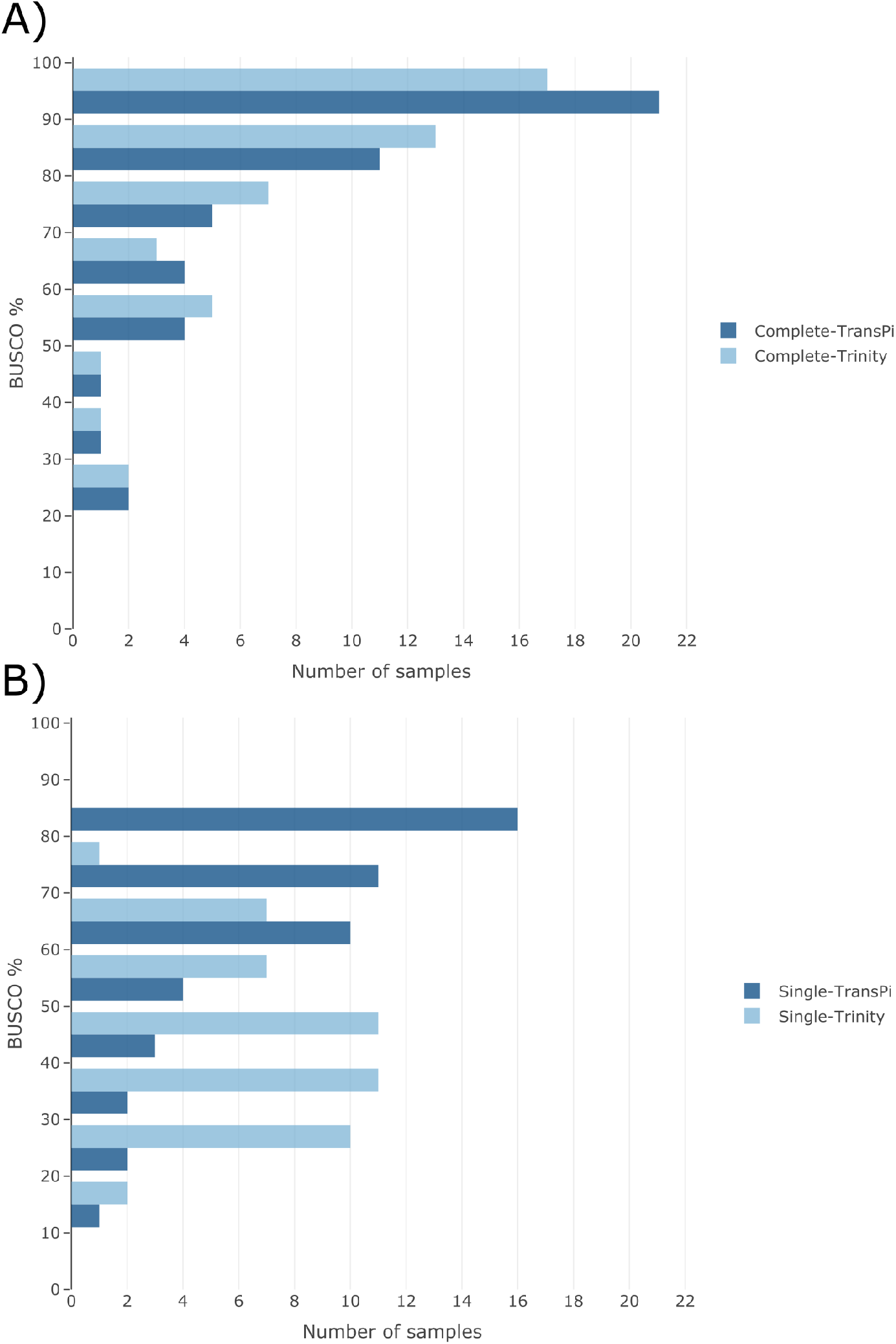
Histogram of number of datasets and BUSCO percentages in 10% bins. Comparison of identified complete (including duplicates) (A) and single-copy (B) BUSCO genes between TransPi and Trinity

Second, there was a significant improvement of the percentage of identified complete single-copy BUSCO genes. Mean percentages with TransPi and Trinity were 67.57±16.75% (median: 72.9%) and 42.03±15.37% (median: 40.8%), respectively. For the single-copy BUSCO genes, 16 datasets obtained scores higher than 80% with TransPi and none with Trinity (Figure 4). For the range of 70-80%, 11 datasets obtained scores in this range when using TransPi, whereas only one dataset in this range was obtained when using Trinity (Figure 4). Statistical test (i.e. Kruskal-Wallis) demonstrated a significant difference for the single-copy BUSCO percentages between TransPi and Trinity (p-value 5.6e-10, Table 3). In the case of the nemertean worm *Malacobdella grossa* (accession SRR1611560), single-copy BUSCO genes had a substantial change from 20.4% for Trinity to 83% with TransPi (Supplementary File 5). Other dataset with significant changes included the crinoid echinoderm *Florometra serratissima* (accession SRR3097584), where the scores for Trinity and TransPi were 41.3% and 87.3%, respectively (Supplementary File 5).

Through the reduction step performed by EvidentialGene in TransPi, an expected substantial decrease of the duplication rate was observed. The means for duplicated BUSCO genes with TransPi and Trinity were 12.0±9.96% (median: 9.7%) and 36.11±20.52% (median: 31.1%), respectively (Figure 3,4; Supplementary Table 3). Kruskal-Wallis tests demonstrated a significant difference for the duplicated BUSCO percentages (p-value of 9.60e-11, Table 3). Even though differences in fragmented BUSCO percentages were not statistically significant, these values were lower for datasets when using TransPi. In the case of missing BUSCO percentages, TransPi scores are higher than Trinity (Supplementary File 5), although differences were not significant. These genes were removed during the reduction step of EvidentialGene (see Discussion). It should be noted that a few datasets were encountered where neither TransPi nor Trinity obtained complete BUSCO percentages higher than 50%. These datasets are: the polychaete annelid *Nephtys caeca* (accession SRR1232685), and the bivalve molluscs *Mercenaria campechiensis* (accession SRR1560359), *Sphaerium nucleus* (accession SRR1561723), and *Cardites antiquatus* (accession SRR1560458) (Supplementary Table 3). However, the majority of the identified complete BUSCO genes in these sets were single-copy in the TransPi assemblies (Supplementary File 5). On the other hand, datasets such as the scleractinian coral *Porites pukoensis* (a accession SRR8491966) were observed with complete BUSCO percentages of 99.4% with both TransPi and Trinity (with high duplication rates in both).

As expected due to the transcripts reduction, the total number of transcripts in TransPi was lower than with Trinity (Supplementary File 6). The mean for TransPi transcripts was 93,351 ± 89,863 (median: 73,435) and for Trinity transcripts 157,130 ± 142,410) (median: 109,261). The reduction of transcript was also observed for the numbers of transcripts larger than 500 bp and 1,000 bp (Figure 5). However, in terms of the longest transcript, the mean for TransPi was 23,684 bp ± 15,374 bp (median: 22,147 bp) and 20,668 bp ± 11,248 bp (median: 18,708 bp) for Trinity (Figure 5). Mapping of sequencing reads to the assembled transcripts showed lower mapping rates obtained with TransPi than with Trinity (Figure 5; Supplementary File 7). The mean of the predicted genes by TransPi and Trinity was 34,659±43,987 (median: 25,280) and 52,106±47,273 (median: 43,783), respectively (Figure 5; Supplementary File 6). This reduction in TransPi vs Trinity mirrors the reduction of duplicated BUSCOs results.

**Figure 5.**
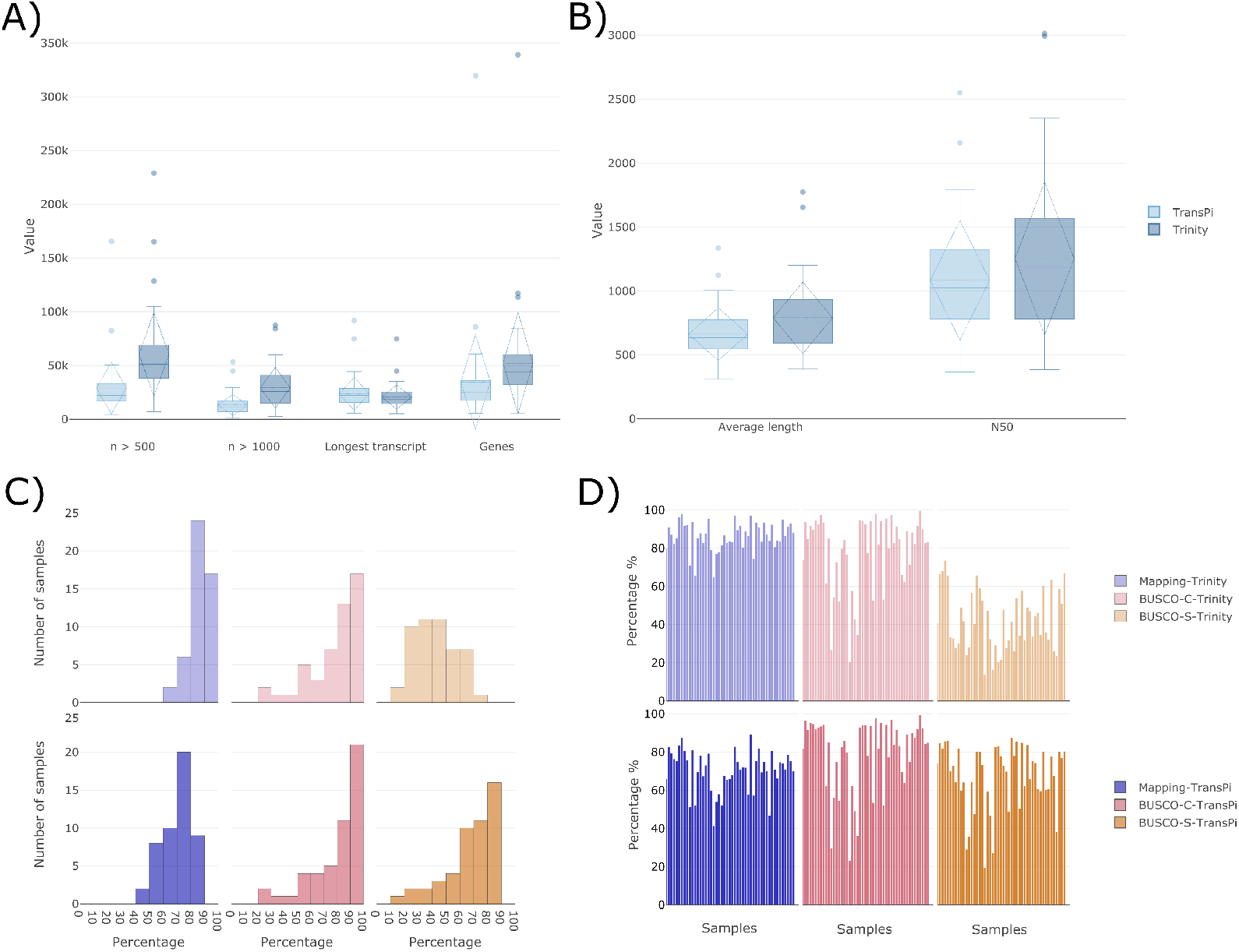
rnaQUAST results comparing TransPi and Trinity. A) Number of transcripts higher than 500bp, 1000bp, longest transcripts and number of predicted genes in the transcriptome. B) Transcripts average length and N50. C) Histograms (10% bins) of all samples and percentage of mapping (reads to transcriptome), BUSCO complete and BUSCO single (light color Trinity, dark color TransPi). D) Percentage of mapping (reads to transcriptome), BUSCO complete and BUSCO single by individual samples (light color Trinity, dark color TransPi).

### 3.3 TransPi Report

The report generated by TransPi is interactive (i.e., a HTML file is generated) and can be viewed with standard web browsers (Supplementary File 10). The report allows the user to comprehensively assess the data by zooming in in the figures, compare datasets, and see detailed info by selecting specific data points. The report summarizes all major steps performed by the pipeline, including quality filtering, assembly metrics, ORF numbers, annotation and KEGG pathway assignment using iPATH3 (Darzi et al. 2018). TransPi provides the user with multiple files for further downstream analyses of the final reference transcriptome. For example, a file with all Gene Ontologies is created and can be directly used as input for TopGO to perform enrichment analysis (Alexa and Rahnenfuhrer, 2016). All final and key intermediate files, including all plots, are stored in the user selected output directory for manual inspection. Additionally, TransPi will save the execution report generated by Nextflow, in which the user can inspect how their system resources are being used in each process (Example in Supplementary File 9).

### 3.4 Additional TransPi options

The dataset of *Porites pukoensis* (SRR8491966) produced a transcriptome with 567,526 sequences. Despite having a high BUSCO completeness (i.e. 99.4%), the majority of these were duplicates (i.e. 61.2%) (Supplementary File 5). Using the filtration step of TransPi, the number of transcripts was reduced by over 39% (from 567,526 to 343,832). The removed 223,694 transcripts had similarities with the *S. microadriaticum* sequences used for filtering (See Methods 2.6). In the case of the “buscoDist” option, the *Octolasmis warwickii* dataset (SRR10527303) was used and 30 genes that were missing from the TransPi assembly but were present in the other assemblies were found (Figure 6).

**Figure 6.**
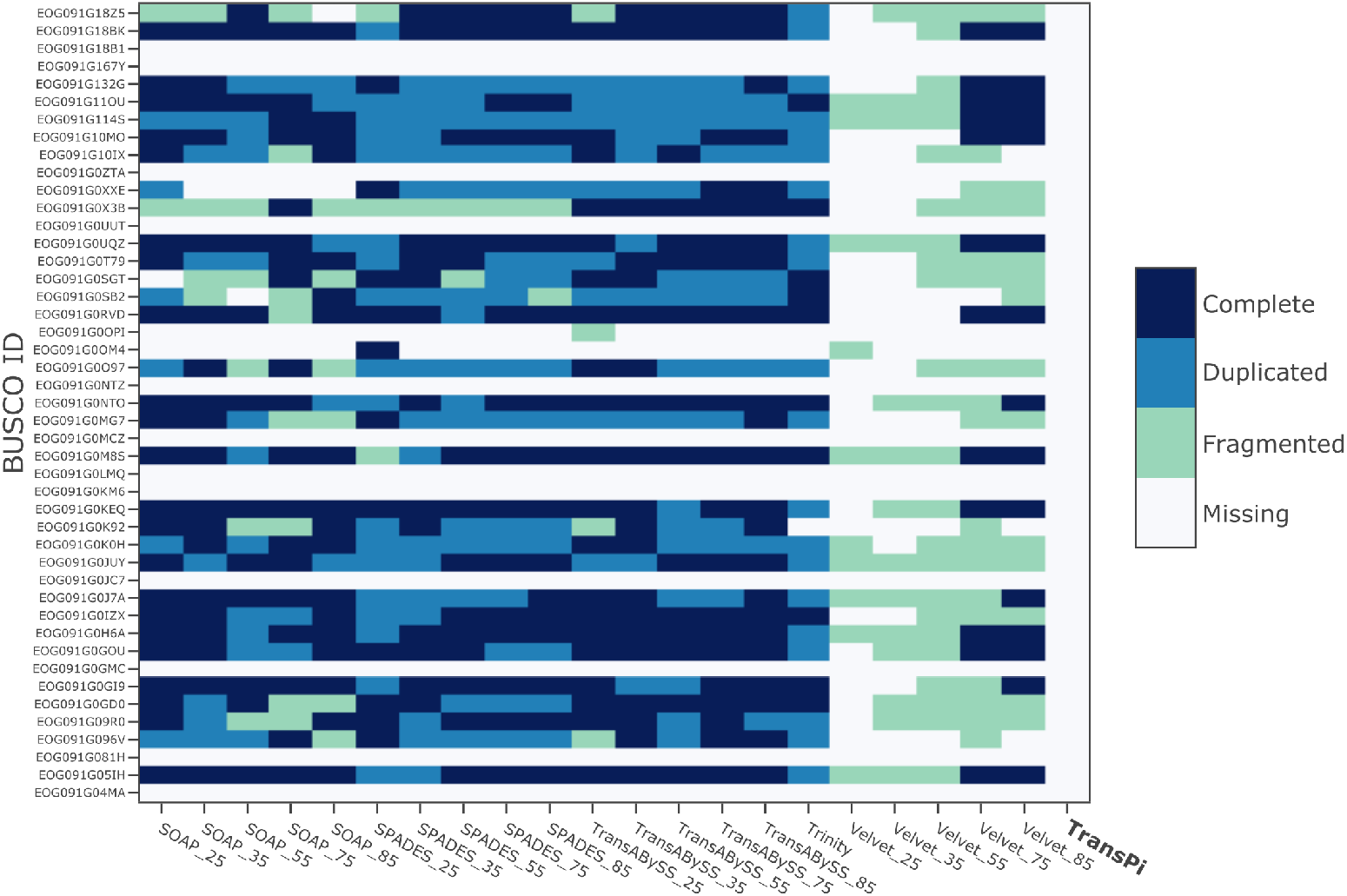
Heatmap of BUSCO gene presence in all assemblers with multiple k-mers that are found missing in TransPi for dataset of Octolasmis warwickii (SRR10527303)

## 4. Discussion

*De novo* transcriptome assemblies are use in several applications such as: differential gene expression (Pita et al., 2018), gene model prediction (Chan et al., 2017), DNA target enrichment bait design (Quek et al., 2020), genome annotation (Testa et al., 2015; Holt and Yandell, 2011), detection of whole-genome duplication (Yang et al., 2019), and phylogenetics (Cheon et al., 2020; Lozano-Fernandez et al., 2019). Even though multiple software are currently available for transcriptome assembly, no single tool is able to generate optimal assemblies given various datasets (Hölzer & Marz 2019). Thus, combining multiple assemblies, generated with various k-mers and software, represents a valuable approach to increase the quality of reference assemblies (Lu et al., 2013). Given the complexity of such analyses, automated workflows are desirable, including the need for standardization, reproducibility, and scalability.

The selection of k-mers is the first step before performing an assembly. The selection of this parameter will modify how the assembly graph is constructed, and it was tested here how k-mer selection can have an effect in the effectiveness of the TransPi pipeline. Tests included different k-mer sizes, combination of k-mers, and different organisms (Table 1). Since TransPi is relying on multiple assemblers and various k-mers, the effect on k-mer selection and their impact on the outcome of the pipeline is minimized. However, k-mer set C consistently resulted in moderately higher BUSCO percentages for single-copy genes and lower duplication levels, respectively. This k-mer set had a wider range of k-mer sizes (from small to long) than the other sets. Small k-mers tend to generate more transcripts but are more prone to misassemblies (Zerbino & Birney, 2008; Gibbons, 2009). On the contrary, longer k-mers produce more contiguous assembly while decreasing transcript numbers (Robertson et al., 2010). Thus, by combining various k-mer sizes (i.e. short and long k-mers), a more comprehensive representation of the transcriptome can be achieved (Peng et al., 2013). The use of different read lengths did not yield significant differences between TransPi and Trinity in all three model organisms (i.e. worm, fly, and mouse) included in this study. Generally, Trinity performed better than TransPi with respect to the ‘complete’ and ‘fragmented’ categories of the metazoa BUSCO genes set. The major advantage of TransPi in the model organisms, however, was the reduction of duplicated BUSCO genes (Figure 2). Furthermore, TransPi had a higher number of non-chimeric transcripts when compare to Trinity alone (Table 2).

**Table 1.** Non-model organisms datasets used in this study.

| Phylum | Class | Order | Species | SRA | # of reads | Length(bp) |
| --- | --- | --- | --- | --- | --- | --- |
| Cnidaria | Anthozoa | Alcyonacea | *Pinnigorgia flava* | ERR3026433 | 30,545,400 | 50 |
| Cnidaria | Anthozoa | Alcyonacea | *Sinularia cruciata* | ERR3026434 | 22,160,908 | 50 |
| Cnidaria | Anthozoa | Alcyonacea | *Tubipora musica* | ERR3026435 | 23,006,724 | 50 |
| Cnidaria | Anthozoa | Helioporacea | *Heliopora coerulea* | ERR3040053 | 29,000,821 | 50 |
| Cnidaria | Anthozoa | Scleractinia | *Acropora palmata* | SRR5569439 | 10,476,071 | 75 |
| Cnidaria | Anthozoa | Scleractinia | *Acropora pulchra* | SRR8601367 | 14,037,157 | 75 |
| Cnidaria | Anthozoa | Scleractinia | *Porites pukoensis* | SRR8491966 | 16,448,725 | 150 |
| Cnidaria | Hydrozoa | Anthoathecata | *Millepora alcicornis* | SRR4294206 | 24,645,545 | 150 |
| Porifera | Homoscleromorpha | Homosclerophorida | *Oscarella pearsei* | SRR1042012 | 11,306,242 | 100 |
| Porifera | Homoscleromorpha | Homosclerophorida | *Corticium candelabrum* | SRR504694 | 18,897,095 | 150 |
| Porifera | Demospongiae | Spongillida | *Ephydatia muelleri* | SRR1041944 | 11,425,188 | 100 |
| Porifera | Demospongiae | Spongillida | *Spongilla lacustris* | SRR1168575 | 5,136,881 | 100 |
| Porifera | Demospongiae | Poecilosclerida | *Mycale phylophylla* | SRR1711043 | 11,408,543 | 100 |
| Porifera | Demospongiae | Haplosclerida | *Haliclona tubifera* | SRR1793376 | 16,356,602 | 100 |
| Porifera | Demospongiae | Dictyoceratida | *Ircinia fasciculata* | SRR7655554 | 13,420,109 | 100 |
| Porifera | Calcarea | Leucosolenida | *Sycon coactum* | SRR504690 | 9,098,097 | 100 |
| Mollusca | Gastropoda | Trochida | *Monodonta labio* | SRR1505119 | 10,388,770 | 100 |
| Mollusca | Bivalvia | Pholadomyoidea | *Lyonsia floridana* | SRR1560310 | 9,919,645 | 100 |
| Mollusca | Bivalvia | Veneroidea | *Mercenaria campechiensis* | SRR1560359 | 11,935,267 | 100 |
| Mollusca | Bivalvia | Trigoniida | *Neotrigonia margaritacea* | SRR1560432 | 11,215,767 | 100 |
| Mollusca | Bivalvia | Veneroidea | *Cardites antiquatus* | SRR1560458 | 11,916,756 | 100 |
| Mollusca | Bivalvia | Veneroidea | *Sphaerium nucleus* | SRR1561723 | 18,539,173 | 100 |
| Mollusca | Bivalvia | Nuculoida | *Ennucula tenuis* | SRR331123 | 14,420,942 | 100 |
| Mollusca | Bivalvia | Ostreoida | *Dimya lima* | SRR3350463 | 5,426,850 | 150 |
| Rotifera | Monogononta | Ploima | *Brachionus plicatilis* | SRR3404576 | 7,403,847 | 150 |
| Arthropoda | Branchiopoda | Diplostraca | *Eoleptestheria cf tacinensis* | SRR5140141 | 5,471,351 | 150 |
| Arthropoda | Remipedia | Nectiopoda | *Godzillignomus frondosus* | SRR8280777 | 14,086,834 | 75 |
| Arthropoda | Arachnida | Solifugae | *Galeodes* sp | SRR8745910 | 6,356,774 | 75 |
| Arthropoda | Hexanauplia | Calanoida | *Neocalanus flemingeri* | SRR5873556 | 4,112,626 | 150 |
| Arthropoda | Hexanauplia | Calanoida | *Calanus finmarchicus* | SRR4113507 | 10,633,606 | 150 |
| Arthropoda | Hexanauplia | Pedunculata | *Octolasmis warwickii* | SRR10527303 | 15,813,391 | 150 |
| Echinodermata | Holothuroidea | Aspidochirotida | *Apostichopus japonicus* | SRR8393254 | 8,289,770 | 150 |
| Echinodermata | Crinoidea | Comatulida | *Florometra* | SRR3097584 | 32,710,859 | 100 |
| Echinodermata | Echinoidea | Echinoida | *Paracentrotus lividus* | ERR1000783 | 6,803,316 | 75 |
| Echinodermata | Echinoidea | Echinoida | *Paracentrotus lividus* | SRR10744002 | 13,583,857 | 75 |
| Xenacoelomorpha | - | Acoela | *Chilidia submaculatum* | SRR3105702 | 6,089,955 | 100 |
| Chaetognatha | Sagittioidea | Aphragmophora | *Krohnitta subtilis* | SRR7754744 | 15,954,007 | 100 |
| Brachiopoda | Rhynchonellata | Rhynchonellida | *Hemithiris psittacea* | SRR1611556 | 9,221,875 | 100 |
| Nemertea | Enopla | Bdellonemertea | *Malacobdella grossa* | SRR1611560 | 8,307,739 | 100 |
| Nemertea | Palaeonemertea | - | *Cephalothrix linearis* | SRR1273789 | 4,869,244 | 75 |
| Phoronida | - | - | *Phoronis psammophila* | SRR1611565 | 12,949,999 | 100 |
| Platyhelminthes | Catenulida | - | *Catenula lemnae* | SRR1796434 | 3,028,636 | 100 |
| Onychophora | Udeonychophora | Euonychophora | *Peripatopsis capensis* | SRR1145776 | 11,638,180 | 100 |
| Onychophora | Udeonychophora | Euonychophora | *Peripatoides novaezealandiae* | SRR8745911 | 5,768,550 | 75 |
| Gastrotricha | - | Macrodasysida | *Macrodasys* sp | SRR1271706 | 3,204,609 | 75 |
| Gastrotricha | - | Chaetonotida | *Lepidodermella squamata* | SRR1273732 | 4,370,938 | 75 |
| Annelida | Polychaeta | Phyllodocida | *Nephtys caeca* | SRR1232685 | 1,576,665 | 75 |
| Gnathostomulida | - | Bursovaginoidea | *Gnathostomula paradoxa* | SRR1271607 | 5,954,962 | 75 |

**Table 2.** Chimera test for model species *C. elegans, D. melanogaster*, and *M. musculus*

| *C. elegans* |  |  |  |  |
| --- | --- | --- | --- | --- |
|  |  | Trinity |  |  |
| Sample | BLASTN hits | # transcripts | % unique | BUSCO v4 - Metazoa DB (n:954) |
| SRR10407355 | 9,538 | 28,219 | 33.80 | C:76.9%[S:65.2%,D:11.7%],F:2.2%,M:20.9% |
| SRR10407357 | 9,014 | 23,526 | 38.32 | C:76.1%[S:65.7%,D:10.4%],F:2.5%,M:21.4% |
| SRR10407359 | 9,310 | 27,734 | 33.57 | C:76.3%[S:65.8%,D:10.5%],F:2.6%,M:21.1% |
|  |  | TransPi |  |  |
| Sample | BLASTN hits | # transcripts | % unique | BUSCO v4 - Metazoa DB (n:954) |
| SRR10407355 | 8,567 | 23,803 | 35.99 | C:75.7%[S:68.9%,D:6.8%],F:2.3%,M:22.0% |
| SRR10407357 | 8,494 | 21,709 | 39.13 | C:75.5%[S:68.7%,D:6.8%],F:2.5%,M:22.0% |
| SRR10407359 | 8,675 | 24,891 | 34.85 | C:75.5%[S:69.6%,D:5.9%],F:2.8%,M:21.7% |
| *D. melanogaster* |  |  |  |  |
|  |  | Trinity |  |  |
| Sample | BLASTN hits | # transcripts | % unique | BUSCO v4 - Metazoa DB (n:954) |
| SRR7716077 | 4,585 | 36,267 | 12.64 | C:97.2%[S:84.6%,D:12.6%],F:1.5%,M:1.3% |
| SRR7716078 | 4,133 | 31,793 | 13.00 | C:93.6%[S:63.6%,D:30.0%],F:1.0%,M:5.4% |
| SRR7716080 | 4,364 | 31,928 | 13.67 | C:90.1%[S:63.4%,D:26.7%],F:3.1%,M:6.8% |
|  |  | TransPi |  |  |
| Sample | BLASTN hits | # transcripts | % unique | BUSCO v4 - Metazoa DB (n:954) |
| SRR7716077 | 4,603 | 29,014 | 15.86 | C:93.9%[S:81.0%,D:12.9%],F:1.3%,M:4.8% |
| SRR7716078 | 4,211 | 25,225 | 16.69 | C:93.8%[S:84.1%,D:9.7%],F:1.5%,M:4.7% |
| SRR7716080 | 4,383 | 25,817 | 16.98 | C:91.4%[S:82.9%,D:8.5%],F:2.2%,M:6.4% |
| *M. musculus* |  |  |  |  |
|  |  | Trinity |  |  |
| Sample | BLASTN hits | # transcripts | % unique | BUSCO v4 - Metazoa DB (n:954) |
| SRR10560364 | 12,360 | 244,523 | 5.05 | C:98.2%[S:53.1%,D:45.1%],F:0.9%,M:0.9% |
| SRR10560365 | 12,807 | 261,000 | 4.91 | C:98.2%[S:48.7%,D:49.5%],F:0.7%,M:1.1% |
| SRR8329326 | 29,885 | 816,077 | 3.66 | C:97.4%[S:36.8%,D:60.6%],F:1.4%,M:1.2% |
|  |  | TransPi |  |  |
| Sample | BLASTN hits | # transcripts | % unique | BUSCO v4 - Metazoa DB (n:954) |
| SRR10560364 | 8,901 | 165,890 | 5.37 | C:97.2%[S:84.6%,D:12.6%],F:1.5%,M:1.3% |
| SRR10560365 | 10,551 | 198,502 | 5.32 | C:96.5%[S:81.0%,D:15.5%],F:1.3%,M:2.2% |
| SRR8329326 | 18,418 | 600,340 | 3.07 | C:94.4%[S:81.7%,D:12.7%],F:2.4%,M:3.2% |

**Table 3.** Statistical tests on non-model organisms

|  | Shapiro-Wilk (p >0.05) |  | Normally distributed | ANOVA | Kruskal-Wallis (p <0.05) | Significant |
| --- | --- | --- | --- | --- | --- | --- |
|  | TransPi | Trinity |  |  |  |  |
| Complete | 5.09E-06 | 1.76E-05 | no | - | 0.6492791 | no |
| Single-copy | 0.0001667 | 0.1081 | no | - | 5.67E-10 | yes |
| Duplicated | 2.51E-07 | 0.008433 | no | - | 9.60E-11 | yes |
| Fragmented | 0.0005825 | 0.002121 | no | - | 0.1375242 | no |
| Missing | 1.81E-08 | 3.61E-09 | no | - | 0.1413003 | no |

It previous studies, it has been shown that using more than 30M read pairs does not significantly improve the quality of the transcriptome assembly (MacManes, 2018; Francis et al., 2013). However, in our tests mixed results were observed when comparing reads quantities and BUSCO scores in each organism respectively (Supplementary File 5). As previously demonstrated, assembly quality and characteristics are data-dependent (Hölzer & Marz 2019). Consequently, to provide a profound conclusion on the effects of reads quantities in *de novo* transcriptome assemblies, a larger number of datasets from a broad range of taxa, in addition to biological replicates for each taxon, are needed. Also, organisms with sources of contamination, for example, of symbiotic origin, prey, parasites or eukaryotic overgrowth in the target tissue, may need higher quantities of reads. In terms of read length, mixed results were observed, and a conclusive comparison of TransPi and Trinity cannot be performed on the sample size used in this study. However, the tests conducted on the three model organisms strongly suggest the usage of longer reads (150bp) should be preferred, because those generally yielded higher-quality transcriptomes with respect to the BUSCO results.

The newly established TransPi pipeline performed significantly better than the Trinity assembler alone on non-model organisms (Figure 3). A high BUSCO completeness with a high number of single-copy BUSCO genes was obtained for the majority of the non-model datasets used here (Figure 3, 4). In the case of the ‘fragmented’ BUSCO genes category, TransPi produced lower scores than Trinity due to the reduction step by EvidentialGene. Since the tool relies on sequence features like coding sequence (CDS) content and length (Gilbert, 2013, 2019), fragmented CDS will be less likely to pass the filtration step. The high number of single-copy BUSCO genes results were statistically significant and are a major difference when comparing with the TransPi results of model organisms. The reduction of transcript duplication is obviously beneficial for studies where the presence of duplicates would bias the interpretation of the results. Another major disadvantage of keeping false isoforms is in phylogenomic analyses. Due to the relative ease of generation and affordability, many phylogenomic studies analyse multi-gene alignments based on transcriptome data instead of full genome data to estimate phylogenies (Lozano-Fernandez et al., 2019; Cheon et al., 2020). By using TransPi, the automation of large-scale phylogenomic approaches, focusing on thousands of proteins from many taxa, can be attained with ease in a scalable and reproducible way.

As expected, the final number of transcripts was consistently lower for TransPi given the reduction performed by EvidentialGene (Figure 5). In some cases (*Malacobdella grossa*) the reduction of transcripts was over 50% (Figure 5; Supplementary File 5). This explains why the mapping percentages for TransPi were also lower than for Trinity. However, having reduced mapping rates (i.e. TransPi) did not affect the content of BUSCO genes in the transcriptomes. For example, in the *Malacobdella grossa* assembly, TransPi mapping was 65.41% versus 82.86% for Trinity (Figure 5; Supplementary File 5), but the difference of complete BUSCO genes was only 0.30% (TransPi=94.0%, Trinity=94.30%) and 62% (TransPi=83.0%, Trinity=20.4%) for single copy BUSCOs, respectively. Thus, the reduction in mapping percentages is due to the reduction of redundant transcripts (including allelic variants) rather than missing information from the assemblies. This pattern was observed throughout the non-model organisms analysed here (Figure 5; Supplementary File 5). In general, when the mapping percentage of TransPi was over 65%, satisfactory BUSCO content in the transcriptomes (i.e. high BUSCO presence and in single copies) was observed. However, there were some cases where both TransPi and Trinity produced equally low BUSCO scores, even though a relative high mapping percentage was obtained (Figure 5; Supplementary File 7). This was the case for a *Catenula lemnae* dataset (SRR1796434), where read mapping percentage was relatively high (74.69% and 89.37% for TransPi and Trinity, respectively), while the BUSCO gene con-tent (complete and single) were <53% (Figure 5; Supplementary File 7). In such cases, the assemblies may not be optimal and likely do not represent the complete transcriptome of the organism. (Figure 5; Supplementary File 7).

For the missing BUSCO category, TransPi produced assemblies with slightly higher values in comparison to the other assemblers. When a BUSCO gene is missing in TransPi (i.e. removed by the EvidentialGene step), in some cases, these genes are found in the other individual assemblies (Figure 6). EvidentialGene aims to keep the most valid biological transcript, discards the likely not valid (based on specific measures), and decreases the redundancy of the multiple assemblers to obtain a non-redundant consensus transcriptome assembly (Gilbert, 2019). However, by doing so, some genes can be categorized as redundants, presumably, because better candidates were selected. To get more insight into cases like the one above, the TransPi option “buscoDist” was used with the *Octolasmis warwickii* dataset. Comparing the missing genes of all generated assemblies and plotting the distribution of the BUSCO genes showed that TransPi had more missing genes that were categorized in other assemblers as to be present (Figure 6). However, a considerable amount of these genes were classified as duplicates by BUSCO. Since the BUSCO scores are indicators of the transcriptome completeness, correcting them will provide a more realistic estimation on the transcriptome quality of a given taxon. This TransPi option offers the user insight into the BUSCO gene content and transcripts reduction by EvidentialGene to help better assess the quality of the assemblies.

In certain cases, significant numbers of BUSCO genes were not retrieved by TransPi, Trinity or any of the assemblers. Although this could be related to assembler performance, other factors have been shown to alter transcriptome quality (e.g. RNA degradation, library preparation, sequencing depth, etc.) (Romero et al., 2014; Sultan 2014). In the non-model organisms, four of the datasets yielded BUSCO complete percentages <50% in TransPi and Trinity (Supplementary File 5). Three of these datasets (i.e. *Mercenaria campechiensis* (SRR1560359), *Sphaerium nucleus* (SRR1561723), *Cardites antiquatus* (SRR1560458)) stem from the same Project and the same taxonomic group, the molluscan class Bivalvia. Extraction of nucleic acids in molluscs is known to be hampered by the presence of mucopolysaccharides and polyphenolic proteins, which can inhibit PCR and lead to biases in RNA preservation and/or the extraction quantities and/or qualities (Rzepecki et al., 1991; Gayral et al., 2011; Knutson et al., 2020). Nevertheless, BUSCO genes that were retrieved exhibited low rates of duplication, highlighting how the incorporation of EvidentialGene into TransPi can also decrease redundancy in cases of low transcriptome completeness. For the fourth dataset (*Nephtys caeca* (SRR1232685), a polychaete annelid), only a small number of 1.5M read pairs were deposited in INSDC databases (i.e. NCBI’s Genbank), which helps to explain the poor results (Supplementary File 5). Deeper sequencing of these particular specimens may well lead to an improved transcriptome. This also might indicate that quantity of reads rather than the quality of input material was the limiting factor for the generation of a complete transcriptome.

TransPi also addresses putative contamination issues that might affect a transcriptome by providing an additional option that performs filtering of “contaminants”. Datasets from organisms like corals can represent a challenge during transcriptome assembly and down-stream analyses due to their endosymbiotic zooxanthellae (Shinzato et al., 2014). Thus, a filtration step is usually performed to remove sequences that do not belong to the target (host) transcriptome (Veglia et al., 2018). The filtration step of TransPi is a useful step in the cases of known contamination sources. For example, in the dataset of the coral *Porites pukoensis* both programs, TransPi and Trinity obtained high BUSCO completeness percentages. However, despite the reduction with EvidentialGene, single-copy BUSCO percentages were low and the percentage of duplicated BUSCO genes was high in both, TransPi and Trinity. Given the shown strong efficiency of TransPi to remove redundancy, the presence of many duplicates in this dataset may indicate the presence of algae (symbionts) transcripts and/or contamination. Also, it has been previously reported that other eukaryotes, particularly fungi, are commonly found in *Porites pukoensis* (Li et al., 2014). This could potentially bias the outcomes and can strongly affect downstream analyses. Thus, using a contaminant filtration step, as performed by TransPi, is beneficial to generate a cleaner and accurate transcriptome assembly and provide the user with a host only transcriptome to be further analyzed.

In summary, TransPi offers researchers working with non-model organisms the opportunity of a comprehensive *de novo* transcriptome analysis, requiring minimum user input but without losing the ability of a thorough analysis. The non-redundant assembly generated by TransPi can be used directly in several downstream analyses including differential expression, gene modelling for genome annotations, bait design and phylogenetics. Another key advantage of using TransPi is that it offers reproducibility of the results with ease, where entire experiments can be repeated with defined versions of all programs included in the workflow. It provides a user-friendly environment, easy deployment, and scalability by employing Nextflow. TransPi also has other additional features to help gain extra insight into the assemblies. We anticipate that TransPi will be a valuable tool for the generation of comprehensive *de novo* non-redundant transcriptome assemblies for non-model organisms.

## Data accessibility

Data are available online: https://github.com/PalMuc/TransPi, 10.5281/zenodo.5060055

## Supplementary material

Script and codes are available online: https://github.com/PalMuc/TransPi, 10.5281/zenodo.5060055

## Acknowledgements

Version 3 of this preprint has been peer-reviewed and recommended by Peer Community In Genomics *Peer Community In Genomics* (https://doi.org/10.24072/pci.genomics.100009). RERV, ME and GW acknowledge funding from the European Union’s Horizon 2020 research and innovation programme under the Marie Skłodowska-Curie grant agreement No 764840 (ITN IGNITE). CGE acknowledges the Advanced Human Capital Program of the National Commission for Scientific and Technological Research (CONICYT) for the Becas-Chile Scholarship awarded to study at LMU. CGE and ME acknowledges funding by Lehre@LMU (project number: W19_F1; Studi_forscht@GEO). GW acknowledges funding through the LMU Munich’s Institutional Strategy LMUexcellent within the framework of the German Excellence Initiative. The authors gratefully acknowledge the Leibniz Supercomputing Centre (LRZ) as a partner of ITN IGNITE for providing computing time and support on its Linux-Cluster and Compute Cloud system.

