## Supplementary Table for "TransPi – a comprehensive TRanscriptome ANalysiS PIpeline for *de novo* transcriptome assembly"

Supplementary Table 1. Model organisms used for kmer, reads length and reads quantity tests.

| Species | # of Reads | SRA | Length(bp) | Species | # of Reads | SRA | Length(bp) | Species | # of Reads | SRA | Length(bp) |
|---|---|---|---|---|---|---|---|---|---|---|---|
| *D. melanogaster* | 15,648,174 | SRR6743334 | 50 | *C. elegans* | 24,017,399 | SRR4181026 | 50 | *M. musculus* | 33,540,466 | SRR10342165 | 50 |
| *D. melanogaster* | 15,181,806 | SRR6743335 | 50 | *C. elegans* | 24,409,279 | SRR4181032 | 50 | *M. musculus* | 43,541,512 | SRR10342166 | 50 |
| *D. melanogaster* | 9,857,308 | SRR6743336 | 50 | *C. elegans* | 24,694,386 | SRR4181033 | 50 | *M. musculus* | 43,440,008 | SRR10342167 | 50 |
| *D. melanogaster* | 15,940,368 | SRR6743337 | 50 | *C. elegans* | 26,159,433 | SRR4181034 | 50 | *M. musculus* | 33,595,655 | SRR9211836 | 50 |
| *D. melanogaster* | 18,874,930 | SRR6743338 | 50 | *C. elegans* | 26,417,070 | SRR4181036 | 50 | *M. musculus* | 33,174,108 | SRR9211837 | 50 |
| *D. melanogaster* | 21,169,448 | SRR6743339 | 50 | *C. elegans* | 24,711,402 | SRR4181037 | 50 | *M. musculus* | 35,030,665 | SRR9211838 | 50 |
| *D. melanogaster* | 16,177,860 | SRR6743340 | 50 | *C. elegans* | 40,302,838 | SRR1585277 | 75 | *M. musculus* | 34,877,241 | SRR9211839 | 50 |
| *D. melanogaster* | 12,436,324 | SRR6743341 | 50 | *C. elegans* | 50,516,835 | SRR1585278 | 75 | *M. musculus* | 33,700,156 | SRR10067458 | 75 |
| *D. melanogaster* | 26,395,792 | SRR6743342 | 50 | *C. elegans* | 41,947,175 | SRR8257107 | 75 | *M. musculus* | 41,236,457 | SRR10067459 | 75 |
| *D. melanogaster* | 87,423,452 | SRR8559044 | 75 | *C. elegans* | 44,969,393 | SRR8257108 | 75 | *M. musculus* | 35,598,598 | SRR10067460 | 75 |
| *D. melanogaster* | 85,714,154 | SRR8559045 | 75 | *C. elegans* | 45,605,396 | SRR8257109 | 75 | *M. musculus* | 41,745,958 | SRR10067461 | 75 |
| *D. melanogaster* | 88,252,694 | SRR8559046 | 75 | *C. elegans* | 49,743,412 | SRR2142254 | 100 | *M. musculus* | 45,329,544 | SRR10067462 | 75 |
| *D. melanogaster* | 82,110,608 | SRR8559047 | 75 | *C. elegans* | 38,836,876 | SRR2142255 | 100 | *M. musculus* | 44,469,310 | SRR10067463 | 75 |
| *D. melanogaster* | 102,413,880 | SRR8559048 | 75 | *C. elegans* | 11,166,310 | SRR2015313 | 100 | *M. musculus* | 182,180,123 | SRR1017092 | 100 |
| *D. melanogaster* | 98,529,578 | SRR8559049 | 75 | *C. elegans* | 11,626,153 | SRR2015311 | 100 | *M. musculus* | 107,280,600 | SRR1017093 | 100 |
| *D. melanogaster* | 30,515,068 | SRR3018852 | 100 | *C. elegans* | 12,567,755 | SRR2012785 | 100 | *M. musculus* | 127,017,374 | SRR1017094 | 100 |
| *D. melanogaster* | 24,284,630 | SRR3018855 | 100 | *C. elegans* | 14,372,593 | SRR2012781 | 100 | *M. musculus* | 102,909,165 | SRR5171077 | 100 |
| *D. melanogaster* | 24,773,404 | SRR3018862 | 100 | *C. elegans* | 20,345,696 | SRR10407359 | 150 | *M. musculus* | 65,372,078 | SRR5171085 | 100 |
| *D. melanogaster* | 30,140,704 | SRR3019045 | 100 | *C. elegans* | 19,948,493 | SRR10407358 | 150 | *M. musculus* | 27,016,597 | SRR5171095 | 100 |
| *D. melanogaster* | 25,828,680 | SRR3019179 | 100 | *C. elegans* | 21,276,093 | SRR10407357 | 150 | *M. musculus* | 21,173,335 | SRR10560364 | 150 |
| *D. melanogaster* | 19,049,236 | SRR3019184 | 100 | *C. elegans* | 23,993,704 | SRR10407356 | 150 | *M. musculus* | 22,731,971 | SRR10560365 | 150 |
| *D. melanogaster* | 8,276,228 | SRR10407358 | 150 | *C. elegans* | 25,110,365 | SRR10407355 | 150 | *M. musculus* | 68,712,046 | SRR8329325 | 150 |
| *D. melanogaster* | 47,552,414 | SRR7716077 | 150 | *C. elegans* | 22,967,010 | SRR10407354 | 150 | *M. musculus* | 59,012,049 | SRR8329326 | 150 |
| *D. melanogaster* | 45,807,064 | SRR7716078 | 150 | | | | | *M. musculus* | 21,930,527 | SRR7079069 | 150 |
| *D. melanogaster* | 29,221,598 | SRR7716079 | 150 | | | | | | | | |
| *D. melanogaster* | 46,936,912 | SRR7716080 | 150 | | | | | | | | |

Supplementary Table 2. Kmer test results and statistical tests.

| | | | | 50bp (A) 25,29,33 | | | | | 50bp (B) 25,31,35 | | | | | 50bp (C) 25,33,37 | | | | |
|---|---|---|---|---|---|---|---|---|---|---|---|---|---|---|---|---|---|---|
| Nº | Taxa | Total pe-reads | SRA | Complete % | Single-copy % | Duplicated % | Fragmented % | Missing % | Complete % | Single-copy % | Duplicated % | Fragmented % | Missing % | Complete % | Single-copy % | Duplicated % | Fragmented % | Missing % |
| 1 | *M. musculus* | 33,540,466 | SRR10342165 | 96.9 | 85.9 | 11 | 1.2 | 1.9 | 96.8 | 86.2 | 10.6 | 1.5 | 1.7 | 97.5 | 86.8 | 10.7 | 1 | 1.5 |
| 2 | *M. musculus* | 43,541,512 | SRR10342166 | 96.4 | 85.5 | 10.9 | 1.6 | 2 | 96.9 | 86.1 | 10.8 | 1.6 | 1.5 | 96.5 | 84.9 | 11.6 | 1.8 | 1.7 |
| 3 | *M. musculus* | 43,440,008 | SRR10342167 | 97 | 86 | 11 | 0.9 | 2.1 | 96.7 | 86 | 10.7 | 1.3 | 2 | 97 | 85.8 | 11.2 | 1 | 2 |
| 4 | *M. musculus* | 33,595,655 | SRR9211836 | 95.8 | 84.9 | 10.9 | 1.7 | 2.5 | 95.9 | 85.9 | 10 | 1.7 | 2.4 | 95.8 | 85 | 10.8 | 1.8 | 2.4 |
| 5 | *M. musculus* | 33174108 | SRR9211837 | 95 | 83.1 | 11.9 | 2.1 | 2.9 | 95.2 | 84.3 | 10.9 | 1.9 | 2.9 | 95 | 83.1 | 11.9 | 2 | 3 |
| 6 | *M. musculus* | 35,030,665 | SRR9211838 | 95.5 | 85.5 | 10 | 1.7 | 2.8 | 95.7 | 84.4 | 11.3 | 1.7 | 2.6 | 95.6 | 85.4 | 10.2 | 1.6 | 2.8 |
| 7 | *M. musculus* | 34,877,241 | SRR9211839 | 95.6 | 84.5 | 11.1 | 1.4 | 3 | 95.4 | 83.7 | 11.7 | 1.7 | 2.9 | 95.6 | 83.9 | 11.7 | 1.5 | 2.9 |
| 1 | *C. elegans* | 24,017,399 | SRR4181026 | 85.5 | 77.9 | 7.6 | 1.7 | 12.8 | 85.4 | 78.1 | 7.3 | 1.8 | 12.8 | 85.4 | 78.2 | 7.2 | 1.8 | 12.8 |
| 2 | *C. elegans* | 24,409,279 | SRR4181032 | 85.3 | 77.6 | 7.7 | 1.7 | 13 | 85 | 77.8 | 7.2 | 1.8 | 13.2 | 85.2 | 77.6 | 7.6 | 1.8 | 13 |
| 3 | *C. elegans* | 24,694,386 | SRR4181033 | 85.3 | 78.4 | 6.9 | 1.8 | 12.9 | 85.5 | 78.9 | 6.6 | 1.6 | 12.9 | 85.2 | 79 | 6.2 | 1.8 | 13 |
| 4 | *C. elegans* | 26,159,433 | SRR4181034 | 85.7 | 78.5 | 7.2 | 1.5 | 12.8 | 85.7 | 78.6 | 7.1 | 1.6 | 12.7 | 85.7 | 78.8 | 6.9 | 1.6 | 12.7 |
| 5 | *C. elegans* | 26,417,070 | SRR4181036 | 85.4 | 79.6 | 5.8 | 1.8 | 12.8 | 85.5 | 79.1 | 6.4 | 1.6 | 12.9 | 85.4 | 79.3 | 6.1 | 1.8 | 12.8 |
| 6 | *C. elegans* | 24,711,402 | SRR4181037 | 85.2 | 78.2 | 7 | 2.1 | 12.7 | 85.1 | 77.9 | 7.2 | 2.1 | 12.8 | 85.1 | 78.7 | 6.4 | 2.1 | 12.8 |
| 1 | *D. melanogaste* | 15,648,174 | SRR6743334 | 44.7 | 40.6 | 4.1 | 7.6 | 47.7 | 44.1 | 39.8 | 4.3 | 8.6 | 47.3 | 44.5 | 40.4 | 4.1 | 8.2 | 47.3 |
| 2 | *D. melanogaste* | 15,181,806 | SRR6743335 | 44.6 | 41.5 | 3.1 | 7.4 | 48 | 43.7 | 40.3 | 3.4 | 7.6 | 48.7 | 43.7 | 40.7 | 3 | 7.5 | 48.8 |
| 3 | *D. melanogaste* | 9,857,308 | SRR6743336 | 37 | 33.6 | 3.4 | 8.1 | 54.9 | 36.9 | 33.7 | 3.2 | 7.8 | 55.3 | 36.8 | 33.8 | 3 | 8 | 55.2 |
| 4 | *D. melanogaste* | 15,940,368 | SRR6743337 | 39.8 | 36.5 | 3.3 | 7 | 53.2 | 40 | 36.8 | 3.2 | 6.9 | 53.1 | 40.1 | 37.2 | 2.9 | 7.3 | 52.6 |
| 5 | *D. melanogaste* | 18,874,930 | SRR6743338 | 48.5 | 45.5 | 3 | 7.5 | 44 | 47.5 | 44.6 | 2.9 | 8.1 | 44.4 | 47.9 | 44.8 | 3.1 | 8.3 | 43.8 |
| 6 | *D. melanogaste* | 21,169,448 | SRR6743339 | 35.1 | 32.2 | 2.9 | 7.8 | 57.1 | 34.2 | 31.3 | 2.9 | 8.7 | 57.1 | 34.2 | 31.5 | 2.7 | 8.4 | 57.4 |
| 7 | *D. melanogaste* | 16,177,860 | SRR6743340 | 41.6 | 39.1 | 2.5 | 7.2 | 51.2 | 41.1 | 38.2 | 2.9 | 7.3 | 51.6 | 41.7 | 39 | 2.7 | 7.5 | 50.8 |
| 8 | *D. melanogaste* | 12,436,324 | SRR6743341 | 40.1 | 36.8 | 3.3 | 7.2 | 52.7 | 39.9 | 36.2 | 3.7 | 7.8 | 52.3 | 39.7 | 36.6 | 3.1 | 8 | 52.3 |
| 9 | *D. melanogaste* | 26,395,792 | SRR6743342 | 66.4 | 60.6 | 5.8 | 6.9 | 26.7 | 66.1 | 60.2 | 5.9 | 7.4 | 26.5 | 66.4 | 60.3 | 6.1 | 6.9 | 26.7 |

| | | | | 75bp (A) 25,29,33,41 | | | | | 75bp (B) 25,31,35,43 | | | | | 75bp (C) 25,33,37,45 | | | | |
|---|---|---|---|---|---|---|---|---|---|---|---|---|---|---|---|---|---|---|
| Nº | Taxa | Total pe-reads | SRA | Complete % | Single-copy % | Duplicated % | Fragmented % | Missing % | Complete % | Single-copy % | Duplicated % | Fragmented % | Missing % | Complete % | Single-copy % | Duplicated % | Fragmented % | Missing % |
| 1 | *M. musculus* | 33,700,156 | SRR10067458 | 98 | 84.3 | 13.7 | 1.2 | 0.8 | 97.7 | 82.8 | 14.9 | 1.3 | 1 | 98.1 | 84.2 | 13.9 | 1.1 | 0.8 |
| 2 | *M. musculus* | 41,236,457 | SRR10067459 | 98 | 82.2 | 15.8 | 1 | 1 | 98.1 | 83 | 15.1 | 0.8 | 1.1 | 98.2 | 82.6 | 15.6 | 0.8 | 1 |
| 3 | *M. musculus* | 35,598,598 | SRR10067460 | 98 | 82.6 | 15.4 | 0.9 | 1.1 | 97.9 | 83.6 | 14.3 | 1.1 | 1 | 97.8 | 82.7 | 15.1 | 0.9 | 1.3 |
| 4 | *M. musculus* | 41,745,958 | SRR10067461 | 93.9 | 82.2 | 11.7 | 3 | 3.1 | 94 | 82.1 | 11.9 | 2.9 | 3.1 | 94.1 | 82.6 | 11.5 | 2.9 | 3 |
| 5 | *M. musculus* | 45,329,544 | SRR10067462 | 96.1 | 82.7 | 13.4 | 1.5 | 2.4 | 95.8 | 82.5 | 13.3 | 1.7 | 2.5 | 95.7 | 82 | 13.7 | 1.9 | 2.4 |
| 6 | *M. musculus* | 44,469,310 | SRR10067463 | 94.5 | 82.6 | 11.9 | 2.4 | 3.1 | 94.5 | 82.6 | 11.9 | 2.5 | 3 | 94.4 | 82.5 | 11.9 | 2.5 | 3.1 |
| 1 | *C. elegans* | 40,302,838 | SRR1585277 | 86.0 | 74.0 | 12.0 | 1.8 | 12.2 | 86.2 | 74.9 | 11.3 | 1.8 | 12.0 | 86.1 | 74.3 | 11.8 | 1.9 | 12.0 |
| 2 | *C. elegans* | 50,516,835 | SRR1585278 | 86.2 | 72.0 | 14.2 | 1.7 | 12.1 | 86.2 | 71.5 | 14.7 | 1.4 | 12.4 | 85.6 | 71.9 | 13.7 | 1.8 | 12.6 |
| 3 | *C. elegans* | 41,947,175 | SRR8257107 | 83.9 | 78.4 | 5.5 | 2.4 | 13.7 | 84.2 | 78.1 | 6.1 | 2.2 | 13.6 | 84.0 | 78.0 | 6.0 | 2.4 | 13.6 |
| 4 | *C. elegans* | 44,969,393 | SRR8257108 | 83.8 | 79.2 | 4.6 | 1.9 | 14.3 | 83.9 | 79.4 | 4.5 | 1.9 | 14.2 | 84.1 | 79.7 | 4.4 | 2.0 | 13.9 |
| 5 | *C. elegans* | 45,605,396 | SRR8257109 | 84.1 | 78.8 | 5.3 | 1.9 | 14.0 | 84.4 | 79.9 | 4.5 | 2.0 | 13.6 | 84.3 | 79.6 | 4.7 | 1.9 | 13.8 |
| 1 | *D. melanogaste* | 87,423,452 | SRR8559044 | 97.1 | 77.5 | 19.6 | 1.1 | 1.8 | 97.5 | 77.5 | 20 | 0.8 | 1.7 | 96.8 | 79.4 | 17.4 | 1.4 | 1.8 |
| 2 | *D. melanogaste* | 85,714,154 | SRR8559045 | 97.6 | 78.8 | 18.8 | 0.7 | 1.7 | 97.3 | 80.2 | 17.1 | 0.6 | 2.1 | 97.2 | 77.8 | 19.4 | 0.8 | 2 |
| 3 | *D. melanogaste* | 88,252,694 | SRR8559046 | 96.6 | 75.6 | 21 | 1.2 | 2.2 | 96.7 | 77.5 | 19.2 | 0.9 | 2.4 | 96.4 | 78.4 | 18 | 1.1 | 2.5 |
| 4 | *D. melanogaste* | 82,110,608 | SRR8559047 | 97.4 | 78.7 | 18.7 | 0.6 | 2 | 97.7 | 78.6 | 19.1 | 0.5 | 1.8 | 97.3 | 78.5 | 18.8 | 0.8 | 1.9 |
| 5 | *D. melanogaste* | 102,413,880 | SRR8559048 | 96.8 | 74.1 | 22.7 | 1.1 | 2.1 | 96.5 | 73.8 | 22.7 | 1.1 | 2.4 | 96.6 | 75.9 | 20.7 | 1.4 | 2 |
| 6 | *D. melanogaste* | 98,529,578 | SRR8559049 | 97.6 | 76.8 | 20.8 | 0.8 | 1.6 | 97.8 | 78.3 | 19.5 | 0.4 | 1.8 | 97.2 | 78.3 | 18.9 | 0.8 | 2 |

| | | | | 100bp (A) 25,35,47,61 | | | | | 100bp (B) 25,37,55,63 | | | | | 100bp (C) 25,41,57,67 | | | | |
|---|---|---|---|---|---|---|---|---|---|---|---|---|---|---|---|---|---|---|
| Nº | Taxa | Total pe-reads | SRA | Complete % | Single-copy % | Duplicated % | Fragmented % | Missing % | Complete % | Single-copy % | Duplicated % | Fragmented % | Missing % | Complete % | Single-copy % | Duplicated % | Fragmented % | Missing % |
| 1 | *M. musculus* | 182,180,123 | SRR1017092 | 96.3 | 76.3 | 20 | 2.1 | 1.6 | 96.9 | 76 | 20.9 | 1.6 | 1.5 | 96.7 | 75.1 | 21.6 | 1.8 | 1.5 |
| 2 | *M. musculus* | 107,280,600 | SRR1017093 | 97 | 75.5 | 21.5 | 1.7 | 1.3 | 96.7 | 76.7 | 20 | 2.6 | 0.7 | 97.3 | 75.2 | 22.1 | 1.8 | 0.9 |
| 3 | *M. musculus* | 127,017,374 | SRR1017094 | 97.3 | 77.2 | 20.1 | 1.3 | 1.4 | 96.3 | 76.2 | 20.1 | 1.9 | 1.8 | 97.4 | 77 | 20.4 | 1.5 | 1.1 |
| 4 | *M. musculus* | 29,892,315 | SRR5171077 | 97.3 | 79.9 | 17.4 | 2.2 | 0.5 | 96.9 | 80.8 | 16.1 | 2.9 | 0.2 | 42.7 | 40.3 | 2.4 | 19.7 | 37.6 |
| 5 | *M. musculus* | 30,611,590 | SRR5171085 | 97.3 | 82.2 | 15.1 | 2.5 | 0.2 | 97.5 | 82.4 | 15.1 | 2 | 0.5 | 97.3 | 79.9 | 17.4 | 2.1 | 0.6 |
| 6 | *M. musculus* | 37,729,075 | SRR5171095 | 53.9 | 50.5 | 3.4 | 19.9 | 26.2 | 54.1 | 50.6 | 3.5 | 19.5 | 26.4 | 53.8 | 50.9 | 2.9 | 20 | 26.2 |
| 1 | *C. elegans* | 49,743,412 | SRR2142254 | 82.3 | 76.4 | 5.9 | 3.3 | 14.4 | 82.4 | 77 | 5.4 | 3.3 | 14.3 | 82.4 | 76.7 | 5.7 | 3.2 | 14.4 |
| 2 | *C. elegans* | 38,836,876 | SRR2142255 | 82.3 | 78.6 | 3.7 | 2.7 | 15 | 82.3 | 78.8 | 3.5 | 2.8 | 14.9 | 82.2 | 77.9 | 4.3 | 2.8 | 15 |
| 3 | *C. elegans* | 11,166,310 | SRR2015313 | 67.1 | 63.9 | 3.2 | 13.1 | 19.8 | 66.9 | 63.3 | 3.6 | 13.2 | 19.9 | 66.6 | 63.4 | 3.2 | 13.9 | 19.5 |
| 4 | *C. elegans* | 11,626,153 | SRR2015311 | 80.4 | 76.6 | 3.8 | 4.6 | 15 | 80.5 | 76.3 | 4.2 | 4.5 | 15 | 80.5 | 76.8 | 3.7 | 4.4 | 15.1 |
| 5 | *C. elegans* | 12,567,755 | SRR2012785 | 85.3 | 73 | 12.3 | 2.6 | 12.1 | 85.1 | 72.3 | 12.8 | 2.6 | 12.3 | 85 | 71.4 | 13.6 | 2.7 | 12.3 |
| 6 | *C. elegans* | 14,372,593 | SRR2012781 | 69.6 | 62.3 | 7.3 | 11.3 | 19.1 | 69.5 | 63.2 | 6.3 | 11.2 | 19.3 | 69.7 | 63.1 | 6.6 | 11.2 | 19.1 |
| 1 | *D. melanogaste* | 30,515,068 | SRR3018852 | 95.4 | 80.2 | 15.2 | 1.3 | 3.3 | 95.4 | 78.3 | 17.1 | 1.6 | 3 | 95.7 | 78.9 | 16.8 | 1.2 | 3.1 |
| 2 | *D. melanogaste* | 24,284,630 | SRR3018855 | 95.8 | 78.6 | 17.2 | 0.8 | 3.4 | 95.6 | 78.4 | 17.2 | 1 | 3.4 | 95.9 | 79 | 16.9 | 0.8 | 3.3 |
| 3 | *D. melanogaste* | 24,773,404 | SRR3018862 | 95 | 80.9 | 14.1 | 1.7 | 3.3 | 95.3 | 81 | 14.3 | 1.7 | 3 | 95.6 | 82.3 | 13.3 | 1.6 | 2.8 |
| 4 | *D. melanogaste* | 30,140,704 | SRR3019045 | 95.6 | 78.9 | 16.7 | 1.3 | 3.1 | 95.7 | 79.2 | 16.5 | 1.5 | 2.8 | 95.3 | 79.8 | 15.5 | 1.5 | 3.2 |
| 5 | *D. melanogaste* | 25,828,680 | SRR3019179 | 94.6 | 80 | 14.6 | 1.6 | 3.8 | 94.9 | 81.2 | 13.7 | 1.4 | 3.7 | 95 | 80 | 15 | 1.5 | 3.5 |
| 6 | *D. melanogaste* | 19,049,236 | SRR3019184 | 95.2 | 80.2 | 15 | 1.4 | 3.4 | 95 | 81.5 | 13.5 | 1.3 | 3.7 | 95.4 | 81.7 | 13.7 | 1 | 3.6 |

| | | | | 150bp (A) 25,31,41,51,65 | | | | | 150bp (B) 25,33,43,53,71 | | | | | 150bp (C) 25,35,55,75,85 | | | | |
|---|---|---|---|---|---|---|---|---|---|---|---|---|---|---|---|---|---|---|
| Nº | Taxa | Total pe-reads | SRA | Complete % | Single-copy % | Duplicated % | Fragmented % | Missing % | Complete % | Single-copy % | Duplicated % | Fragmented % | Missing % | Complete % | Single-copy % | Duplicated % | Fragmented % | Missing % |
| 1 | *M. musculus* | 21,173,335 | SRR10560364 | 97.1 | 79 | 18.1 | 1.4 | 1.5 | 97.2 | 81.5 | 15.7 | 1.3 | 1.5 | 97.3 | 81.1 | 16.2 | 1.3 | 1.4 |
| 2 | *M. musculus* | 22,731,971 | SRR10560365 | 96.7 | 78.6 | 18.1 | 1.7 | 1.6 | 97.1 | 79.3 | 17.8 | 1.2 | 1.7 | 97.1 | 78.8 | 18.3 | 1.1 | 1.8 |
| 3 | *M. musculus* | 68,712,046 | SRR8329325 | 94.6 | 82.6 | 12 | 2.7 | 2.7 | 94.8 | 83.1 | 11.7 | 3.1 | 2.1 | 93.7 | 83.2 | 10.5 | 3.5 | 2.8 |
| 4 | *M. musculus* | 59,012,049 | SRR8329326 | 95.9 | 81.5 | 14.4 | 2.2 | 1.9 | 95.6 | 81.9 | 13.7 | 2.6 | 1.8 | 95.9 | 81.4 | 14.5 | 2.1 | 2 |
| 5 | *M. musculus* | 21,930,527 | SRR7079069 | 95.7 | 74.5 | 21.2 | 2.4 | 1.9 | 95.5 | 75.2 | 20.3 | 2.7 | 1.8 | 96.1 | 75.8 | 20.3 | 2.5 | 1.4 |
| 1 | *C. elegans* | 20,345,696 | SRR10407359 | 85.9 | 76.2 | 9.7 | 1.7 | 12.4 | 85.7 | 75.3 | 10.4 | 1.8 | 12.5 | 85.7 | 76 | 9.7 | 1.7 | 12.6 |
| 2 | *C. elegans* | 19,948,493 | SRR10407358 | 86.4 | 75.1 | 11.3 | 1.3 | 12.3 | 85.8 | 74.8 | 11 | 1.4 | 12.8 | 86.3 | 75.8 | 10.5 | 1.3 | 12.4 |
| 3 | *C. elegans* | 21,276,093 | SRR10407357 | 85.4 | 74.4 | 11 | 2 | 12.6 | 85 | 74 | 11 | 2 | 13 | 85.6 | 73.6 | 12 | 2 | 12.4 |
| 4 | *C. elegans* | 23,993,704 | SRR10407356 | 85.7 | 76 | 9.7 | 2 | 12.3 | 85.3 | 75.5 | 9.8 | 2.2 | 12.5 | 85.7 | 76.2 | 9.5 | 1.8 | 12.5 |
| 5 | *C. elegans* | 25,110,365 | SRR10407355 | 85.7 | 75.8 | 9.9 | 2.1 | 12.2 | 85.3 | 74.8 | 10.5 | 1.9 | 12.8 | 85.9 | 75.3 | 10.6 | 2 | 12.1 |
| 6 | *C. elegans* | 22,967,010 | SRR10407354 | 84.7 | 74.7 | 10 | 2.7 | 12.6 | 85 | 75.2 | 9.8 | 2.5 | 12.5 | 84.9 | 74.8 | 10.1 | 2.4 | 12.7 |
| 1 | *D. melanogaste* | 8,276,228 | SRR10407358 | 83 | 78.3 | 4.7 | 2.6 | 14.4 | 83.1 | 78 | 5.1 | 2.5 | 14.4 | 83 | 79.2 | 3.8 | 2.4 | 14.6 |
| 2 | *D. melanogaste* | 47,552,414 | SRR7716077 | 96.2 | 80.1 | 16.1 | 1.3 | 2.5 | 95.5 | 77.3 | 18.2 | 1.7 | 2.8 | 96.1 | 80.8 | 15.3 | 1.4 | 2.5 |
| 3 | *D. melanogaste* | 45,807,064 | SRR7716078 | 95.2 | 81.8 | 13.4 | 1.8 | 3 | 95.6 | 81.9 | 13.7 | 1.6 | 2.8 | 95.4 | 83.9 | 11.5 | 1.8 | 2.8 |
| 4 | *D. melanogaste* | 29,221,598 | SRR7716079 | 91.1 | 82 | 9.1 | 4 | 4.9 | 91.1 | 82 | 9.1 | 4 | 4.9 | 91.3 | 80.9 | 10.4 | 4 | 4.7 |
| 5 | *D. melanogaste* | 46,936,912 | SRR7716080 | 93.4 | 79.8 | 13.6 | 3.1 | 3.5 | 92.6 | 80.3 | 12.3 | 3.4 | 4 | 93.4 | 80.4 | 13 | 2.8 | 3.8 |

| | | | | 50bp (A) 25,29,33 | | | | | 50bp (B) 25,31,35 | | | | | 50bp (C) 25,33,37 | | | | |
|---|---|---|---|---|---|---|---|---|---|---|---|---|---|---|---|---|---|---|
| Nº | Taxa | Total pe-reads | SRA | Complete % | Single-copy % | Duplicated % | Fragmented % | Missing % | Complete % | Single-copy % | Duplicated % | Fragmented % | Missing % | Complete % | Single-copy % | Duplicated % | Fragmented % | Missing % |
| 1 | *M. musculus* | 33,540,466 | SRR10342165 | 95.9 | 67.8 | 28.1 | 3 | 1.1 | 95.9 | 67.3 | 28.6 | 3 | 1.1 | 95.8 | 67.1 | 28.7 | 3.1 | 1.1 |
| 2 | *M. musculus* | 43,541,512 | SRR10342166 | 97 | 61 | 36 | 1.8 | 1.2 | 97.1 | 62.3 | 34.8 | 1.8 | 1.1 | 97 | 62 | 35 | 1.9 | 1.1 |
| 3 | *M. musculus* | 43,440,008 | SRR10342167 | 97.4 | 62.1 | 35.3 | 1.6 | 1 | 97.4 | 62.9 | 34.5 | 1.6 | 1 | 97.4 | 62.4 | 35 | 1.6 | 1 |
| 4 | *M. musculus* | 33,595,655 | SRR9211836 | 93.9 | 77.8 | 16.1 | 3.9 | 2.2 | 94 | 77.8 | 16.2 | 3.9 | 2.1 | 93.9 | 77.3 | 16.6 | 4 | 2.1 |
| 5 | *M. musculus* | 33,174,108 | SRR9211837 | 93.6 | 79.6 | 14 | 3.9 | 2.5 | 93.6 | 79.8 | 13.8 | 3.9 | 2.5 | 93.6 | 79.6 | 14 | 3.9 | 2.5 |
| 6 | *M. musculus* | 35,030,665 | SRR9211838 | 93.6 | 78.7 | 14.9 | 4.1 | 2.3 | 93.6 | 78 | 15.6 | 4.1 | 2.3 | 93.6 | 78.5 | 15.1 | 4.1 | 2.3 |
| 7 | *M. musculus* | 34,877,241 | SRR9211839 | 93.3 | 78 | 15.3 | 4.3 | 2.4 | 93.3 | 77.6 | 15.7 | 4.3 | 2.4 | 93.3 | 77.8 | 15.5 | 4.3 | 2.4 |
| 1 | *C. elegans* | 24,017,399 | SRR4181026 | 84.6 | 77.4 | 7.2 | 2.1 | 13.3 | 84.5 | 77 | 7.5 | 2.2 | 13.3 | 84.6 | 77.5 | 7.1 | 2.1 | 13.3 |
| 2 | *C. elegans* | 24,409,279 | SRR4181032 | 84.9 | 77.6 | 7.3 | 2.5 | 12.6 | 84.7 | 77.5 | 7.2 | 2.5 | 12.8 | 84.9 | 77.7 | 7.2 | 2.5 | 12.6 |
| 3 | *C. elegans* | 24,694,386 | SRR4181033 | 84.7 | 77.1 | 7.6 | 2.7 | 12.6 | 84.6 | 77.2 | 7.4 | 2.7 | 12.7 | 84.7 | 77.4 | 7.3 | 2.7 | 12.6 |
| 4 | *C. elegans* | 26,159,433 | SRR4181034 | 85.1 | 77 | 8.1 | 2.2 | 12.7 | 85 | 77.1 | 7.9 | 2.4 | 12.6 | 85.1 | 76.4 | 8.7 | 2.2 | 12.7 |
| 5 | *C. elegans* | 26,417,070 | SRR4181036 | 85.1 | 77.3 | 7.8 | 2.6 | 12.3 | 85.1 | 77.1 | 8 | 2.6 | 12.3 | 85 | 76.8 | 8.2 | 2.7 | 12.3 |
| 6 | *C. elegans* | 24,711,402 | SRR4181037 | 85.1 | 78.5 | 6.6 | 1.8 | 13.1 | 85.1 | 78.4 | 6.7 | 1.8 | 13.1 | 85.1 | 78.4 | 6.7 | 1.8 | 13.1 |
| 1 | *D. melanogaste* | 15,648,174 | SRR6743334 | 41.1 | 33.5 | 7.6 | 14.8 | 44.1 | 41.1 | 33 | 8.1 | 14.7 | 44.2 | 40.7 | 32.9 | 7.8 | 15.3 | 44 |
| 2 | *D. melanogaste* | 15,181,806 | SRR6743335 | 38.6 | 30.5 | 8.1 | 16 | 45.4 | 38.6 | 30.2 | 8.4 | 15.7 | 45.7 | 38.7 | 31.2 | 7.5 | 15.4 | 45.9 |
| 3 | *D. melanogaste* | 9,857,308 | SRR6743336 | 31.9 | 26.8 | 5.1 | 16.5 | 51.6 | 31.9 | 26.6 | 5.3 | 16.6 | 51.5 | 31.9 | 26.3 | 5.6 | 16.7 | 51.4 |
| 4 | *D. melanogaste* | 15,940,368 | SRR6743337 | 35.6 | 27.3 | 8.3 | 13.7 | 50.7 | 35.6 | 27.3 | 8.3 | 13.7 | 50.7 | 35.4 | 27.9 | 7.5 | 13.9 | 50.7 |
| 5 | *D. melanogaste* | 18,874,930 | SRR6743338 | 43.3 | 35.6 | 7.7 | 15.6 | 41.1 | 43.4 | 35.4 | 8 | 15.3 | 41.3 | 43.4 | 35.3 | 8.1 | 15.5 | 41.1 |
| 6 | *D. melanogaste* | 21,169,448 | SRR6743339 | 31.9 | 22.9 | 9 | 12.4 | 55.7 | 32.2 | 23 | 9.2 | 12.7 | 55.1 | 32.2 | 23.2 | 9 | 12.2 | 55.6 |
| 7 | *D. melanogaste* | 16,177,860 | SRR6743340 | 37.8 | 29.6 | 8.2 | 13.2 | 49 | 37.9 | 30.5 | 7.4 | 12.7 | 49.4 | 38 | 30 | 8 | 12.6 | 49.4 |
| 8 | *D. melanogaste* | 12,436,324 | SRR6743341 | 35.3 | 28.6 | 6.7 | 14.7 | 50 | 36.3 | 30.5 | 5.8 | 14 | 49.7 | 36 | 29.3 | 6.7 | 14.6 | 49.4 |
| 9 | *D. melanogaste* | 26,395,792 | SRR6743342 | 60.7 | 49.5 | 11.2 | 12.2 | 27.1 | 60.8 | 49.1 | 11.7 | 12.2 | 27 | 60.6 | 49.3 | 11.3 | 12.2 | 27.2 |

| | | | | 75bp (A) 25,29,33,41 | | | | | 75bp (B) 25,31,35,43 | | | | | 75bp (C) 25,33,37,45 | | | | |
|---|---|---|---|---|---|---|---|---|---|---|---|---|---|---|---|---|---|---|
| Nº | Taxa | Total pe-reads | SRA | Complete % | Single-copy % | Duplicated % | Fragmented % | Missing % | Complete % | Single-copy % | Duplicated % | Fragmented % | Missing % | Complete % | Single-copy % | Duplicated % | Fragmented % | Missing % |
| 1 | *M. musculus* | 33,700,156 | SRR10067458 | 97.9 | 65.4 | 32.5 | 1.5 | 0.6 | 98.1 | 66.7 | 31.4 | 1.4 | 0.5 | 98.2 | 66.1 | 32.1 | 1.3 | 0.5 |
| 2 | *M. musculus* | 41,236,457 | SRR10067459 | 98.5 | 58.4 | 40.1 | 1 | 0.5 | 98.6 | 59.8 | 38.8 | 0.9 | 0.5 | 98.4 | 58.2 | 40.2 | 1 | 0.6 |
| 3 | *M. musculus* | 35,598,598 | SRR10067460 | 98.6 | 60.2 | 38.4 | 0.6 | 0.8 | 98.6 | 61.3 | 37.3 | 0.6 | 0.8 | 98.8 | 59.6 | 39.2 | 0.6 | 0.6 |
| 4 | *M. musculus* | 41,745,958 | SRR10067461 | 93.6 | 51.1 | 42.5 | 4.1 | 2.3 | 93.4 | 51.2 | 42.2 | 4.3 | 2.3 | 93.4 | 52 | 41.4 | 4.3 | 2.3 |
| 5 | *M. musculus* | 45,329,544 | SRR10067462 | 96.4 | 54.6 | 41.8 | 1.8 | 1.8 | 96.4 | 54.1 | 42.3 | 1.8 | 1.8 | 96.3 | 55.4 | 40.9 | 1.9 | 1.8 |
| 6 | *M. musculus* | 44,469,310 | SRR10067463 | 94 | 52.2 | 41.8 | 3.7 | 2.3 | 94 | 52.2 | 41.8 | 3.7 | 2.3 | 93.9 | 52.1 | 41.8 | 3.9 | 2.2 |
| 1 | *C. elegans* | 40,302,838 | SRR1585277 | 86.2 | 69.7 | 16.5 | 2.1 | 11.7 | 86.2 | 70.2 | 16 | 2.1 | 11.7 | 86.2 | 69.4 | 16.8 | 2.1 | 11.7 |
| 2 | *C. elegans* | 50,516,835 | SRR1585278 | 86.4 | 68.8 | 17.6 | 1.8 | 11.8 | 86.3 | 68.5 | 17.8 | 1.8 | 11.9 | 86.5 | 69.4 | 17.1 | 1.8 | 11.7 |
| 3 | *C. elegans* | 41,947,175 | SRR8257107 | 84.8 | 75.6 | 9.2 | 2.4 | 12.8 | 84.7 | 75.4 | 9.3 | 2.4 | 12.9 | 84.7 | 75.8 | 8.9 | 2.4 | 12.9 |
| 4 | *C. elegans* | 44,969,393 | SRR8257108 | 84.7 | 75.6 | 9.1 | 1.9 | 13.4 | 84.7 | 75.7 | 9 | 1.9 | 13.4 | 84.7 | 75.1 | 9.6 | 1.9 | 13.4 |
| 5 | *C. elegans* | 45,605,396 | SRR8257109 | 85.1 | 75.8 | 9.3 | 1.7 | 13.2 | 85 | 76.4 | 8.6 | 1.7 | 13.3 | 85.1 | 76.2 | 8.9 | 1.7 | 13.2 |
| 1 | *D. melanogaste* | 87,423,452 | SRR8559044 | 98 | 57.1 | 40.9 | 0.8 | 1.2 | 97.9 | 58.6 | 39.3 | 0.9 | 1.2 | 97.9 | 58.5 | 39.4 | 0.9 | 1.2 |
| 2 | *D. melanogaste* | 85,714,154 | SRR8559045 | 98.5 | 54.1 | 44.4 | 0.6 | 0.9 | 98.4 | 53.5 | 44.9 | 0.7 | 0.9 | 98.5 | 54.6 | 43.9 | 0.6 | 0.9 |
| 3 | *D. melanogaste* | 88,252,694 | SRR8559046 | 98.3 | 52.5 | 45.8 | 0.6 | 1.1 | 98.4 | 53.5 | 44.9 | 0.5 | 1.1 | 98.4 | 53.1 | 45.3 | 0.5 | 1.1 |
| 4 | *D. melanogaste* | 82,110,608 | SRR8559047 | 98.1 | 54.7 | 43.4 | 0.7 | 1.2 | 97.9 | 55 | 42.9 | 0.8 | 1.3 | 98.1 | 54.4 | 43.7 | 0.7 | 1.2 |
| 5 | *D. melanogaste* | 102,413,880 | SRR8559048 | 98.3 | 49.2 | 49.1 | 0.8 | 0.9 | 98.3 | 49.7 | 48.6 | 0.8 | 0.9 | 98.3 | 49.4 | 48.9 | 0.8 | 0.9 |
| 6 | *D. melanogaste* | 98,529,578 | SRR8559049 | 98.2 | 55.4 | 42.8 | 0.8 | 1 | 98.2 | 56.6 | 41.6 | 0.9 | 0.9 | 98.3 | 55.8 | 42.5 | 0.8 | 0.9 |

| | | | | 100bp (A) 25,35,47,61 | | | | | 100bp (B) 25,37,55,63 | | | | | 100bp (C) 25,41,57,67 | | | | |
|---|---|---|---|---|---|---|---|---|---|---|---|---|---|---|---|---|---|---|
| Nº | Taxa | Total pe-reads | SRA | Complete % | Single-copy % | Duplicated % | Fragmented % | Missing % | Complete % | Single-copy % | Duplicated % | Fragmented % | Missing % | Complete % | Single-copy % | Duplicated % | Fragmented % | Missing % |
| 1 | *M. musculus* | 182,180,123 | SRR1017092 | 98.9 | 34.3 | 64.6 | 0.8 | 0.3 | 98.9 | 31.5 | 67.4 | 0.8 | 0.3 | 98.7 | 32.9 | 65.8 | 0.9 | 0.4 |
| 2 | *M. musculus* | 107,280,600 | SRR1017093 | 98.8 | 35.2 | 63.6 | 0.9 | 0.3 | 99.2 | 34.7 | 64.5 | 0.5 | 0.3 | 98.9 | 35.4 | 63.5 | 0.5 | 0.6 |
| 3 | *M. musculus* | 127,017,374 | SRR1017094 | 99.1 | 33.6 | 65.5 | 0.5 | 0.4 | 99.1 | 35 | 64.1 | 0.6 | 0.3 | 98.9 | 33.8 | 65.1 | 0.7 | 0.4 |
| 4 | *M. musculus* | 102,909,165 | SRR5171077 | 43.3 | 34.7 | 8.6 | 30.4 | 26.3 | 43.2 | 34.5 | 8.7 | 30.5 | 26.3 | 43.4 | 34.4 | 9 | 30.3 | 26.3 |
| 5 | *M. musculus* | 65,372,078 | SRR5171085 | 98.1 | 40.3 | 57.8 | 1.8 | 0.1 | 98 | 39.5 | 58.5 | 1.9 | 0.1 | 98 | 41.5 | 56.5 | 1.8 | 0.2 |
| 6 | *M. musculus* | 27,016,597 | SRR5171095 | 55.3 | 39.6 | 15.7 | 28.2 | 16.5 | 55.7 | 40.8 | 14.9 | 27.7 | 16.6 | 55.3 | 40.6 | 14.7 | 28.1 | 16.6 |
| 1 | *C. elegans* | 49,743,412 | SRR2142254 | 82.1 | 68.8 | 13.3 | 3.9 | 14 | 82.1 | 67.8 | 14.3 | 3.9 | 14 | 82 | 68.2 | 13.8 | 3.9 | 14.1 |
| 2 | *C. elegans* | 38,836,876 | SRR2142255 | 82.1 | 70.9 | 11.2 | 3.3 | 14.6 | 82.1 | 69.6 | 12.5 | 3.3 | 14.6 | 82.1 | 70.8 | 11.3 | 3.3 | 14.6 |
| 3 | *C. elegans* | 11,166,310 | SRR2015313 | 63.5 | 46 | 17.5 | 17.2 | 19.3 | 63.5 | 45.8 | 17.7 | 17.2 | 19.3 | 63.5 | 46.1 | 17.4 | 17.2 | 19.3 |
| 4 | *C. elegans* | 11,626,153 | SRR2015311 | 79.4 | 59.5 | 19.9 | 6 | 14.6 | 79.5 | 60.1 | 19.4 | 5.9 | 14.6 | 79.3 | 59.5 | 19.8 | 6 | 14.7 |
| 5 | *C. elegans* | 12,567,755 | SRR2012785 | 85.7 | 68.9 | 16.8 | 2.6 | 11.7 | 85.7 | 67.6 | 18.1 | 2.6 | 11.7 | 85.7 | 68.1 | 17.6 | 2.6 | 11.7 |
| 6 | *C. elegans* | 14,372,593 | SRR2012781 | 69.4 | 59.1 | 10.3 | 12.7 | 17.9 | 69.4 | 58.2 | 11.2 | 12.7 | 17.9 | 69.3 | 58.7 | 10.6 | 12.8 | 17.9 |
| 1 | *D. melanogaste* | 30,515,068 | SRR3018852 | 97.2 | 61.2 | 36 | 0.6 | 2.2 | 97.2 | 60.9 | 36.3 | 0.6 | 2.2 | 97.3 | 61 | 36.3 | 0.5 | 2.2 |
| 2 | *D. melanogaste* | 24,284,630 | SRR3018855 | 96.5 | 62.1 | 34.4 | 1 | 2.5 | 96.6 | 64 | 32.6 | 1 | 2.4 | 96.6 | 62.8 | 33.8 | 1 | 2.4 |
| 3 | *D. melanogaste* | 24,773,404 | SRR3018862 | 96.9 | 64.9 | 32 | 1.2 | 1.9 | 97 | 64.6 | 32.4 | 1.1 | 1.9 | 96.9 | 63.9 | 33 | 1.1 | 2 |
| 4 | *D. melanogaste* | 30,140,704 | SRR3019045 | 97 | 61.5 | 35.5 | 0.9 | 2.1 | 97.1 | 61.7 | 35.4 | 0.8 | 2.1 | 97 | 60.2 | 36.8 | 0.9 | 2.1 |
| 5 | *D. melanogaste* | 25,828,680 | SRR3019179 | 96.6 | 62 | 34.6 | 1.1 | 2.3 | 96.4 | 62.8 | 33.6 | 1.1 | 2.5 | 96.6 | 61.6 | 35 | 1.1 | 2.3 |
| 6 | *D. melanogaste* | 19,049,236 | SRR3019184 | 96.3 | 67.4 | 28.9 | 1.3 | 2.4 | 96.4 | 66.8 | 29.6 | 1.3 | 2.3 | 96.4 | 66.8 | 29.6 | 1.3 | 2.3 |

| | | | | 150bp (A) 25,31,41,51,65 | | | | | 150bp (B) 25,33,43,53,71 | | | | | 150bp (C) 25,35,55,75,85 | | | | |
|---|---|---|---|---|---|---|---|---|---|---|---|---|---|---|---|---|---|---|
| Nº | Taxa | Total pe-reads | SRA | Complete % | Single-copy % | Duplicated % | Fragmented % | Missing % | Complete % | Single-copy % | Duplicated % | Fragmented % | Missing % | Complete % | Single-copy % | Duplicated % | Fragmented % | Missing % |
| 1 | *M. musculus* | 21,173,335 | SRR10560364 | 98.4 | 48.7 | 49.7 | 0.6 | 12 | 98.4 | 47.8 | 50.6 | 0.6 | 1 | 98.4 | 48.8 | 49.6 | 0.6 | 1 |
| 2 | *M. musculus* | 22,731,971 | SRR10560365 | 98.6 | 45.3 | 53.3 | 0.7 | 0.7 | 98.5 | 45.9 | 52.6 | 0.6 | 0.9 | 98.5 | 46.7 | 51.8 | 0.6 | 0.9 |
| 3 | *M. musculus* | 68,712,046 | SRR8329325 | 95.5 | 39.3 | 56.2 | 3.2 | 1.3 | 95.5 | 38.1 | 57.4 | 3.2 | 1.3 | 95.5 | 38.5 | 57 | 3.2 | 1.3 |
| 4 | *M. musculus* | 59,012,049 | SRR8329326 | 97.3 | 36.6 | 60.7 | 1.5 | 1.2 | 97.2 | 36.2 | 61 | 1.6 | 1.2 | 97.5 | 36.9 | 60.6 | 1.3 | 1.2 |
| 5 | *M. musculus* | 21,930,527 | SRR7079069 | 97.6 | 41.6 | 56 | 1.6 | 0.8 | 97.5 | 40 | 57.5 | 1.6 | 0.9 | **97.9** | **40.4** | **57.5** | **1.4** | **0.7** |
| 1 | *C. elegans* | 20,345,696 | SRR10407359 | 86.8 | 71.8 | 15 | 1.6 | 11.6 | 86.8 | 72 | 14.8 | 1.6 | 11.6 | 86.8 | 72.9 | 13.9 | 1.6 | 11.6 |
| 2 | *C. elegans* | 19,948,493 | SRR10407358 | 87 | 71.6 | 15.4 | 1.5 | 11.5 | 87 | 70.8 | 16.2 | 1.5 | 11.5 | 86.8 | 71.5 | 15.3 | 1.6 | 11.6 |
| 3 | *C. elegans* | 21,276,093 | SRR10407357 | 86.6 | 71.4 | 15.2 | 1.9 | 11.5 | 86.7 | 70.2 | 16.5 | 1.8 | 11.5 | 86.7 | 71.1 | 15.6 | 1.8 | 11.5 |
| 4 | *C. elegans* | 23,993,704 | SRR10407356 | 86.3 | 74 | 12.3 | 2 | 11.7 | 86.2 | 74.3 | 11.9 | 2 | 11.8 | 86.4 | 73 | 13.4 | 1.9 | 11.7 |
| 5 | *C. elegans* | 25,110,365 | SRR10407355 | 86.5 | 72 | 14.5 | 1.6 | 11.9 | 86.4 | 73.2 | 13.2 | 1.7 | 11.9 | 86.2 | 72.3 | 13.9 | 1.7 | 12.1 |
| 6 | *C. elegans* | 22,967,010 | SRR2012781 | 86 | 70.3 | 15.7 | 1.9 | 12.1 | 86.3 | 71.2 | 15.1 | 1.9 | 11.8 | 86.3 | 71.1 | 15.2 | 1.9 | 11.8 |
| 1 | *D. melanogaste* | 8,276,228 | SRR10407358 | 84.4 | 74.7 | 9.7 | 2.5 | 13.1 | 84.4 | 75.3 | 9.1 | 2.6 | 13 | 84.5 | 75.1 | 9.4 | 2.5 | 13 |
| 2 | *D. melanogaste* | 47,552,414 | SRR7716077 | 97.3 | 57.6 | 39.7 | 0.8 | 1.9 | 97.1 | 58 | 39.1 | 0.9 | 2 | 97.3 | 58.2 | 39.1 | 0.8 | 1.9 |
| 3 | *D. melanogaste* | 45,807,064 | SRR7716078 | 96.4 | 61.5 | 34.9 | 1.5 | 2.1 | 96.4 | 62.4 | 34 | 1.5 | 2.1 | 96.6 | 62.1 | 34.5 | 1.4 | 2 |
| 4 | *D. melanogaste* | 29,221,598 | SRR7716079 | 91.7 | 64 | 27.7 | 4.6 | 3.7 | 91.6 | 63.5 | 28.1 | 4.7 | 3.7 | 91.7 | 63.5 | 28.2 | 4.7 | 3.6 |
| 5 | *D. melanogaste* | 46,936,912 | SRR7716080 | 94.2 | 63.7 | 30.5 | 3.1 | 2.7 | 94.2 | 62.3 | 31.9 | 3 | 2.8 | 94.1 | 64.1 | 30 | 3 | 2.9 |

| Nº | Taxa | Total pe-reads | SRA | 50bp (A) 25,29,33 | | | | | 50bp (B) 25,31,35 | | | | | 50bp (C) 25,33,37 | | | | |
|---|---|---|---|---|---|---|---|---|---|---|---|---|---|---|---|---|---|---|
| | | | | Complete % | Single-copy % | Duplicated % | Fragmented % | Missing % | Complete % | Single-copy % | Duplicated % | Fragmented % | Missing % | Complete % | Single-copy % | Duplicated % | Fragmented % | Missing % |
| 1 | *M. musculus* | 33,540,466 | SRR10342165 | 96.8 | 86.5 | 10.3 | 0.8 | 2.4 | 96.6 | 86.9 | 9.7 | 1.2 | 2.2 | 97.1 | 87.1 | 10 | 0.7 | 2.2 |
| 2 | *M. musculus* | 43,541,512 | SRR10342166 | 96.4 | 86.5 | 9.9 | 1.2 | 2.4 | 96.7 | 86.8 | 9.9 | 1 | 2.3 | 96.5 | 85.4 | 11.1 | 1 | 2.5 |
| 3 | *M. musculus* | 43,440,008 | SRR10342167 | 97 | 86.8 | 10.2 | 0.7 | 2.3 | 96.8 | 87.5 | 9.3 | 1 | 2.2 | 97.1 | 86.8 | 10.3 | 0.8 | 2.1 |
| 4 | *M. musculus* | 33,595,655 | SRR9211836 | 95.5 | 86.1 | 9.4 | 0.9 | 3.6 | 95.8 | 87 | 8.8 | 0.8 | 3.4 | 95.4 | 86.2 | 9.2 | 1 | 3.6 |
| 5 | *M. musculus* | 33,174,108 | SRR9211837 | 93.8 | 84.6 | 9.2 | 1.9 | 4.3 | 93.9 | 85 | 8.9 | 2 | 4.1 | 94 | 84.5 | 9.5 | 1.9 | 4.1 |
| 6 | *M. musculus* | 35,030,665 | SRR9211838 | 94.9 | 86.4 | 8.5 | 1.5 | 3.6 | 95.2 | 85.6 | 9.6 | 1.3 | 3.5 | 95.3 | 86.6 | 8.7 | 1.3 | 3.4 |
| 7 | *M. musculus* | 34,877,241 | SRR9211839 | 94.6 | 85.2 | 9.4 | 1.4 | 4 | 94.8 | 84.1 | 10.7 | 1.4 | 3.8 | 94.8 | 84.4 | 10.4 | 1.4 | 3.8 |
| 1 | *C. elegans* | 24,017,399 | SRR4181026 | 75.2 | 70.2 | 5 | 2.7 | 22.1 | 75.4 | 70.6 | 4.8 | 2.6 | 22 | 75.2 | 70.4 | 4.8 | 2.7 | 22.1 |
| 2 | *C. elegans* | 24,409,279 | SRR4181032 | 75.4 | 70.3 | 5.1 | 2.2 | 22.4 | 75.2 | 70.1 | 5.1 | 2.2 | 22.6 | 75.5 | 69.8 | 5.7 | 2.2 | 22.3 |
| 3 | *C. elegans* | 24,694,386 | SRR4181033 | 75.7 | 71 | 4.7 | 2.2 | 22.1 | 76 | 72.1 | 3.9 | 2.2 | 21.8 | 75.8 | 71.6 | 4.2 | 2.2 | 22 |
| 4 | *C. elegans* | 26,159,433 | SRR4181034 | 75.7 | 70.4 | 5.3 | 2.3 | 22 | 76.1 | 70.6 | 5.5 | 2.1 | 21.8 | 75.9 | 71 | 4.9 | 2.3 | 21.8 |
| 5 | *C. elegans* | 26,417,070 | SRR4181036 | 75.6 | 71.1 | 4.5 | 2.1 | 22.3 | 75.8 | 70.9 | 4.9 | 2 | 22.2 | 75.9 | 70.9 | 5 | 2 | 22.1 |
| 6 | *C. elegans* | 24,711,402 | SRR4181037 | 75.3 | 69.6 | 5.7 | 2.4 | 22.3 | 75.3 | 69.6 | 5.7 | 2.3 | 22.4 | 75.1 | 70 | 5.1 | 2.3 | 22.6 |
| 1 | *D. melanogaste* | 15,648,174 | SRR6743334 | 39.4 | 36.8 | 2.6 | 5.9 | 54.7 | 39.5 | 36.6 | 2.9 | 6.5 | 54 | 39.8 | 37.1 | 2.7 | 6 | 54.2 |
| 2 | *D. melanogaste* | 15,181,806 | SRR6743335 | 38.4 | 36.8 | 1.6 | 6.4 | 55.2 | 38.1 | 36.3 | 1.8 | 5.8 | 56.1 | 37.9 | 36.2 | 1.7 | 6.4 | 55.7 |
| 3 | *D. melanogaste* | 9,857,308 | SRR6743336 | 30.6 | 28.5 | 2.1 | 6.7 | 62.7 | 30.1 | 27.9 | 2.2 | 6.5 | 63.4 | 30.1 | 28.4 | 1.7 | 6.6 | 63.3 |
| 4 | *D. melanogaste* | 15,940,368 | SRR6743337 | 34.3 | 31.9 | 2.4 | 4.5 | 61.2 | 33.5 | 31.1 | 2.4 | 5 | 61.5 | 34.6 | 32.6 | 2 | 4.9 | 60.5 |
| 5 | *D. melanogaste* | 18,874,930 | SRR6743338 | 42.7 | 40 | 2.7 | 6.4 | 50.9 | 41.8 | 39.1 | 2.7 | 6.9 | 51.3 | 42.5 | 40.1 | 2.4 | 6.6 | 50.9 |
| 6 | *D. melanogaste* | 21,169,448 | SRR6743339 | 29.8 | 28.1 | 1.7 | 6.7 | 63.5 | 29.7 | 28.2 | 1.5 | 7.1 | 63.2 | 29.5 | 28.2 | 1.3 | 6.2 | 64.3 |
| 7 | *D. melanogaste* | 16,177,860 | SRR6743340 | 37.7 | 35.5 | 2.2 | 5.3 | 57 | 37.1 | 35 | 2.1 | 5.1 | 57.8 | 38.2 | 36.5 | 1.7 | 4.8 | 57 |
| 8 | *D. melanogaste* | 12,436,324 | SRR6743341 | 33.5 | 30.8 | 2.7 | 6.4 | 60.1 | 33.7 | 31 | 2.7 | 6.5 | 59.8 | 33.9 | 32 | 1.9 | 6.8 | 59.3 |
| 9 | *D. melanogaste* | 26,395,792 | SRR6743342 | 62.7 | 57 | 5.7 | 5.9 | 31.4 | 62.3 | 57.9 | 4.4 | 6.4 | 31.3 | 62.3 | 57.2 | 5.1 | 6.2 | 31.5 |

| Nº | Taxa | Total pe-reads | SRA | 75bp (A) 25,29,33,41 | | | | | 75bp (B) 25,31,35,43 | | | | | 75bp (C) 25,33,37,45 | | | | |
|---|---|---|---|---|---|---|---|---|---|---|---|---|---|---|---|---|---|---|
| | | | | Complete % | Single-copy % | Duplicated % | Fragmented % | Missing % | Complete % | Single-copy % | Duplicated % | Fragmented % | Missing % | Complete % | Single-copy % | Duplicated % | Fragmented % | Missing % |
| 1 | *M. musculus* | 33,700,156 | SRR10067458 | 96 | 84.5 | 11.5 | 1.9 | 2.1 | 95.7 | 83.2 | 12.5 | 2 | 2.3 | 96 | 84.1 | 11.9 | 2 | 2 |
| 2 | *M. musculus* | 41,236,457 | SRR10067459 | 96.6 | 83.2 | 13.4 | 1.2 | 2.2 | 96.9 | 83.5 | 13.4 | 0.8 | 2.3 | 97.3 | 83.8 | 13.5 | 0.8 | 1.9 |
| 3 | *M. musculus* | 35,598,598 | SRR10067460 | 96.2 | 82.8 | 13.4 | 0.8 | 3 | 96.4 | 83.8 | 12.6 | 0.9 | 2.7 | 96.1 | 83.3 | 12.8 | 0.9 | 3 |
| 4 | *M. musculus* | 41,745,958 | SRR10067461 | 94.4 | 84.4 | 10 | 1.9 | 3.7 | 93.9 | 83.9 | 10 | 2.1 | 4 | 94.4 | 84.2 | 10.2 | 1.9 | 3.7 |
| 5 | *M. musculus* | 45,329,544 | SRR10067462 | 95.8 | 85.1 | 10.7 | 1.2 | 3 | 96 | 86 | 10 | 1.2 | 2.8 | 95.8 | 85.2 | 10.6 | 0.9 | 3.3 |
| 6 | *M. musculus* | 44,469,310 | SRR10067463 | 94.5 | 84.6 | 9.9 | 2.2 | 3.3 | 94.6 | 84.4 | 10.2 | 2 | 3.4 | 94.5 | 84.5 | 10 | 2.1 | 3.4 |
| 1 | *C. elegans* | 40,302,838 | SRR1585277 | 75.3 | 65.6 | 9.7 | 2.4 | 22.3 | 75.6 | 66.7 | 8.9 | 2.5 | 21.9 | 75.4 | 66.1 | 9.3 | 2.4 | 22.2 |
| 2 | *C. elegans* | 50,516,835 | SRR1585278 | 75.5 | 63.9 | 11.6 | 2.5 | 22 | 75.4 | 63.9 | 11.5 | 2.2 | 22.4 | 75.2 | 64.4 | 10.8 | 2.2 | 22.6 |
| 3 | *C. elegans* | 41,947,175 | SRR8257107 | 73.9 | 69.9 | 4 | 2.7 | 23.4 | 74.2 | 69.7 | 4.5 | 2.6 | 23.2 | 74.7 | 70.4 | 4.3 | 2.5 | 22.8 |
| 4 | *C. elegans* | 44,969,393 | SRR8257108 | 74.7 | 71.6 | 3.1 | 2.5 | 22.8 | 74.9 | 71.4 | 3.5 | 2.3 | 22.8 | 75 | 71.5 | 3.5 | 2.2 | 22.8 |
| 5 | *C. elegans* | 45,605,396 | SRR8257109 | 74.6 | 70.2 | 4.4 | 2.2 | 23.2 | 75.1 | 71.1 | 4 | 2.3 | 22.6 | 74.7 | 70.8 | 3.9 | 2.3 | 23 |
| 1 | *D. melanogaste* | 87,423,452 | SRR8559044 | 94.7 | 76.1 | 18.6 | 1 | 4.3 | 94.5 | 75.9 | 18.6 | 0.9 | 4.6 | 95.3 | 77.5 | 17.8 | 0.7 | 4 |
| 2 | *D. melanogaste* | 85,714,154 | SRR8559045 | 94.6 | 75.7 | 18.9 | 0.8 | 4.6 | 95.2 | 78.5 | 16.7 | 0.6 | 4.2 | 94.8 | 76.7 | 18.1 | 0.6 | 4.6 |
| 3 | *D. melanogaste* | 88,252,694 | SRR8559046 | 95.2 | 75.6 | 19.6 | 0.8 | 4 | 95.2 | 77.3 | 17.9 | 0.6 | 4.2 | 95.5 | 78.1 | 17.4 | 0.5 | 4 |
| 4 | *D. melanogaste* | 82,110,608 | SRR8559047 | 95.5 | 77.6 | 17.9 | 0.4 | 4.1 | 95.5 | 77.3 | 18.2 | 0.6 | 3.9 | 95.7 | 78.6 | 17.1 | 0.7 | 3.6 |
| 5 | *D. melanogaste* | 102,413,880 | SRR8559048 | 94.7 | 72.1 | 22.6 | 0.9 | 4.4 | 94.9 | 72.3 | 22.6 | 0.9 | 4.2 | 94.8 | 73.5 | 21.3 | 1.2 | 4 |
| 6 | *D. melanogaste* | 98,529,578 | SRR8559049 | 95 | 75.3 | 19.7 | 0.6 | 4.4 | 95.2 | 76.4 | 18.8 | 0.6 | 4.2 | 94.4 | 77.4 | 17 | 0.8 | 4.8 |

| Nº | Taxa | Total pe-reads | SRA | 100bp (A) 25,35,47,61 | | | | | 100bp (B) 25,37,55,63 | | | | | 100bp (C) 25,41,57,67 | | | | |
|---|---|---|---|---|---|---|---|---|---|---|---|---|---|---|---|---|---|---|
| | | | | Complete % | Single-copy % | Duplicated % | Fragmented % | Missing % | Complete % | Single-copy % | Duplicated % | Fragmented % | Missing % | Complete % | Single-copy % | Duplicated % | Fragmented % | Missing % |
| 1 | *M. musculus* | 182,180,123 | **SRR1017092** | 95.3 | 79.6 | 15.7 | 1.8 | 2.9 | 95.7 | 78.6 | 17.1 | 1.5 | 2.8 | 95.3 | 77.9 | 17.4 | 1.5 | 3.2 |
| 2 | *M. musculus* | 107,280,600 | **SRR1017093** | 96 | 76.5 | 19.5 | 1.8 | 2.2 | 96 | 80.7 | 15.3 | 1.4 | 2.6 | 96.9 | 78.8 | 18.1 | 1.2 | 1.9 |
| 3 | *M. musculus* | 127,017,374 | **SRR1017094** | 96.2 | 80.9 | 15.3 | 0.7 | 3.1 | 96 | 79.4 | 16.6 | 1.3 | 2.7 | 96.8 | 80.8 | 16 | 0.9 | 2.3 |
| 5 | *M. musculus* | 102,909,165 | SRR5171077 | 96.2 | 81 | 15.2 | 2 | 1.8 | 96.1 | 82.7 | 13.4 | 2.1 | 1.8 | 33.9 | 33.3 | 0.6 | 23.1 | 43 |
| 6 | *M. musculus* | 65,372,078 | SRR5171085 | 96.4 | 81.4 | 15 | 1.5 | 2.1 | 96.7 | 82.5 | 14.2 | 1.3 | 2 | 96.1 | 80.2 | 15.9 | 1.7 | 2.2 |
| 7 | *M. musculus* | 27,016,597 | SRR5171095 | 45.3 | 43.1 | 2.2 | 21.4 | 33.3 | 44.9 | 42.7 | 2.2 | 21.3 | 33.8 | 45 | 43 | 2 | 21.4 | 33.6 |
| 1 | *C. elegans* | 49,743,412 | SRR2142254 | 73.1 | 70.1 | 3 | 3.1 | 23.8 | 73.3 | 70.9 | 2.4 | 3.1 | 23.6 | 73.3 | 70.6 | 2.7 | 2.9 | 23.8 |
| 2 | *C. elegans* | 38,836,876 | SRR2142255 | 73.5 | 72.1 | 1.4 | 2.8 | 23.7 | 73.4 | 71.4 | 2 | 3 | 23.6 | 73.2 | 70.9 | 2.3 | 3.2 | 23.6 |
| 3 | *C. elegans* | 11,166,310 | SRR2015313 | 56.6 | 55.1 | 1.5 | 11.6 | 31.8 | 56.4 | 55.6 | 0.8 | 12.1 | 31.5 | 56.4 | 55.5 | 0.9 | 12.1 | 31.5 |
| 4 | *C. elegans* | 11,626,153 | SRR2015311 | 69.5 | 67.6 | 1.9 | 5.6 | 24.9 | 69.7 | 67.9 | 1.8 | 5.7 | 24.6 | 69.7 | 68.1 | 1.6 | 5.7 | 24.6 |
| 5 | *C. elegans* | 12,567,755 | SRR2012785 | 74.8 | 64.7 | 10.1 | 2.6 | 22.6 | 75.1 | 65.1 | 10 | 2.5 | 22.4 | 75.1 | 64.2 | 10.9 | 2.4 | 22.5 |
| 6 | *C. elegans* | 14,372,593 | SRR2012781 | 53.9 | 48.7 | 5.2 | 13.1 | 33 | 53.5 | 48.7 | 4.8 | 13.3 | 33.2 | 53.7 | 48.6 | 5.1 | 13.2 | 33.1 |
| 1 | *D. melanogaste* | 30,515,068 | SRR3018852 | 94.1 | 81.1 | 13 | 0.5 | 5.4 | 94.1 | 78.4 | 15.7 | 0.5 | 5.4 | 94.7 | 80.3 | 14.4 | 0.4 | 4.9 |
| 2 | *D. melanogaste* | 24,284,630 | SRR3018855 | 93.5 | 77.3 | 16.2 | 0.9 | 5.6 | 93.5 | 78.2 | 15.3 | 0.8 | 5.7 | 93.6 | 76.9 | 16.7 | 1.3 | 5.1 |
| 3 | *D. melanogaste* | 24,773,404 | SRR3018862 | 94.2 | 82.3 | 11.9 | 0.8 | 5 | 94.7 | 82 | 12.7 | 0.9 | 4.4 | 94.9 | 83.9 | 11 | 0.6 | 4.5 |
| 4 | *D. melanogaste* | 30,140,704 | SRR3019045 | 94.4 | 80.6 | 13.8 | 0.4 | 5.2 | 94.2 | 80 | 14.2 | 0.6 | 5.2 | 93.7 | 80.5 | 13.2 | 0.5 | 5.8 |
| 5 | *D. melanogaste* | 25,828,680 | SRR3019179 | 94 | 80.8 | 13.2 | 0.5 | 5.5 | 93.9 | 80.8 | 13.1 | 0.6 | 5.5 | 94.1 | 81.4 | 12.7 | 0.4 | 5.5 |
| 6 | *D. melanogaste* | 19,049,236 | SRR3019184 | 93.6 | 80.9 | 12.7 | 0.9 | 5.5 | 93.6 | 82.5 | 11.1 | 1.2 | 5.2 | 93.7 | 82.5 | 11.2 | 1.2 | 5.1 |

| Nº | Taxa | Total pe-reads | SRA | 150bp (A) 25,31,41,51,65 | | | | | 150bp (B) 25,33,43,53,71 | | | | | 150bp (C) 25,35,55,75,85 | | | | |
|---|---|---|---|---|---|---|---|---|---|---|---|---|---|---|---|---|---|---|
| | | | | Complete % | Single-copy % | Duplicated % | Fragmented % | Missing % | Complete % | Single-copy % | Duplicated % | Fragmented % | Missing % | Complete % | Single-copy % | Duplicated % | Fragmented % | Missing % |
| 1 | *M. musculus* | 21,173,335 | SRR10560364 | 96.9 | 82.2 | 14.7 | 1.6 | 1.5 | 96.8 | 83.3 | 13.5 | 1.5 | 1.7 | 96.7 | 83.4 | 13.3 | 1.6 | 1.7 |
| 2 | *M. musculus* | 22,731,971 | SRR10560365 | 96.4 | 79.1 | 17.3 | 1.5 | 2.1 | 96.7 | 80.3 | 16.4 | 1.4 | 1.9 | 96.5 | 79.9 | 16.6 | 1.2 | 2.3 |
| 3 | *M. musculus* | 68,712,046 | SRR8329325 | 94.5 | 84.3 | 10.2 | 1.6 | 3.9 | 94.6 | 85.1 | 9.5 | 2.1 | 3.3 | 94 | 84.6 | 9.4 | 2.4 | 3.6 |
| 4 | *M. musculus* | 59,012,049 | SRR8329326 | 96 | 82.7 | 13.3 | 1.3 | 3.7 | 95.6 | 83 | 12.6 | 1.7 | 2.7 | 95.9 | 82.6 | 13.3 | 1.3 | 2.8 |
| 5 | *M. musculus* | 21,930,527 | SRR7079069 | 93.6 | 78.3 | 15.3 | 2.6 | 3.8 | 94.3 | 80 | 14.3 | 2.1 | 3.6 | | | | | |
| 1 | *C. elegans* | 20,345,696 | SRR10407359 | 75.6 | 69.1 | 6.5 | 2.4 | 22 | 75.3 | 68.1 | 7.2 | 2.7 | 22 | 75.5 | 69.5 | 6 | 2.7 | 21.8 |
| 2 | *C. elegans* | 19,948,493 | SRR10407358 | 76 | 69.3 | 6.7 | 2.2 | 21.8 | 76.1 | 69.4 | 6.7 | 2.1 | 21.8 | 76.1 | 69.6 | 6.5 | 2.1 | 21.8 |
| 3 | *C. elegans* | 21,276,093 | SRR10407357 | 75.4 | 67.5 | 7.9 | 2.5 | 22.1 | 75.4 | 67.1 | 8.3 | 2.3 | 22.3 | 75.7 | 66.2 | 9.5 | 2.3 | 22 |
| 4 | *C. elegans* | 23,993,704 | SRR10407356 | 75.5 | 67.5 | 8 | 2.5 | 22 | 75.4 | 68.4 | 7 | 2.4 | 22.2 | 75.7 | 67.5 | 8.2 | 2.3 | 22 |
| 5 | *C. elegans* | 25,110,365 | SRR10407355 | 75.7 | 68.3 | 7.4 | 2.1 | 22.2 | 75.2 | 67.7 | 7.5 | 2.3 | 22.5 | 75.6 | 68.2 | 7.4 | 2.3 | 22.1 |
| 6 | *C. elegans* | 22,967,010 | SRR10407354 | 75.2 | 68.4 | 6.8 | 2.8 | 22 | 75.4 | 68.8 | 6.6 | 2.7 | 21.9 | 74.9 | 68 | 6.9 | 2.8 | 22.3 |
| 1 | *D. melanogaste* | 8,276,228 | SRR10407358 | 74.2 | 71 | 3.2 | 3 | 22.8 | 74 | 70.1 | 3.9 | 2.9 | 23.1 | 74.3 | 71.3 | 3 | 3 | 22.7 |
| 2 | *D. melanogaste* | 47,552,414 | SRR7716077 | 94.5 | 80 | 14.5 | 1.3 | 4.2 | 93.8 | 77.8 | 16 | 1.3 | 4.9 | 94.8 | 81.4 | 13.4 | 1.2 | 4 |
| 3 | *D. melanogaste* | 45,807,064 | SRR7716078 | 93.4 | 81.7 | 11.7 | 1 | 5.6 | 93.2 | 81.4 | 11.8 | 1 | 5.8 | 93.8 | 83.6 | 10.2 | 1.2 | 5 |
| 4 | *D. melanogaste* | 29,221,598 | SRR7716079 | 87.2 | 78.8 | 8.4 | 4 | 8.8 | 87.2 | 79.8 | 7.4 | 4 | 8.8 | 86.9 | 78.1 | 8.8 | 4 | 9.1 |
| 5 | *D. melanogaste* | 46,936,912 | SRR7716080 | 91.2 | 81.1 | 10.1 | 2.4 | 6.4 | 91.3 | 82.5 | 8.8 | 2.3 | 6.4 | 91 | 79.9 | 11.1 | 2.1 | 6.9 |

| | | | | 50bp (A) 25,29,33 | | | | | 50bp (B) 25,31,35 | | | | | 50bp (C) 25,33,37 | | | | |
|---|---|---|---|---|---|---|---|---|---|---|---|---|---|---|---|---|---|---|
| **Nº** | **Taxa** | **Total pe-reads** | **SRA** | **Complete %** | **Single-copy %** | **Duplicated %** | **Fragmented %** | **Missing %** | **Complete %** | **Single-copy %** | **Duplicated %** | **Fragmented %** | **Missing %** | **Complete %** | **Single-copy %** | **Duplicated %** | **Fragmented %** | **Missing %** |
| 1 | *M. musculus* | 33,540,466 | SRR10342165 | 96.7 | 69.7 | 27 | 1.8 | 1.5 | 96.9 | 69.1 | 27.8 | 1.7 | 1.4 | 96.6 | 69.5 | 27.1 | 1.7 | 1.7 |
| 2 | *M. musculus* | 43,541,512 | SRR10342166 | 97.7 | 63.2 | 34.5 | 0.9 | 1.4 | 97.8 | 63.6 | 34.2 | 0.8 | 1.4 | 97.8 | 63.6 | 34.2 | 0.8 | 1.4 |
| 3 | *M. musculus* | 43,440,008 | SRR10342167 | 97.6 | 62.9 | 34.7 | 1.2 | 1.2 | 97.8 | 63.5 | 34.3 | 1.2 | 1 | 97.5 | 63.1 | 34.4 | 1.3 | 1.2 |
| 4 | *M. musculus* | 33,595,655 | SRR9211836 | 92.3 | 79 | 13.3 | 3.8 | 3.9 | 92.4 | 78.5 | 13.9 | 3.8 | 3.8 | 92.4 | 79.1 | 13.3 | 3.9 | 3.7 |
| 5 | *M. musculus* | 33,174,108 | SRR9211837 | 91.4 | 79.8 | 11.6 | 4.2 | 4.4 | 91.5 | 79.7 | 11.8 | 4.2 | 4.3 | 91.5 | 80 | 11.5 | 4.2 | 4.3 |
| 6 | *M. musculus* | 35,030,665 | SRR9211838 | 92 | 78.9 | 13.1 | 4.3 | 3.7 | 91.8 | 78 | 13.8 | 4.4 | 3.8 | 91.8 | 78.4 | 13.4 | 4.4 | 3.8 |
| 7 | *M. musculus* | 34,877,241 | SRR9211839 | 91.4 | 79.8 | 11.6 | 4.9 | 3.7 | 91.2 | 79.7 | 11.5 | 5.1 | 3.7 | 91.3 | 80.3 | 11 | 4.9 | 3.8 |
| 1 | *C. elegans* | 24,017,399 | SRR4181026 | 74.8 | 69.7 | 5.1 | 3.1 | 22.1 | 74.7 | 69.5 | 5.2 | 3.2 | 22.1 | 74.8 | 70.0 | 4.8 | 3.1 | 22.1 |
| 2 | *C. elegans* | 24,409,279 | SRR4181032 | 75.1 | 69.8 | 5.3 | 2.4 | 22.5 | 75.2 | 69.3 | 5.9 | 2.4 | 22.4 | 75.2 | 69.6 | 5.6 | 2.4 | 22.4 |
| 3 | *C. elegans* | 24,694,386 | SRR4181033 | 75.4 | 69.4 | 6.0 | 2.7 | 21.9 | 75.3 | 69.5 | 5.8 | 2.7 | 22.0 | 75.4 | 69.7 | 5.7 | 2.7 | 21.9 |
| 4 | *C. elegans* | 26,159,433 | SRR4181034 | 74.9 | 68.2 | 6.7 | 3.0 | 22.1 | 74.7 | 68.3 | 6.4 | 3.2 | 22.1 | 74.9 | 68.0 | 6.9 | 3.0 | 22.1 |
| 5 | *C. elegans* | 26,417,070 | SRR4181036 | 75.0 | 69.4 | 5.6 | 3.2 | 21.8 | 75.2 | 69.3 | 5.9 | 3.0 | 21.8 | 75.1 | 69.0 | 6.1 | 3.1 | 21.8 |
| 6 | *C. elegans* | 24,711,402 | SRR4181037 | 75.4 | 69.9 | 5.5 | 2.5 | 22.1 | 75.2 | 69.7 | 5.5 | 2.5 | 22.3 | 75.2 | 69.9 | 5.3 | 2.5 | 22.3 |
| 1 | *D. melanogaste* | 15,648,174 | SRR6743334 | 34.6 | 28.6 | 6 | 13.1 | 52.3 | 34.4 | 28 | 6.4 | 13.4 | 52.2 | 34.2 | 28.1 | 6.1 | 13.5 | 52.3 |
| 2 | *D. melanogaste* | 15,181,806 | SRR6743335 | 32.6 | 27 | 5.6 | 12.8 | 54.6 | 32.6 | 27.1 | 5.5 | 12.5 | 54.9 | 32.8 | 27.5 | 5.3 | 12.3 | 54.9 |
| 3 | *D. melanogaste* | 9,857,308 | SRR6743336 | 26 | 22.1 | 3.9 | 13.1 | 60.9 | 25.9 | 22.1 | 3.8 | 13.4 | 60.7 | 26 | 21.8 | 4.2 | 13.2 | 60.8 |
| 4 | *D. melanogaste* | 15,940,368 | SRR6743337 | 29.8 | 24.6 | 5.2 | 9.6 | 60.6 | 29.9 | 24.2 | 5.7 | 9.5 | 60.6 | 29.7 | 24.5 | 5.2 | 9.7 | 60.6 |
| 5 | *D. melanogaste* | 18,874,930 | SRR6743338 | 36.7 | 31.8 | 4.9 | 13.1 | 50.2 | 36.7 | 31.6 | 5.1 | 12.8 | 50.5 | 36.8 | 31.3 | 5.5 | 12.7 | 50.5 |
| 6 | *D. melanogaste* | 21,169,448 | SRR6743339 | 26.3 | 20.5 | 5.8 | 10 | 63.7 | 26.8 | 20.3 | 6.5 | 9.1 | 64.1 | 26.8 | 21.5 | 5.3 | 9 | 64.2 |
| 7 | *D. melanogaste* | 16,177,860 | SRR6743340 | 33.2 | 27.5 | 5.7 | 10.9 | 55.9 | 33.4 | 27.7 | 5.7 | 10.5 | 56.1 | 33.7 | 27.6 | 6.1 | 10.1 | 56.2 |
| 8 | *D. melanogaste* | 12,436,324 | SRR6743341 | 28.9 | 24 | 4.9 | 11.2 | 59.9 | 29.9 | 25.7 | 4.2 | 10.7 | 59.4 | 29.5 | 24.7 | 4.8 | 10.9 | 59.6 |
| 9 | *D. melanogaste* | 26,395,792 | SRR6743342 | 55.7 | 47.6 | 8.1 | 11 | 33.3 | 55.5 | 46.3 | 9.2 | 11.1 | 33.4 | 55.5 | 46.3 | 9.2 | 11 | 33.5 |

| | | | | 75bp (A) 25,29,33,41 | | | | | 75bp (B) 25,31,35,43 | | | | | 75bp (C) 25,33,37,45 | | | | |
|---|---|---|---|---|---|---|---|---|---|---|---|---|---|---|---|---|---|---|
| **Nº** | **Taxa** | **Total pe-reads** | **SRA** | **Complete %** | **Single-copy %** | **Duplicated %** | **Fragmented %** | **Missing %** | **Complete %** | **Single-copy %** | **Duplicated %** | **Fragmented %** | **Missing %** | **Complete %** | **Single-copy %** | **Duplicated %** | **Fragmented %** | **Missing %** |
| 1 | *M. musculus* | 33,700,156 | SRR10067458 | 96.7 | 66.8 | 29.9 | 1.8 | 1.5 | 96.6 | 67.5 | 29.1 | 1.8 | 1.6 | 96.9 | 66.7 | 30.2 | 1.6 | 1.5 |
| 2 | *M. musculus* | 41,236,457 | SRR10067459 | 97.2 | 59.9 | 37.3 | 1.5 | 1.3 | 97.3 | 59.9 | 37.4 | 1.4 | 1.3 | 97.1 | 58.7 | 38.4 | 1.4 | 1.5 |
| 3 | *M. musculus* | 35,598,598 | SRR10067460 | 98 | 62.7 | 35.3 | 0.9 | 1.1 | 97.6 | 64 | 33.6 | 1.2 | 1.2 | 97.8 | 61.7 | 36.1 | 0.9 | 1.3 |
| 4 | *M. musculus* | 41,745,958 | SRR10067461 | 94.1 | 52.9 | 41.2 | 3.4 | 2.5 | 94.4 | 53.4 | 41 | 3.4 | 2.2 | 94.1 | 52.9 | 41.2 | 3.4 | 2.5 |
| 5 | *M. musculus* | 45,329,544 | SRR10067462 | 95.5 | 54.3 | 41.2 | 2.3 | 2.2 | 95.7 | 54.5 | 41.2 | 2.1 | 2.2 | 95.4 | 54.8 | 40.6 | 2.3 | 2.3 |
| 6 | *M. musculus* | 44,469,310 | SRR10067463 | 94.1 | 54.3 | 39.8 | 3.8 | 2.1 | 94 | 54.1 | 39.9 | 4 | 2 | 94 | 55.6 | 38.4 | 3.9 | 2.1 |
| 1 | *C. elegans* | 40,302,838 | SRR1585277 | 76.3 | 64.5 | 11.8 | 2.6 | 21.1 | 76.3 | 64.8 | 11.5 | 2.6 | 21.1 | 76.2 | 63.6 | 12.6 | 2.6 | 21.2 |
| 2 | *C. elegans* | 50,516,835 | SRR1585278 | 76.5 | 64.2 | 12.3 | 2.6 | 20.9 | 76.4 | 63.6 | 12.8 | 2.6 | 21 | 76.3 | 63.2 | 13.1 | 2.6 | 21.1 |
| 3 | *C. elegans* | 41,947,175 | SRR8257107 | 75.3 | 69.4 | 5.9 | 2.8 | 21.9 | 75.2 | 69 | 6.2 | 2.7 | 22.1 | 75.2 | 69.6 | 5.6 | 2.7 | 22.1 |
| 4 | *C. elegans* | 44,969,393 | SRR8257108 | 75 | 68.9 | 6.1 | 2.8 | 22.2 | 75.1 | 69 | 6.1 | 2.7 | 22.2 | 74.9 | 68.4 | 6.5 | 2.8 | 22.3 |
| 5 | *C. elegans* | 45,605,396 | SRR8257109 | 75.3 | 69.3 | 6 | 2.7 | 22 | 75 | 70.1 | 4.9 | 2.7 | 22.3 | 75.3 | 70.3 | 5 | 2.5 | 22.2 |
| 1 | *D. melanogaste* | 87,423,452 | SRR8559044 | 94.7 | 57.8 | 36.9 | 0.9 | 4.4 | 94.5 | 56.3 | 38.2 | 1 | 4.5 | 94.8 | 57.9 | 36.9 | 0.9 | 4.3 |
| 2 | *D. melanogaste* | 85,714,154 | SRR8559045 | 95.4 | 55.3 | 40.1 | 0.8 | 3.8 | 95.2 | 54 | 41.2 | 0.9 | 3.9 | 95.4 | 56.2 | 39.2 | 0.8 | 3.8 |
| 3 | *D. melanogaste* | 88,252,694 | SRR8559046 | 95.5 | 54.4 | 41.1 | 0.8 | 3.7 | 95.4 | 54.4 | 41 | 0.8 | 3.8 | 95.2 | 54.2 | 41 | 0.9 | 3.9 |
| 4 | *D. melanogaste* | 82,110,608 | SRR8559047 | 95.1 | 54.7 | 40.4 | 0.6 | 4.3 | 94.9 | 56.2 | 38.7 | 0.8 | 4.3 | 95.1 | 56.1 | 39 | 0.6 | 4.3 |
| 5 | *D. melanogaste* | 102,413,880 | SRR8559048 | 95.1 | 49.3 | 45.8 | 0.8 | 4.1 | 95.5 | 49.8 | 45.7 | 0.9 | 3.6 | 95.4 | 49.4 | 46 | 0.8 | 3.8 |
| 6 | *D. melanogaste* | 98,529,578 | SRR8559049 | 95.3 | 56.2 | 39.1 | 0.6 | 4.1 | 95.1 | 56.4 | 38.7 | 0.6 | 4.3 | 95.6 | 57.5 | 38.1 | 0.6 | 3.8 |

| | | | | 100bp (A) 25,35,47,61 | | | | | 100bp (B) 25,37,55,63 | | | | | 100bp (C) 25,41,57,67 | | | | |
|---|---|---|---|---|---|---|---|---|---|---|---|---|---|---|---|---|---|---|
| **Nº** | **Taxa** | **Total pe-reads** | **SRA** | **Complete %** | **Single-copy %** | **Duplicated %** | **Fragmented %** | **Missing %** | **Complete %** | **Single-copy %** | **Duplicated %** | **Fragmented %** | **Missing %** | **Complete %** | **Single-copy %** | **Duplicated %** | **Fragmented %** | **Missing %** |
| 1 | *M. musculus* | 182180123 | SRR1017092 | 97.6 | 39.1 | 58.5 | 1.2 | 1.2 | 97.9 | 37 | 60.9 | 1.4 | 0.7 | 98 | 35 | 63 | 1 | 1 |
| 2 | *M. musculus* | 107280600 | SRR1017093 | 97.8 | 38.3 | 59.5 | 1.4 | 0.8 | 98.3 | 39.2 | 59.1 | 0.6 | 1.1 | 98.3 | 38.3 | 60 | 0.5 | 1.2 |
| 3 | *M. musculus* | 127017374 | SRR1017094 | 98.3 | 36.9 | 61.4 | 0.6 | 1.1 | 98.4 | 36.8 | 61.6 | 0.6 | 1 | 98.4 | 36.5 | 61.9 | 0.8 | 0.8 |
| 5 | *M. musculus* | 102,909,165 | SRR5171077 | 34.8 | 28.7 | 6.1 | 28.9 | 36.3 | 34.7 | 28.8 | 5.9 | 29.1 | 36.2 | 34.8 | 28.8 | 6 | 28.9 | 36.3 |
| 6 | *M. musculus* | 65,372,078 | SRR5171085 | 97.8 | 41.7 | 56.1 | 1.5 | 0.7 | 97.9 | 41.6 | 56.3 | 1.5 | 0.6 | 97.6 | 42.1 | 55.5 | 1.5 | 0.9 |
| 7 | *M. musculus* | 27,016,597 | SRR5171095 | 47.9 | 36.5 | 11.4 | 27.9 | 24.2 | 47.8 | 37 | 10.8 | 28 | 24.2 | 47.8 | 36.9 | 10.9 | 28 | 24.2 |
| 1 | *C. elegans* | 49,743,412 | SRR2142254 | 72.8 | 63.6 | 9.2 | 3.5 | 23.7 | 72.9 | 63 | 9.9 | 3.5 | 23.6 | 72.7 | 63.8 | 8.9 | 3.5 | 23.8 |
| 2 | *C. elegans* | 38,836,876 | SRR2142255 | 73 | 65.7 | 7.3 | 3.4 | 23.6 | 73.2 | 65.1 | 8.1 | 3.4 | 23.4 | 73 | 65.5 | 7.5 | 3.4 | 23.6 |
| 3 | *C. elegans* | 11,166,310 | SRR2015313 | 53.6 | 42 | 11.6 | 14.5 | 31.9 | 53.8 | 42.6 | 11.2 | 14 | 32.2 | 53.7 | 41.6 | 12.1 | 14.2 | 32.1 |
| 4 | *C. elegans* | 11,626,153 | SRR2015311 | 68.4 | 54.9 | 13.5 | 6.4 | 25.2 | 68.3 | 55.1 | 13.2 | 6.7 | 25 | 68.6 | 55.7 | 12.9 | 6.3 | 25.1 |
| 5 | *C. elegans* | 12,567,755 | SRR2012785 | 75.9 | 62.5 | 13.4 | 2.5 | 21.6 | 75.8 | 62.6 | 13.2 | 2.5 | 21.7 | 75.8 | 63.1 | 12.7 | 2.6 | 21.6 |
| 6 | *C. elegans* | 14,372,593 | SRR2012781 | | | | | | | | | | | | | | | |
| 1 | *D. melanogaste* | 30,515,068 | SRR3018852 | 94.7 | 66.4 | 28.3 | 0.6 | 4.7 | 94.9 | 65.4 | 29.5 | 0.5 | 4.6 | 94.5 | 65.9 | 28.6 | 0.5 | 5 |
| 2 | *D. melanogaste* | 24,284,630 | SRR3018855 | 93.4 | 64.2 | 29.2 | 1 | 5.6 | 93.8 | 65.5 | 28.3 | 0.9 | 5.3 | 93.8 | 64 | 29.8 | 0.9 | 5.3 |
| 3 | *D. melanogaste* | 24,773,404 | SRR3018862 | 94.7 | 67.9 | 26.8 | 0.7 | 4.6 | 94.8 | 67.8 | 27 | 0.7 | 4.5 | 94.8 | 67.4 | 27.4 | 0.7 | 4.5 |
| 4 | *D. melanogaste* | 30,140,704 | SRR3019045 | 94.2 | 65.5 | 28.7 | 1 | 4.8 | 94.4 | 65 | 29.4 | 0.9 | 4.7 | 94.2 | 64.7 | 29.5 | 0.9 | 4.9 |
| 5 | *D. melanogaste* | 25,828,680 | SRR3019179 | 94.4 | 64.6 | 29.8 | 0.7 | 4.9 | 94.4 | 65.7 | 28.7 | 0.7 | 4.9 | 94.3 | 64.3 | 30 | 0.7 | 5 |
| 6 | *D. melanogaste* | 19,049,236 | SRR3019184 | 94.3 | 70.3 | 24 | 0.9 | 4.8 | 94.3 | 70.5 | 23.8 | 0.9 | 4.8 | 94.1 | 69.9 | 24.2 | 0.9 | 5 |

| | | | | 150bp (A) 25,31,41,51,65 | | | | | 150bp (B) 25,33,43,53,71 | | | | | 150bp (C) 25,35,55,75,85 | | | | |
|---|---|---|---|---|---|---|---|---|---|---|---|---|---|---|---|---|---|---|
| **Nº** | **Taxa** | **Total pe-reads** | **SRA** | **Complete %** | **Single-copy %** | **Duplicated %** | **Fragmented %** | **Missing %** | **Complete %** | **Single-copy %** | **Duplicated %** | **Fragmented %** | **Missing %** | **Complete %** | **Single-copy %** | **Duplicated %** | **Fragmented %** | **Missing %** |
| 1 | *M. musculus* | 21,173,335 | SRR10560364 | 98.1 | 50.5 | 47.6 | 1.2 | 0.7 | 97.9 | 50.1 | 47.8 | 1 | 1.1 | 98 | 51.6 | 46.4 | 1 | 1 |
| 2 | *M. musculus* | 22,731,971 | SRR10560365 | 98.1 | 46.2 | 51.9 | 0.8 | 1.1 | 98.1 | 47.6 | 50.5 | 0.8 | 1.1 | 98.1 | 47.4 | 50.7 | 0.8 | 1.1 |
| 4 | *M. musculus* | 68,712,046 | SRR8329325 | 95.5 | 42.3 | 53.2 | 3 | 1.5 | 95.8 | 41.9 | 53.9 | 2.8 | 1.4 | 96.1 | 40 | 56.1 | 2.5 | 1.4 |
| 5 | *M. musculus* | 59,012,049 | SRR8329326 | 97.2 | 36.9 | 60.3 | 1.4 | 1.4 | 97.4 | 39.3 | 58.1 | 1 | 1.6 | 97.5 | 39.8 | 57.7 | 1.3 | 1.2 |
| 6 | *M. musculus* | 21,930,527 | SRR7079069 | 96.9 | 47.6 | 49.3 | 1.7 | 1.4 | 97.1 | 44.1 | 53 | 1.4 | 1.5 | 97.1 | 44.4 | 52.7 | 1.6 | 1.3 |
| 1 | *C. elegans* | 20,345,696 | SRR10407359 | 76.6 | 65.8 | 10.8 | 2.7 | 20.7 | 76.6 | 65.6 | 11 | 2.7 | 20.7 | 76.8 | 66.6 | 10.2 | 2.6 | 20.6 |
| 2 | *C. elegans* | 19,948,493 | SRR10407358 | 77 | 64.8 | 12.2 | 2.1 | 20.9 | 76.9 | 63.9 | 13 | 2.1 | 21 | 76.9 | 65.2 | 11.7 | 2.1 | 21 |
| 3 | *C. elegans* | 21,276,093 | SRR10407357 | 76.6 | 65 | 11.6 | 2.5 | 20.9 | 76.7 | 64.4 | 12.3 | 2.6 | 20.7 | 76.5 | 65 | 11.5 | 2.5 | 21 |
| 4 | *C. elegans* | 23,993,704 | SRR10407356 | 76.3 | 66.2 | 10.1 | 2.4 | 21.3 | 76.4 | 67.2 | 9.2 | 2.3 | 21.3 | 76.5 | 66.2 | 10.3 | 2.4 | 21.1 |
| 5 | *C. elegans* | 25,110,365 | SRR10407355 | 76.4 | 65.8 | 10.6 | 2.2 | 21.4 | 76.5 | 67.4 | 9.1 | 2.3 | 21.2 | 76.3 | 66 | 10.3 | 2.2 | 21.5 |
| 6 | *C. elegans* | 22,967,010 | SRR10407354 | | | | | | | | | | | | | | | |
| 1 | *D. melanogaste* | 8,276,228 | SRR10407358 | 75.2 | 69.1 | 6.1 | 2.6 | 22.2 | 75.2 | 68.9 | 6.3 | 2.6 | 22.2 | 75.4 | 68.6 | 6.8 | 2.6 | 22 |
| 2 | *D. melanogaste* | 47,552,414 | SRR7716077 | 94.4 | 60.6 | 33.8 | 1.2 | 4.4 | 94.2 | 60.4 | 33.8 | 1.3 | 4.5 | 94.3 | 60.9 | 33.4 | 1.2 | 4.5 |
| 3 | *D. melanogaste* | 45,807,064 | SRR7716078 | 93.6 | 63 | 30.6 | 1 | 5.4 | 93.4 | 63.7 | 29.7 | 1.2 | 5.4 | 93.7 | 63.3 | 30.4 | 1.2 | 5.1 |
| 4 | *D. melanogaste* | 29,221,598 | SRR7716079 | 86.3 | 64 | 22.3 | 4.7 | 9 | 86.3 | 62.9 | 23.4 | 4.7 | 9 | 86.1 | 63.7 | 22.4 | 4.7 | 9.2 |
| 5 | *D. melanogaste* | 46,936,912 | SRR7716080 | 89.9 | 63.7 | 26.2 | 3.4 | 6.7 | 90.1 | 63.1 | 27 | 3.1 | 6.8 | 89.8 | 64.2 | 25.6 | 3.1 | 7.1 |

**50bp**

**All data by busco**

| Shapiro-Wilk normality test (p > 0.05) | | | | | | |
|---|---|---|---|---|---|---|
| | **TransPi** | **Trinity** | **Normally distributed** | **ANOVA** | **Kruskal-lis test (p < 0.05)** | **Significant** |
| Complete | 2.75E-08 | 1.79E-08 | no | - | 0.08948458 | no |
| Single-copy | 1.24E-08 | 1.84E-08 | no | - | 3.88E-05 | yes |
| Duplicated | 9.00E-06 | 1.56E-10 | no | - | 7.94E-07 | yes |
| Fragmented | 1.26E-09 | 1.50E-08 | no | - | 7.82E-06 | yes |
| Missing | 4.45E-08 | 1.13E-07 | no | - | 0.270551 | no |

**All data by kmers**

| Shapiro-Wilk normality test (p > 0.05) | | | | | | | |
|---|---|---|---|---|---|---|---|
| | **KmerA** | **KmerB** | **KmerC** | **Normally distributed** | **ANOVA** | **Kruskal-Wallis test (p < 0.05)** | **Significant** |
| Complete | 4.18E-04 | 3.97E-04 | 4.68E-04 | no | - | 0.9916716 | no |
| Single-copy | 0.0002572 | 2.43E-04 | 2.60E-04 | no | - | 9.97E-01 | no |
| Duplicated | 1.23E-02 | 1.10E-02 | 7.28E-03 | no | - | 9.61E-01 | no |
| Fragmented | 5.45E-05 | 3.48E-05 | 5.16E-05 | no | - | 0.7647052 | no |
| Missing | 5.31E-04 | 5.74E-04 | 6.30E-04 | no | - | 0.995189 | no |

***C. elegans***

| Shapiro-Wilk normality test (p > 0.05) | | | | | | | |
|---|---|---|---|---|---|---|---|
| | **KmerA** | **KmerB** | **KmerC** | **Normally distributed** | **ANOVA** | **Kruskal-Wallis test (p < 0.05)** | **Significant** |
| Complete | 9.39E-01 | 6.21E-01 | 9.15E-01 | yes | | - | no |
| Single-copy | 0.348 | 3.26E-01 | 8.57E-01 | yes | | - | no |
| Duplicated | 4.66E-01 | 6.86E-01 | 6.19E-01 | yes | | - | no |
| Fragmented | 0.2584 | 1.06E-01 | 4.78E-02 | no | - | 0.9246305 | no |
| Missing | 9.40E-01 | 7.98E-01 | 9.26E-01 | yes | | - | no |

***D. melanogaster***

| Shapiro-Wilk normality test (p > 0.05) | | | | | | | |
|---|---|---|---|---|---|---|---|
| | **KmerA** | **KmerB** | **KmerC** | **Normally distributed** | **ANOVA** | **Kruskal-Wallis test (p < 0.05)** | **Significant** |
| Complete | 4.53E-03 | 2.83E-03 | 3.58E-03 | no | - | 0.9992701 | no |
| Single-copy | 0.2013 | 1.23E-01 | 1.72E-01 | yes | | - | no |
| Duplicated | 5.53E-02 | 4.67E-02 | 3.22E-02 | no | - | 8.62E-01 | no |
| Fragmented | 0.003331 | 1.31E-02 | 8.40E-03 | no | - | 0.8590528 | no |
| Missing | 4.31E-03 | 3.00E-03 | 4.47E-03 | no | - | 0.9945684 | no |

***M. musculus***

| Shapiro-Wilk normality test (p > 0.05) | | | | | | | |
|---|---|---|---|---|---|---|---|
| | **KmerA** | **KmerB** | **KmerC** | **Normally distributed** | **ANOVA** | **Kruskal-Wallis test (p < 0.05)** | **Significant** |
| Complete | 1.58E-01 | 1.43E-01 | 1.67E-01 | yes | | - | no |
| Single-copy | 0.003992 | 5.32E-03 | 7.02E-03 | no | - | 9.49E-01 | no |
| Duplicated | 4.20E-04 | 5.65E-04 | 5.01E-04 | no | - | 9.16E-01 | no |
| Fragmented | 0.01943 | 2.37E-03 | 1.74E-02 | no | - | 0.9574024 | no |
| Missing | 1.94E-01 | 2.00E-01 | 2.85E-01 | yes | | - | no |

**75bp**

**All data by busco**

| Shapiro-Wilk normality test ($p > 0.05$) | | | | | | |
|---|---|---|---|---|---|---|
| | TransPi | Trinity | Normally distributed | ANOVA | Kruskal-lis test ($p < 0.05$) | Significant |
| Complete | 3.17E-08 | 2.23E-08 | no | - | 0.0128621 | yes |
| Single-copy | 8.48E-03 | 5.68E-05 | no | - | 3.10E-16 | yes |
| Duplicated | 4.36E-03 | 3.36E-07 | no | - | 4.21E-08 | yes |
| Fragmented | 1.70E-02 | 3.96E-06 | no | - | 0.6100149 | no |
| Missing | 5.23E-09 | 1.69E-09 | no | - | 0.004015174 | yes |

**All data by kmers**

| Shapiro-Wilk normality test ($p > 0.05$) | | | | | | | |
|---|---|---|---|---|---|---|---|
| | KmerA | KmerB | KmerC | Normally distributed | ANOVA | Kruskal-Wallis test ($p < 0.05$) | Significant |
| Complete | 3.13E-04 | 3.58E-04 | 3.95E-04 | no | - | 0.9472942 | no |
| Single-copy | 0.3689 | 2.08E-01 | 3.16E-01 | yes | | - | no |
| Duplicated | 2.54E-01 | 2.91E-01 | 1.21E-01 | yes | | - | no |
| Fragmented | 2.54E-01 | 5.70E-01 | 9.43E-02 | yes | | - | no |
| Missing | 1.26E-04 | 1.18E-04 | 1.05E-04 | no | - | 0.9927532 | no |

***C. elegans***

| Shapiro-Wilk normality test ($p > 0.05$) | | | | | | | |
|---|---|---|---|---|---|---|---|
| | KmerA | KmerB | KmerC | Normally distributed | ANOVA | Kruskal-Wallis test ($p < 0.05$) | Significant |
| Complete | 1.53E-01 | 5.49E-02 | 3.38E-01 | yes | | - | no |
| Single-copy | 0.3546 | 5.36E-01 | 3.85E-01 | yes | | - | no |
| Duplicated | 3.66E-01 | 4.23E-01 | 4.02E-01 | yes | | - | no |
| Fragmented | 0.0344 | 9.78E-01 | 8.22E-02 | no | - | 0.8758086 | no |
| Missing | 4.83E-01 | 4.60E-01 | 2.33E-01 | yes | | - | no |

***D. melanogaster***

| Shapiro-Wilk normality test ($p > 0.05$) | | | | | | | |
|---|---|---|---|---|---|---|---|
| | KmerA | KmerB | KmerC | Normally distributed | ANOVA | Kruskal-Wallis test ($p < 0.05$) | Significant |
| Complete | 3.31E-01 | 1.66E-01 | 1.71E-01 | yes | | - | no |
| Single-copy | 0.008752 | 1.15E-02 | 5.24E-03 | no | - | 9.06E-01 | no |
| Duplicated | 6.48E-03 | 9.53E-03 | 5.17E-03 | no | - | 8.19E-01 | no |
| Fragmented | 0.03849 | 6.10E-01 | 4.93E-02 | no | - | 0.5969728 | no |
| Missing | 1.71E-01 | 1.29E-01 | 7.37E-02 | yes | | - | no |

***M. musculus***

| Shapiro-Wilk normality test ($p > 0.05$) | | | | | | | |
|---|---|---|---|---|---|---|---|
| | KmerA | KmerB | KmerC | Normally distributed | ANOVA | Kruskal-Wallis test ($p < 0.05$) | Significant |
| Complete | 2.64E-02 | 4.90E-02 | 4.46E-02 | no | - | 0.9843138 | no |
| Single-copy | 0.008143 | 8.00E-03 | 6.67E-03 | no | - | 9.64E-01 | no |
| Duplicated | 5.07E-03 | 5.69E-03 | 4.04E-03 | no | - | 9.84E-01 | no |
| Fragmented | 0.08928 | 1.48E-01 | 9.83E-02 | yes | | - | no |
| Missing | 1.03E-01 | 1.51E-01 | 1.71E-01 | yes | | - | no |

100bp

**All data by busco**

| Shapiro-Wilk normality test (p > 0.05) | | | | | | |
|---|---|---|---|---|---|---|
| | **TransPi** | **Trinity** | **Normally distributed** | **ANOVA** | **Kruskal-lis test (p < 0.05)** | **Significant** |
| Complete | 4.65E-08 | 1.48E-07 | no | - | 0.1989282 | no |
| Single-copy | 9.73E-09 | 5.19E-06 | no | - | 5.97E-14 | yes |
| Duplicated | 9.20E-05 | 2.08E-05 | no | - | 3.25E-08 | yes |
| Fragmented | 2.09E-10 | 3.50E-10 | no | - | 0.1442448 | no |
| Missing | 1.59E-06 | 1.33E-06 | no | - | 0.2283286 | no |

**All data by kmers**

| Shapiro-Wilk normality test (p > 0.05) | | | | | | | |
|---|---|---|---|---|---|---|---|
| | **KmerA** | **KmerB** | **KmerC** | **Normally distributed** | **ANOVA** | **Kruskal-Wallis test (p < 0.05)** | **Significant** |
| Complete | 4.96E-04 | 4.43E-04 | 7.01E-04 | no | - | 0.9586657 | no |
| Single-copy | 0.000158 | 2.79E-04 | 2.43E-04 | no | - | 8.35E-01 | no |
| Duplicated | 3.11E-02 | 2.06E-02 | 4.29E-02 | no | - | 9.52E-01 | no |
| Fragmented | 1.29E-05 | 1.31E-05 | 2.98E-05 | no | - | 0.9505026 | no |
| Missing | 4.03E-03 | 4.10E-03 | 3.87E-03 | no | - | 0.8619683 | no |

***C. elegans***

| Shapiro-Wilk normality test (p > 0.05) | | | | | | | |
|---|---|---|---|---|---|---|---|
| | **KmerA** | **KmerB** | **KmerC** | **Normally distributed** | **ANOVA** | **Kruskal-Wallis test (p < 0.05)** | **Significant** |
| Complete | 2.73E-02 | 2.45E-02 | 2.69E-02 | no | - | 0.985216 | no |
| Single-copy | 0.4004 | 5.54E-01 | 3.71E-01 | yes | | - | no |
| Duplicated | 4.20E-01 | 1.41E-01 | 2.85E-01 | yes | | - | no |
| Fragmented | 0.01098 | 1.03E-02 | 9.51E-03 | no | - | 0.9975463 | no |
| Missing | 1.45E-01 | 1.29E-01 | 1.31E-01 | yes | | - | yes |

***D. melanogaster***

| Shapiro-Wilk normality test (p > 0.05) | | | | | | | |
|---|---|---|---|---|---|---|---|
| | **KmerA** | **KmerB** | **KmerC** | **Normally distributed** | **ANOVA** | **Kruskal-Wallis test (p < 0.05)** | **Significant** |
| Complete | 6.43E-01 | 2.64E-01 | 6.03E-01 | yes | | - | no |
| Single-copy | 0.00425 | 9.98E-03 | 9.18E-03 | no | - | 9.91E-01 | no |
| Duplicated | 6.16E-03 | 1.21E-02 | 9.19E-03 | no | - | 9.79E-01 | no |
| Fragmented | 0.9776 | 9.68E-01 | 8.77E-01 | yes | | - | no |
| Missing | 2.00E-01 | 3.87E-01 | 1.46E-01 | yes | | - | no |

***M. musculus***

| Shapiro-Wilk normality test (p > 0.05) | | | | | | | |
|---|---|---|---|---|---|---|---|
| | **KmerA** | **KmerB** | **KmerC** | **Normally distributed** | **ANOVA** | **Kruskal-Wallis test (p < 0.05)** | **Significant** |
| Complete | 1.62E-04 | 1.84E-04 | 4.94E-04 | no | - | 0.9391398 | no |
| Single-copy | 0.006094 | 8.98E-03 | 6.52E-03 | no | - | 8.80E-01 | no |
| Duplicated | 7.89E-03 | 7.46E-03 | 1.70E-02 | no | - | 9.99E-01 | no |
| Fragmented | 0.0002216 | 2.86E-04 | 1.03E-03 | no | - | 0.9874748 | no |
| Missing | 1.56E-04 | 1.52E-04 | 9.68E-04 | bo | - | 0.6108901 | no |

**150bp**

**All data by busco**

| Shapiro-Wilk normality test (p > 0.05) | | | | | | |
|---|---|---|---|---|---|---|
| | **TransPi** | **Trinity** | **Normally distributed** | **ANOVA** | **Kruskal-lis test (p < 0.05)** | **Significant** |
| Complete | 6.85E-06 | 3.70E-06 | no | - | 0.01975166 | yes |
| Single-copy | 0.001852 | 2.35E-05 | no | - | 1.82E-16 | yes |
| Duplicated | 2.39E-02 | 4.23E-05 | no | - | 8.22E-09 | yes |
| Fragmented | 0.005526 | 8.59E-05 | no | - | 0.0309718 | yes |
| Missing | 2.36E-07 | 9.30E-08 | no | - | 0.02033037 | yes |

**All data by kmers**

| Shapiro-Wilk normality test (p > 0.05) | | | | | | | |
|---|---|---|---|---|---|---|---|
| | **KmerA** | **KmerB** | **KmerC** | **Normally distributed** | **ANOVA** | **skal-Wallis test (p < 0.05)** | **Significant** |
| Complete | 9.13E-03 | 6.26E-03 | 1.12E-02 | no | - | 0.8977174 | no |
| Single-copy | 0.1016 | 2.78E-02 | 2.24E-01 | no | - | 8.76E-01 | no |
| Duplicated | 6.19E-01 | 4.48E-01 | 3.14E-01 | yes | | - | no |
| Fragmented | 0.2288 | 4.94E-01 | 1.79E-01 | yes | | - | no |
| Missing | 1.14E-03 | 9.82E-04 | 1.41E-03 | no | - | 0.8893338 | no |

***C. elegans***

| Shapiro-Wilk normality test (p > 0.05) | | | | | | | |
|---|---|---|---|---|---|---|---|
| | **KmerA** | **KmerB** | **KmerC** | **Normally distributed** | **ANOVA** | **skal-Wallis test (p < 0.05)** | **Significant** |
| Complete | 8.33E-01 | 3.86E-01 | 4.14E-01 | yes | | - | no |
| Single-copy | 0.1817 | 8.21E-02 | 1.84E-01 | yes | | - | no |
| Duplicated | 2.45E-02 | 6.72E-02 | 1.04E-01 | no | - | 9.88E-01 | no |
| Fragmented | 0.3497 | 8.97E-01 | 7.18E-01 | yes | | - | no |
| Missing | 2.21E-01 | 1.03E-01 | 2.07E-01 | yes | | - | no |

***D. melanogaster***

| Shapiro-Wilk normality test (p > 0.05) | | | | | | | |
|---|---|---|---|---|---|---|---|
| | **KmerA** | **KmerB** | **KmerC** | **Normally distributed** | **ANOVA** | **skal-Wallis test (p < 0.05)** | **Significant** |
| Complete | 5.86E-02 | 6.16E-02 | 5.82E-02 | yes | | - | no |
| Single-copy | 0.04833 | 4.84E-02 | 5.58E-02 | no | - | 9.29E-01 | no |
| Duplicated | 2.75E-01 | 2.53E-01 | 2.17E-01 | yes | | - | no |
| Fragmented | 0.8729 | 8.30E-01 | 7.73E-01 | yes | | - | no |
| Missing | 5.89E-04 | 6.52E-04 | 6.12E-04 | no | - | 0.9709572 | no |

***M. musculus***

| Shapiro-Wilk normality test (p > 0.05) | | | | | | | |
|---|---|---|---|---|---|---|---|
| | **KmerA** | **KmerB** | **KmerC** | **Normally distributed** | **ANOVA** | **skal-Wallis test (p < 0.05)** | **Significant** |
| Complete | 8.47E-01 | 2.69E-01 | 4.23E-01 | yes | | - | no |
| Single-copy | 0.01682 | 1.24E-02 | 1.36E-02 | no | - | 9.80E-01 | no |
| Duplicated | 1.56E-02 | 1.21E-02 | 1.86E-02 | no | - | 9.99E-01 | no |
| Fragmented | 0.8411 | 2.41E-01 | 2.41E-01 | yes | | - | no |
| Missing | 7.62E-06 | 4.41E-01 | 3.30E-01 | no | - | 0.7756218 | no |

Supplementary Table 3. Non-model organisms results.

| Category | Sample | Program | Score | Reads | Length | ID |
|---|---|---|---|---|---|---|
| Complete | ERR3026433_Pinnigorgia_flava | Transpi | 96.3 | 30,545,400 | 50 | A |
| Complete | ERR3026434_Sinularia_cruciata | Transpi | 91.6 | 22,160,908 | 50 | A |
| Complete | ERR3026435_Tubipora_musica | Transpi | 95.3 | 23,006,724 | 50 | A |
| Complete | ERR3040053_Heliopora_coerulea | Transpi | 94.6 | 29,000,821 | 50 | A |
| Complete | SRR8280777_Godzilliognomus_frondosus | Transpi | 87.5 | 14,086,834 | 75 | B |
| Complete | SRR8745910_Galeodes_sp | Transpi | 84.3 | 6,356,774 | 75 | B |
| Complete | SRR8745911_Peripatoides_novaezealandiae_onyc | Transpi | 84.9 | 5,768,550 | 75 | B |
| Complete | ERR1000783_Paracentrotus_lividus | Transpi | 81.7 | 6,803,316 | 75 | B |
| Complete | SRR10744002_Paracentrotus_lividus | Transpi | 94.3 | 13,583,857 | 75 | B |
| Complete | SRR1232685_Nephtys_caeca | Transpi | 29.6 | 1,576,665 | 75 | B |
| Complete | SRR1271607_Gnathostomula_paradoxa | Transpi | 56 | 5,954,962 | 75 | B |
| Complete | SRR1271706_Macrodasys_sp | Transpi | 74.7 | 3,204,609 | 75 | B |
| Complete | SRR1273732_Lepidodermella_squamata | Transpi | 54.4 | 4,370,938 | 75 | B |
| Complete | SRR1273789_Cephalothrix_linearis | Transpi | 82.6 | 4,869,244 | 75 | B |
| Complete | SRR5569439_Acropora_palmata | Transpi | 69.6 | 10,476,071 | 75 | B |
| Complete | SRR8601367_Acropora_pulchra | Transpi | 92.4 | 14,037,157 | 75 | B |
| Complete | SRR1041944_Ephydatia_muelleri | Transpi | 92 | 11,425,188 | 100 | C |
| Complete | SRR1042012_Oscarella_pearsei | Transpi | 92.6 | 11,306,242 | 100 | C |
| Complete | SRR1168575_Spongilla_lacustris | Transpi | 85 | 5,136,881 | 100 | C |
| Complete | SRR1711043_Mycale_phylophylla_adult | Transpi | 77.9 | 11,408,543 | 100 | C |
| Complete | SRR1793376_Haliclona_tubifera | Transpi | 93.5 | 16,356,602 | 100 | C |
| Complete | SRR504690_Sycon_coactum | Transpi | 83.8 | 9,098,097 | 100 | C |
| Complete | SRR1145776_Peripatopsis_capensis | Transpi | 62.4 | 11,638,180 | 100 | C |
| Complete | SRR1505119_Monodonta_labio | Transpi | 85.7 | 10,388,770 | 100 | C |
| Complete | SRR1560310_Lyonsia_floridana | Transpi | 79.6 | 9,919,645 | 100 | C |
| Complete | SRR1560359_Mercenaria_campechiensis | Transpi | 22.9 | 11,935,267 | 100 | C |
| Complete | SRR1560432_Neotrigonia_margaritacea | Transpi | 62.1 | 11,215,767 | 100 | C |
| Complete | SRR1560458_Cardites_antiquatus | Transpi | 49 | 11,916,756 | 100 | C |
| Complete | SRR1561723_Sphaerium_nucleus | Transpi | 36 | 18,539,173 | 100 | C |
| Complete | SRR1611556_Hemithiris_psittacea | Transpi | 92.8 | 9,221,875 | 100 | C |
| Complete | SRR1611560_Malacobdella_grossa | Transpi | 94 | 8,307,739 | 100 | C |
| Complete | SRR1611565_Phoronis_psammophila | Transpi | 94.1 | 12,949,999 | 100 | C |
| Complete | SRR1796434_Catenula_lemnae | Transpi | 53.5 | 3,028,636 | 100 | C |
| Complete | SRR3097584_Florometra | Transpi | 97.8 | 32,710,859 | 100 | C |
| Complete | SRR3105702_Childia_submaculatum | Transpi | 81.8 | 6,089,955 | 100 | C |
| Complete | SRR331123_Ennucula_tenuis | Transpi | 95.2 | 14,420,942 | 100 | C |
| Complete | SRR7655554_Ircinia_fasciculata | Transpi | 74.9 | 13,420,109 | 100 | C |
| Complete | SRR7754744_Krohnitta_subtilis | Transpi | 90 | 15,954,007 | 100 | C |
| Complete | SRR10527303_Octolasmis_warwickii | Transpi | 93.4 | 15,813,391 | 150 | D |
| Complete | SRR3350463_Dimya_lima | Transpi | 52.1 | 5,426,850 | 150 | D |
| Complete | SRR3404576_Brachionus_plicatilis | Transpi | 94.3 | 7,403,847 | 150 | D |
| Complete | SRR4113507_Calanus_finmarchicus | Transpi | 77.2 | 10,633,606 | 150 | D |
| Complete | SRR4294206_Millepora_alcicornis | Transpi | 96.9 | 24,645,545 | 150 | D |
| Complete | SRR504694_Corticium_candelabrum | Transpi | 91.5 | 18,897,095 | 150 | D |
| Complete | SRR5140141_Eoleptestheria_cf_ticinensis | Transpi | 83.1 | 5,471,351 | 150 | D |
| Complete | SRR5626553_Nautilus_pompilius | Transpi | 63.7 | 16,461,197 | 150 | D |
| Complete | SRR5873556_Neocalanus_Flemingeri | Transpi | 89.1 | 4,112,626 | 150 | D |
| Complete | SRR8393254_Apostichopus_japonicus | Transpi | 92 | 8,289,770 | 150 | D |
| Complete | SRR8491966_Porites_pukoensis | Transpi | 99.4 | 16,448,725 | 150 | D |
| Single-Copy | ERR3026433_Pinnigorgia_flava | Transpi | 84.7 | 30,545,400 | 50 | A |

| | | | | | | |
|---|---|---|---|---|---|---|
| Single-Copy | ERR3026434_Sinularia_cruciata | Transpi | 81.9 | 22,160,908 | 50 | A |
| Single-Copy | ERR3026435_Tubipora_musica | Transpi | 85.6 | 23,006,724 | 50 | A |
| Single-Copy | ERR3040053_Heliopora_coerulea | Transpi | 86 | 29,000,821 | 50 | A |
| Single-Copy | SRR8280777_Godzilliognomus_frondosus | Transpi | 77.7 | 14,086,834 | 75 | B |
| Single-Copy | SRR8745910_Galeodes_sp | Transpi | 76.9 | 6,356,774 | 75 | B |
| Single-Copy | SRR8745911_Peripatoides_novaezealandiae_onyc | Transpi | 80.3 | 5,768,550 | 75 | B |
| Single-Copy | ERR1000783_Paracentrotus_lividus | Transpi | 74.1 | 6,803,316 | 75 | B |
| Single-Copy | SRR10744002_Paracentrotus_lividus | Transpi | 81.6 | 13,583,857 | 75 | B |
| Single-Copy | SRR1232685_Nephtys_caeca | Transpi | 29 | 1,576,665 | 75 | B |
| Single-Copy | SRR1271607_Gnathostomula_paradoxa | Transpi | 35.8 | 5,954,962 | 75 | B |
| Single-Copy | SRR1271706_Macrodasys_sp | Transpi | 64.4 | 3,204,609 | 75 | B |
| Single-Copy | SRR1273732_Lepidodermella_squamata | Transpi | 47.4 | 4,370,938 | 75 | B |
| Single-Copy | SRR1273789_Cephalothrix_linearis | Transpi | 80.2 | 4,869,244 | 75 | B |
| Single-Copy | SRR5569439_Acropora_palmata | Transpi | 60.6 | 10,476,071 | 75 | B |
| Single-Copy | SRR8601367_Acropora_pulchra | Transpi | 80 | 14,037,157 | 75 | B |
| Single-Copy | SRR1041944_Ephydatia_muelleri | Transpi | 70.1 | 11,425,188 | 100 | C |
| Single-Copy | SRR1042012_Oscarella_pearsei | Transpi | 72.9 | 11,306,242 | 100 | C |
| Single-Copy | SRR1168575_Spongilla_lacustris | Transpi | 64.1 | 5,136,881 | 100 | C |
| Single-Copy | SRR1711043_Mycale_phylophylla_adult | Transpi | 72.9 | 11,408,543 | 100 | C |
| Single-Copy | SRR1793376_Haliclona_tubifera | Transpi | 69.8 | 16,356,602 | 100 | C |
| Single-Copy | SRR504690_Sycon_coactum | Transpi | 65.8 | 9,098,097 | 100 | C |
| Single-Copy | SRR1145776_Peripatopsis_capensis | Transpi | 59.6 | 11,638,180 | 100 | C |
| Single-Copy | SRR1505119_Monodonta_labio | Transpi | 80.2 | 10,388,770 | 100 | C |
| Single-Copy | SRR1560310_Lyonsia_floridana | Transpi | 73.1 | 9,919,645 | 100 | C |
| Single-Copy | SRR1560359_Mercenaria_campechiensis | Transpi | 19.3 | 11,935,267 | 100 | C |
| Single-Copy | SRR1560432_Neotrigonia_margaritacea | Transpi | 59.4 | 11,215,767 | 100 | C |
| Single-Copy | SRR1560458_Cardites_antiquatus | Transpi | 46.6 | 11,916,756 | 100 | C |
| Single-Copy | SRR1561723_Sphaerium_nucleus | Transpi | 27.1 | 18,539,173 | 100 | C |
| Single-Copy | SRR1611556_Hemithiris_psittacea | Transpi | 82.5 | 9,221,875 | 100 | C |
| Single-Copy | SRR1611560_Malacobdella_grossa | Transpi | 83 | 8,307,739 | 100 | C |
| Single-Copy | SRR1611565_Phoronis_psammophila | Transpi | 77.8 | 12,949,999 | 100 | C |
| Single-Copy | SRR1796434_Catenula_lemnae | Transpi | 48.9 | 3,028,636 | 100 | C |
| Single-Copy | SRR3097584_Florometra | Transpi | 87.3 | 32,710,859 | 100 | C |
| Single-Copy | SRR3105702_Childia_submaculatum | Transpi | 77.9 | 6,089,955 | 100 | C |
| Single-Copy | SRR331123_Ennucula_tenuis | Transpi | 85.3 | 14,420,942 | 100 | C |
| Single-Copy | SRR7655554_Ircinia_fasciculata | Transpi | 60.3 | 13,420,109 | 100 | C |
| Single-Copy | SRR7754744_Krohnitta_subtilis | Transpi | 60.7 | 15,954,007 | 100 | C |
| Single-Copy | SRR10527303_Octolasmis_warwickii | Transpi | 64.4 | 15,813,391 | 150 | D |
| Single-Copy | SRR3350463_Dimya_lima | Transpi | 50.4 | 5,426,850 | 150 | D |
| Single-Copy | SRR3404576_Brachionus_plicatilis | Transpi | 84.9 | 7,403,847 | 150 | D |
| Single-Copy | SRR4113507_Calanus_finmarchicus | Transpi | 62.1 | 10,633,606 | 150 | D |
| Single-Copy | SRR4294206_Millepora_alcicornis | Transpi | 83.6 | 24,645,545 | 150 | D |
| Single-Copy | SRR504694_Corticium_candelabrum | Transpi | 75.3 | 18,897,095 | 150 | D |
| Single-Copy | SRR5140141_Eoleptestheria_cf_ticinensis | Transpi | 74.2 | 5,471,351 | 150 | D |
| Single-Copy | SRR5626553_Nautilus_pompilius | Transpi | 59.5 | 16,461,197 | 150 | D |
| Single-Copy | SRR5873556_Neocalanus_Flemingeri | Transpi | 80.2 | 4,112,626 | 150 | D |
| Single-Copy | SRR8393254_Apostichopus_japonicus | Transpi | 67.4 | 8,289,770 | 150 | D |
| Single-Copy | SRR8491966_Porites_pukoensis | Transpi | 38.2 | 16,448,725 | 150 | D |
| Duplicated | ERR3026433_Pinnigorgia_flava | Transpi | 11.6 | 30,545,400 | 50 | A |
| Duplicated | ERR3026434_Sinularia_cruciata | Transpi | 9.7 | 22,160,908 | 50 | A |
| Duplicated | ERR3026435_Tubipora_musica | Transpi | 9.7 | 23,006,724 | 50 | A |

| | | | | | | |
|---|---|---|---|---|---|---|
| Duplicated | ERR3040053_Heliopora_coerulea | Transpi | 8.6 | 29,000,821 | 50 | A |
| Duplicated | SRR8280777_Godzilliognomus_frondosus | Transpi | 9.8 | 14,086,834 | 75 | B |
| Duplicated | SRR8745910_Galeodes_sp | Transpi | 7.4 | 6,356,774 | 75 | B |
| Duplicated | SRR8745911_Peripatoides_novaezealandiae_onyc | Transpi | 4.6 | 5,768,550 | 75 | B |
| Duplicated | ERR1000783_Paracentrotus_lividus | Transpi | 7.6 | 6,803,316 | 75 | B |
| Duplicated | SRR10744002_Paracentrotus_lividus | Transpi | 12.7 | 13,583,857 | 75 | B |
| Duplicated | SRR1232685_Nephtys_caeca | Transpi | 0.6 | 1,576,665 | 75 | B |
| Duplicated | SRR1271607_Gnathostomula_paradoxa | Transpi | 20.2 | 5,954,962 | 75 | B |
| Duplicated | SRR1271706_Macrodasys_sp | Transpi | 10.3 | 3,204,609 | 75 | B |
| Duplicated | SRR1273732_Lepidodermella_squamata | Transpi | 7 | 4,370,938 | 75 | B |
| Duplicated | SRR1273789_Cephalothrix_linearis | Transpi | 2.4 | 4,869,244 | 75 | B |
| Duplicated | SRR5569439_Acropora_palmata | Transpi | 9 | 10,476,071 | 75 | B |
| Duplicated | SRR8601367_Acropora_pulchra | Transpi | 12.4 | 14,037,157 | 75 | B |
| Duplicated | SRR1041944_Ephydatia_muelleri | Transpi | 21.9 | 11,425,188 | 100 | C |
| Duplicated | SRR1042012_Oscarella_pearsei | Transpi | 19.7 | 11,306,242 | 100 | C |
| Duplicated | SRR1168575_Spongilla_lacustris | Transpi | 20.9 | 5,136,881 | 100 | C |
| Duplicated | SRR1711043_Mycale_phylophylla_adult | Transpi | 5 | 11,408,543 | 100 | C |
| Duplicated | SRR1793376_Haliclona_tubifera | Transpi | 23.7 | 16,356,602 | 100 | C |
| Duplicated | SRR504690_Sycon_coactum | Transpi | 18 | 9,098,097 | 100 | C |
| Duplicated | SRR1145776_Peripatopsis_capensis | Transpi | 2.8 | 11,638,180 | 100 | C |
| Duplicated | SRR1505119_Monodonta_labio | Transpi | 5.5 | 10,388,770 | 100 | C |
| Duplicated | SRR1560310_Lyonsia_floridana | Transpi | 6.5 | 9,919,645 | 100 | C |
| Duplicated | SRR1560359_Mercenaria_campechiensis | Transpi | 3.6 | 11,935,267 | 100 | C |
| Duplicated | SRR1560432_Neotrigonia_margaritacea | Transpi | 2.7 | 11,215,767 | 100 | C |
| Duplicated | SRR1560458_Cardites_antiquatus | Transpi | 2.4 | 11,916,756 | 100 | C |
| Duplicated | SRR1561723_Sphaerium_nucleus | Transpi | 8.9 | 18,539,173 | 100 | C |
| Duplicated | SRR1611556_Hemithiris_psittacea | Transpi | 10.3 | 9,221,875 | 100 | C |
| Duplicated | SRR1611560_Malacobdella_grossa | Transpi | 11 | 8,307,739 | 100 | C |
| Duplicated | SRR1611565_Phoronis_psammophila | Transpi | 16.3 | 12,949,999 | 100 | C |
| Duplicated | SRR1796434_Catenula_lemnae | Transpi | 4.6 | 3,028,636 | 100 | C |
| Duplicated | SRR3097584_Florometra | Transpi | 10.5 | 32,710,859 | 100 | C |
| Duplicated | SRR3105702_Childia_submaculatum | Transpi | 3.9 | 6,089,955 | 100 | C |
| Duplicated | SRR331123_Ennucula_tenuis | Transpi | 9.9 | 14,420,942 | 100 | C |
| Duplicated | SRR7655554_Ircinia_fasciculata | Transpi | 14.6 | 13,420,109 | 100 | C |
| Duplicated | SRR7754744_Krohnitta_subtilis | Transpi | 29.3 | 15,954,007 | 100 | C |
| Duplicated | SRR10527303_Octolasmis_warwickii | Transpi | 29 | 15,813,391 | 150 | D |
| Duplicated | SRR3350463_Dimya_lima | Transpi | 1.7 | 5,426,850 | 150 | D |
| Duplicated | SRR3404576_Brachionus_plicatilis | Transpi | 9.4 | 7,403,847 | 150 | D |
| Duplicated | SRR4113507_Calanus_finmarchicus | Transpi | 15.1 | 10,633,606 | 150 | D |
| Duplicated | SRR4294206_Millepora_alcicornis | Transpi | 13.3 | 24,645,545 | 150 | D |
| Duplicated | SRR504694_Corticium_candelabrum | Transpi | 16.2 | 18,897,095 | 150 | D |
| Duplicated | SRR5140141_Eoleptestheria_cf_ticinensis | Transpi | 8.9 | 5,471,351 | 150 | D |
| Duplicated | SRR5626553_Nautilus_pompilius | Transpi | 4.2 | 16,461,197 | 150 | D |
| Duplicated | SRR5873556_Neocalanus_Flemingeri | Transpi | 8.9 | 4,112,626 | 150 | D |
| Duplicated | SRR8393254_Apostichopus_japonicus | Transpi | 24.6 | 8,289,770 | 150 | D |
| Duplicated | SRR8491966_Porites_pukoensis | Transpi | 61.2 | 16,448,725 | 150 | D |
| Fragmented | ERR3026433_Pinnigorgia_flava | Transpi | 1.7 | 30,545,400 | 50 | A |
| Fragmented | ERR3026434_Sinularia_cruciata | Transpi | 5.3 | 22,160,908 | 50 | A |
| Fragmented | ERR3026435_Tubipora_musica | Transpi | 2.2 | 23,006,724 | 50 | A |
| Fragmented | ERR3040053_Heliopora_coerulea | Transpi | 3.8 | 29,000,821 | 50 | A |
| Fragmented | SRR8280777_Godzilliognomus_frondosus | Transpi | 5.4 | 14,086,834 | 75 | B |

| | | | | | | |
|---|---|---|---|---|---|---|
| Fragmented | SRR8745910_Galeodes_sp | Transpi | 11.3 | 6,356,774 | 75 | B |
| Fragmented | SRR8745911_Peripatoides_novaezealandiae_onyc | Transpi | 6.4 | 5,768,550 | 75 | B |
| Fragmented | ERR1000783_Paracentrotus_lividus | Transpi | 9.8 | 6,803,316 | 75 | B |
| Fragmented | SRR10744002_Paracentrotus_lividus | Transpi | 3.2 | 13,583,857 | 75 | B |
| Fragmented | SRR1232685_Nephtys_caeca | Transpi | 17.3 | 1,576,665 | 75 | B |
| Fragmented | SRR1271607_Gnathostomula_paradoxa | Transpi | 22 | 5,954,962 | 75 | B |
| Fragmented | SRR1271706_Macrodasys_sp | Transpi | 12.2 | 3,204,609 | 75 | B |
| Fragmented | SRR1273732_Lepidodermella_squamata | Transpi | 20.7 | 4,370,938 | 75 | B |
| Fragmented | SRR1273789_Cephalothrix_linearis | Transpi | 10.8 | 4,869,244 | 75 | B |
| Fragmented | SRR5569439_Acropora_palmata | Transpi | 17.9 | 10,476,071 | 75 | B |
| Fragmented | SRR8601367_Acropora_pulchra | Transpi | 5.5 | 14,037,157 | 75 | B |
| Fragmented | SRR1041944_Ephydatia_muelleri | Transpi | 2.7 | 11,425,188 | 100 | C |
| Fragmented | SRR1042012_Oscarella_pearsei | Transpi | 2.1 | 11,306,242 | 100 | C |
| Fragmented | SRR1168575_Spongilla_lacustris | Transpi | 9 | 5,136,881 | 100 | C |
| Fragmented | SRR1711043_Mycale_phylophylla_adult | Transpi | 11 | 11,408,543 | 100 | C |
| Fragmented | SRR1793376_Haliclona_tubifera | Transpi | 1.4 | 16,356,602 | 100 | C |
| Fragmented | SRR504690_Sycon_coactum | Transpi | 7 | 9,098,097 | 100 | C |
| Fragmented | SRR1145776_Peripatopsis_capensis | Transpi | 22.2 | 11,638,180 | 100 | C |
| Fragmented | SRR1505119_Monodonta_labio | Transpi | 8.9 | 10,388,770 | 100 | C |
| Fragmented | SRR1560310_Lyonsia_floridana | Transpi | 14.5 | 9,919,645 | 100 | C |
| Fragmented | SRR1560359_Mercenaria_campechiensis | Transpi | 18 | 11,935,267 | 100 | C |
| Fragmented | SRR1560432_Neotrigonia_margaritacea | Transpi | 19.8 | 11,215,767 | 100 | C |
| Fragmented | SRR1560458_Cardites_antiquatus | Transpi | 23.1 | 11,916,756 | 100 | C |
| Fragmented | SRR1561723_Sphaerium_nucleus | Transpi | 32.1 | 18,539,173 | 100 | C |
| Fragmented | SRR1611556_Hemithiris_psittacea | Transpi | 3.1 | 9,221,875 | 100 | C |
| Fragmented | SRR1611560_Malacobdella_grossa | Transpi | 2.4 | 8,307,739 | 100 | C |
| Fragmented | SRR1611565_Phoronis_psammophila | Transpi | 4.5 | 12,949,999 | 100 | C |
| Fragmented | SRR1796434_Catenula_lemnae | Transpi | 16.3 | 3,028,636 | 100 | C |
| Fragmented | SRR3097584_Florometra | Transpi | 0.6 | 32,710,859 | 100 | C |
| Fragmented | SRR3105702_Childia_submaculatum | Transpi | 3.4 | 6,089,955 | 100 | C |
| Fragmented | SRR331123_Ennucula_tenuis | Transpi | 3.2 | 14,420,942 | 100 | C |
| Fragmented | SRR7655554_Ircinia_fasciculata | Transpi | 15 | 13,420,109 | 100 | C |
| Fragmented | SRR7754744_Krohnitta_subtilis | Transpi | 6.2 | 15,954,007 | 100 | C |
| Fragmented | SRR10527303_Octolasmis_warwickii | Transpi | 2.7 | 15,813,391 | 150 | D |
| Fragmented | SRR3350463_Dimya_lima | Transpi | 22.5 | 5,426,850 | 150 | D |
| Fragmented | SRR3404576_Brachionus_plicatilis | Transpi | 0.8 | 7,403,847 | 150 | D |
| Fragmented | SRR4113507_Calanus_finmarchicus | Transpi | 13.3 | 10,633,606 | 150 | D |
| Fragmented | SRR4294206_Millepora_alcicornis | Transpi | 1 | 24,645,545 | 150 | D |
| Fragmented | SRR504694_Corticium_candelabrum | Transpi | 4.6 | 18,897,095 | 150 | D |
| Fragmented | SRR5140141_Eoleptestheria_cf_ticinensis | Transpi | 11.1 | 5,471,351 | 150 | D |
| Fragmented | SRR5626553_Nautilus_pompilius | Transpi | 25.3 | 16,461,197 | 150 | D |
| Fragmented | SRR5873556_Neocalanus_Flemingeri | Transpi | 6 | 4,112,626 | 150 | D |
| Fragmented | SRR8393254_Apostichopus_japonicus | Transpi | 6 | 8,289,770 | 150 | D |
| Fragmented | SRR8491966_Porites_pukoensis | Transpi | 0.5 | 16,448,725 | 150 | D |
| Missing | ERR3026433_Pinnigorgia_flava | Transpi | 2 | 30,545,400 | 50 | A |
| Missing | ERR3026434_Sinularia_cruciata | Transpi | 3.1 | 22,160,908 | 50 | A |
| Missing | ERR3026435_Tubipora_musica | Transpi | 2.5 | 23,006,724 | 50 | A |
| Missing | ERR3040053_Heliopora_coerulea | Transpi | 1.6 | 29,000,821 | 50 | A |
| Missing | SRR8280777_Godzilliognomus_frondosus | Transpi | 7.1 | 14,086,834 | 75 | B |
| Missing | SRR8745910_Galeodes_sp | Transpi | 4.4 | 6,356,774 | 75 | B |
| Missing | SRR8745911_Peripatoides_novaezealandiae_onyc | Transpi | 8.7 | 5,768,550 | 75 | B |

| | | | | | | |
|---|---|---|---|---|---|---|
| Missing | ERR1000783_Paracentrotus_lividus | Transpi | 8.5 | 6,803,316 | 75 | B |
| Missing | SRR10744002_Paracentrotus_lividus | Transpi | 2.5 | 13,583,857 | 75 | B |
| Missing | SRR1232685_Nephtys_caeca | Transpi | 53.1 | 1,576,665 | 75 | B |
| Missing | SRR1271607_Gnathostomula_paradoxa | Transpi | 22 | 5,954,962 | 75 | B |
| Missing | SRR1271706_Macrodasys_sp | Transpi | 13.1 | 3,204,609 | 75 | B |
| Missing | SRR1273732_Lepidodermella_squamata | Transpi | 24.9 | 4,370,938 | 75 | B |
| Missing | SRR1273789_Cephalothrix_linearis | Transpi | 6.6 | 4,869,244 | 75 | B |
| Missing | SRR5569439_Acropora_palmata | Transpi | 12.5 | 10,476,071 | 75 | B |
| Missing | SRR8601367_Acropora_pulchra | Transpi | 2.1 | 14,037,157 | 75 | B |
| Missing | SRR1041944_Ephydatia_muelleri | Transpi | 5.3 | 11,425,188 | 100 | C |
| Missing | SRR1042012_Oscarella_pearsei | Transpi | 5.3 | 11,306,242 | 100 | C |
| Missing | SRR1168575_Spongilla_lacustris | Transpi | 6 | 5,136,881 | 100 | C |
| Missing | SRR1711043_Mycale_phylophylla_adult | Transpi | 11.1 | 11,408,543 | 100 | C |
| Missing | SRR1793376_Haliclona_tubifera | Transpi | 5.1 | 16,356,602 | 100 | C |
| Missing | SRR504690_Sycon_coactum | Transpi | 9.2 | 9,098,097 | 100 | C |
| Missing | SRR1145776_Peripatopsis_capensis | Transpi | 15.4 | 11,638,180 | 100 | C |
| Missing | SRR1505119_Monodonta_labio | Transpi | 5.4 | 10,388,770 | 100 | C |
| Missing | SRR1560310_Lyonsia_floridana | Transpi | 5.9 | 9,919,645 | 100 | C |
| Missing | SRR1560359_Mercenaria_campechiensis | Transpi | 59.1 | 11,935,267 | 100 | C |
| Missing | SRR1560432_Neotrigonia_margaritacea | Transpi | 18.1 | 11,215,767 | 100 | C |
| Missing | SRR1560458_Cardites_antiquatus | Transpi | 27.9 | 11,916,756 | 100 | C |
| Missing | SRR1561723_Sphaerium_nucleus | Transpi | 31.9 | 18,539,173 | 100 | C |
| Missing | SRR1611556_Hemithiris_psittacea | Transpi | 4.1 | 9,221,875 | 100 | C |
| Missing | SRR1611560_Malacobdella_grossa | Transpi | 3.6 | 8,307,739 | 100 | C |
| Missing | SRR1611565_Phoronis_psammophila | Transpi | 1.4 | 12,949,999 | 100 | C |
| Missing | SRR1796434_Catenula_lemnae | Transpi | 30.2 | 3,028,636 | 100 | C |
| Missing | SRR3097584_Florometra | Transpi | 1.6 | 32,710,859 | 100 | C |
| Missing | SRR3105702_Childia_submaculatum | Transpi | 14.8 | 6,089,955 | 100 | C |
| Missing | SRR331123_Ennucula_tenuis | Transpi | 1.6 | 14,420,942 | 100 | C |
| Missing | SRR7655554_Ircinia_fasciculata | Transpi | 10.1 | 13,420,109 | 100 | C |
| Missing | SRR7754744_Krohnitta_subtilis | Transpi | 3.8 | 15,954,007 | 100 | C |
| Missing | SRR10527303_Octolasmis_warwickii | Transpi | 3.9 | 15,813,391 | 150 | D |
| Missing | SRR3350463_Dimya_lima | Transpi | 25.4 | 5,426,850 | 150 | D |
| Missing | SRR3404576_Brachionus_plicatilis | Transpi | 4.9 | 7,403,847 | 150 | D |
| Missing | SRR4113507_Calanus_finmarchicus | Transpi | 9.5 | 10,633,606 | 150 | D |
| Missing | SRR4294206_Millepora_alcicornis | Transpi | 2.1 | 24,645,545 | 150 | D |
| Missing | SRR504694_Corticium_candelabrum | Transpi | 3.9 | 18,897,095 | 150 | D |
| Missing | SRR5140141_Eoleptestheria_cf_ticinensis | Transpi | 5.8 | 5,471,351 | 150 | D |
| Missing | SRR5626553_Nautilus_pompilius | Transpi | 11 | 16,461,197 | 150 | D |
| Missing | SRR5873556_Neocalanus_Flemingeri | Transpi | 4.9 | 4,112,626 | 150 | D |
| Missing | SRR8393254_Apostichopus_japonicus | Transpi | 2 | 8,289,770 | 150 | D |
| Missing | SRR8491966_Porites_pukoensis | Transpi | 0.1 | 16,448,725 | 150 | D |
| Complete | ERR3026433_Pinnigorgia_flava | Trinity | 93.6 | 30,545,400 | 50 | A |
| Complete | ERR3026434_Sinularia_cruciata | Trinity | 84.9 | 22,160,908 | 50 | A |
| Complete | ERR3026435_Tubipora_musica | Trinity | 91.6 | 23,006,724 | 50 | A |
| Complete | ERR3040053_Heliopora_coerulea | Trinity | 89.5 | 29,000,821 | 50 | A |
| Complete | SRR8280777_Godzilliognomus_frondosus | Trinity | 82.2 | 14,086,834 | 75 | B |
| Complete | SRR8745910_Galeodes_sp | Trinity | 82.8 | 6,356,774 | 75 | B |
| Complete | SRR8745911_Peripatoides_novaezealandiae_onyc | Trinity | 83.3 | 5,768,550 | 75 | B |
| Complete | ERR1000783_Paracentrotus_lividus | Trinity | 73.9 | 6,803,316 | 75 | B |
| Complete | SRR10744002_Paracentrotus_lividus | Trinity | 93.4 | 13,583,857 | 75 | B |

| | | | | | | |
|---|---|---|---|---|---|---|
| Complete | SRR1232685_Nephtys_caeca | Trinity | 26.6 | 1,576,665 | 75 | B |
| Complete | SRR1271607_Gnathostomula_paradoxa | Trinity | 54.1 | 5,954,962 | 75 | B |
| Complete | SRR1271706_Macrodasys_sp | Trinity | 72.6 | 3,204,609 | 75 | B |
| Complete | SRR1273732_Lepidodermella_squamata | Trinity | 52.1 | 4,370,938 | 75 | B |
| Complete | SRR1273789_Cephalothrix_linearis | Trinity | 79.7 | 4,869,244 | 75 | B |
| Complete | SRR5569439_Acropora_palmata | Trinity | 66.1 | 10,476,071 | 75 | B |
| Complete | SRR8601367_Acropora_pulchra | Trinity | 89.8 | 14,037,157 | 75 | B |
| Complete | SRR1041944_Ephydatia_muelleri | Trinity | 94.4 | 11,425,188 | 100 | C |
| Complete | SRR1042012_Oscarella_pearsei | Trinity | 92.6 | 11,306,242 | 100 | C |
| Complete | SRR1168575_Spongilla_lacustris | Trinity | 85 | 5,136,881 | 100 | C |
| Complete | SRR1711043_Mycale_phylophylla_adult | Trinity | 77.4 | 11,408,543 | 100 | C |
| Complete | SRR1793376_Haliclona_tubifera | Trinity | 94.2 | 16,356,602 | 100 | C |
| Complete | SRR504690_Sycon_coactum | Trinity | 82.6 | 9,098,097 | 100 | C |
| Complete | SRR1145776_Peripatopsis_capensis | Trinity | 61.6 | 11,638,180 | 100 | C |
| Complete | SRR1505119_Monodonta_labio | Trinity | 84.2 | 10,388,770 | 100 | C |
| Complete | SRR1560310_Lyonsia_floridana | Trinity | 76.7 | 9,919,645 | 100 | C |
| Complete | SRR1560359_Mercenaria_campechiensis | Trinity | 20.3 | 11,935,267 | 100 | C |
| Complete | SRR1560432_Neotrigonia_margaritacea | Trinity | 57.7 | 11,215,767 | 100 | C |
| Complete | SRR1560458_Cardites_antiquatus | Trinity | 42.7 | 11,916,756 | 100 | C |
| Complete | SRR1561723_Sphaerium_nucleus | Trinity | 34.6 | 18,539,173 | 100 | C |
| Complete | SRR1611556_Hemithiris_psittacea | Trinity | 94.5 | 9,221,875 | 100 | C |
| Complete | SRR1611560_Malacobdella_grossa | Trinity | 94.3 | 8,307,739 | 100 | C |
| Complete | SRR1611565_Phoronis_psammophila | Trinity | 92.6 | 12,949,999 | 100 | C |
| Complete | SRR1796434_Catenula_lemnae | Trinity | 52.5 | 3,028,636 | 100 | C |
| Complete | SRR3097584_Florometra | Trinity | 98 | 32,710,859 | 100 | C |
| Complete | SRR3105702_Childia_submaculatum | Trinity | 81.8 | 6,089,955 | 100 | C |
| Complete | SRR331123_Ennucula_tenuis | Trinity | 94.2 | 14,420,942 | 100 | C |
| Complete | SRR7655554_Ircinia_fasciculata | Trinity | 71.4 | 13,420,109 | 100 | C |
| Complete | SRR7754744_Krohnitta_subtilis | Trinity | 88.2 | 15,954,007 | 100 | C |
| Complete | SRR10527303_Octolasmis_warwickii | Trinity | 97.5 | 15,813,391 | 150 | D |
| Complete | SRR3350463_Dimya_lima | Trinity | 53.1 | 5,426,850 | 150 | D |
| Complete | SRR3404576_Brachionus_plicatilis | Trinity | 95.5 | 7,403,847 | 150 | D |
| Complete | SRR4113507_Calanus_finmarchicus | Trinity | 79.8 | 10,633,606 | 150 | D |
| Complete | SRR4294206_Millepora_alcicornis | Trinity | 97.3 | 24,645,545 | 150 | D |
| Complete | SRR504694_Corticium_candelabrum | Trinity | 91.2 | 18,897,095 | 150 | D |
| Complete | SRR5140141_Eoleptestheria_cf_ticinensis | Trinity | 84.8 | 5,471,351 | 150 | D |
| Complete | SRR5626553_Nautilus_pompilius | Trinity | 62.2 | 16,461,197 | 150 | D |
| Complete | SRR5873556_Neocalanus_Flemingeri | Trinity | 88.8 | 4,112,626 | 150 | D |
| Complete | SRR8393254_Apostichopus_japonicus | Trinity | 91.6 | 8,289,770 | 150 | D |
| Complete | SRR8491966_Porites_pukoensis | Trinity | 99.4 | 16,448,725 | 150 | D |
| Single-Copy | ERR3026433_Pinnigorgia_flava | Trinity | 66.6 | 30,545,400 | 50 | A |
| Single-Copy | ERR3026434_Sinularia_cruciata | Trinity | 68 | 22,160,908 | 50 | A |
| Single-Copy | ERR3026435_Tubipora_musica | Trinity | 73.5 | 23,006,724 | 50 | A |
| Single-Copy | ERR3040053_Heliopora_coerulea | Trinity | 65.7 | 29,000,821 | 50 | A |
| Single-Copy | SRR8280777_Godzilliognomus_frondosus | Trinity | 63.6 | 14,086,834 | 75 | B |
| Single-Copy | SRR8745910_Galeodes_sp | Trinity | 51 | 6,356,774 | 75 | B |
| Single-Copy | SRR8745911_Peripatoides_novaezealandiae_onyc | Trinity | 66.7 | 5,768,550 | 75 | B |
| Single-Copy | ERR1000783_Paracentrotus_lividus | Trinity | 40.8 | 6,803,316 | 75 | B |
| Single-Copy | SRR10744002_Paracentrotus_lividus | Trinity | 30 | 13,583,857 | 75 | B |
| Single-Copy | SRR1232685_Nephtys_caeca | Trinity | 23.9 | 1,576,665 | 75 | B |
| Single-Copy | SRR1271607_Gnathostomula_paradoxa | Trinity | 28.1 | 5,954,962 | 75 | B |

| | | | | | | |
|---|---|---|---|---|---|---|
| Single-Copy | SRR1271706_Macrodasys_sp | Trinity | 56.5 | 3,204,609 | 75 | B |
| Single-Copy | SRR1273732_Lepidodermella_squamata | Trinity | 40.4 | 4,370,938 | 75 | B |
| Single-Copy | SRR1273789_Cephalothrix_linearis | Trinity | 65.7 | 4,869,244 | 75 | B |
| Single-Copy | SRR5569439_Acropora_palmata | Trinity | 46.2 | 10,476,071 | 75 | B |
| Single-Copy | SRR8601367_Acropora_pulchra | Trinity | 58.7 | 14,037,157 | 75 | B |
| Single-Copy | SRR1041944_Ephydatia_muelleri | Trinity | 33.3 | 11,425,188 | 100 | C |
| Single-Copy | SRR1042012_Oscarella_pearsei | Trinity | 32.7 | 11,306,242 | 100 | C |
| Single-Copy | SRR1168575_Spongilla_lacustris | Trinity | 41.7 | 5,136,881 | 100 | C |
| Single-Copy | SRR1711043_Mycale_phylophylla_adult | Trinity | 47.8 | 11,408,543 | 100 | C |
| Single-Copy | SRR1793376_Haliclona_tubifera | Trinity | 27.8 | 16,356,602 | 100 | C |
| Single-Copy | SRR504690_Sycon_coactum | Trinity | 47 | 9,098,097 | 100 | C |
| Single-Copy | SRR1145776_Peripatopsis_capensis | Trinity | 48.7 | 11,638,180 | 100 | C |
| Single-Copy | SRR1505119_Monodonta_labio | Trinity | 58.9 | 10,388,770 | 100 | C |
| Single-Copy | SRR1560310_Lyonsia_floridana | Trinity | 52.4 | 9,919,645 | 100 | C |
| Single-Copy | SRR1560359_Mercenaria_campechiensis | Trinity | 13.7 | 11,935,267 | 100 | C |
| Single-Copy | SRR1560432_Neotrigonia_margaritacea | Trinity | 47.3 | 11,215,767 | 100 | C |
| Single-Copy | SRR1560458_Cardites_antiquatus | Trinity | 32.3 | 11,916,756 | 100 | C |
| Single-Copy | SRR1561723_Sphaerium_nucleus | Trinity | 16.4 | 18,539,173 | 100 | C |
| Single-Copy | SRR1611556_Hemithiris_psittacea | Trinity | 29.1 | 9,221,875 | 100 | C |
| Single-Copy | SRR1611560_Malacobdella_grossa | Trinity | 20.4 | 8,307,739 | 100 | C |
| Single-Copy | SRR1611565_Phoronis_psammophila | Trinity | 21.6 | 12,949,999 | 100 | C |
| Single-Copy | SRR1796434_Catenula_lemnae | Trinity | 31.7 | 3,028,636 | 100 | C |
| Single-Copy | SRR3097584_Florometra | Trinity | 41.3 | 32,710,859 | 100 | C |
| Single-Copy | SRR3105702_Childia_submaculatum | Trinity | 25.9 | 6,089,955 | 100 | C |
| Single-Copy | SRR331123_Ennucula_tenuis | Trinity | 53.8 | 14,420,942 | 100 | C |
| Single-Copy | SRR7655554_Ircinia_fasciculata | Trinity | 35.9 | 13,420,109 | 100 | C |
| Single-Copy | SRR7754744_Krohnitta_subtilis | Trinity | 32.2 | 15,954,007 | 100 | C |
| Single-Copy | SRR10527303_Octolasmis_warwickii | Trinity | 27.6 | 15,813,391 | 150 | D |
| Single-Copy | SRR3350463_Dimya_lima | Trinity | 34 | 5,426,850 | 150 | D |
| Single-Copy | SRR3404576_Brachionus_plicatilis | Trinity | 57.7 | 7,403,847 | 150 | D |
| Single-Copy | SRR4113507_Calanus_finmarchicus | Trinity | 31.7 | 10,633,606 | 150 | D |
| Single-Copy | SRR4294206_Millepora_alcicornis | Trinity | 48.7 | 24,645,545 | 150 | D |
| Single-Copy | SRR504694_Corticium_candelabrum | Trinity | 33.7 | 18,897,095 | 150 | D |
| Single-Copy | SRR5140141_Eoleptestheria_cf_ticinensis | Trinity | 44.5 | 5,471,351 | 150 | D |
| Single-Copy | SRR5626553_Nautilus_pompilius | Trinity | 34.4 | 16,461,197 | 150 | D |
| Single-Copy | SRR5873556_Neocalanus_Flemingeri | Trinity | 60.2 | 4,112,626 | 150 | D |
| Single-Copy | SRR8393254_Apostichopus_japonicus | Trinity | 26.2 | 8,289,770 | 150 | D |
| Single-Copy | SRR8491966_Porites_pukoensis | Trinity | 23.5 | 16,448,725 | 150 | D |
| Duplicated | ERR3026433_Pinnigorgia_flava | Trinity | 27 | 30,545,400 | 50 | A |
| Duplicated | ERR3026434_Sinularia_cruciata | Trinity | 16.9 | 22,160,908 | 50 | A |
| Duplicated | ERR3026435_Tubipora_musica | Trinity | 18.1 | 23,006,724 | 50 | A |
| Duplicated | ERR3040053_Heliopora_coerulea | Trinity | 23.8 | 29,000,821 | 50 | A |
| Duplicated | SRR8280777_Godzilliognomus_frondosus | Trinity | 18.6 | 14,086,834 | 75 | B |
| Duplicated | SRR8745910_Galeodes_sp | Trinity | 31.8 | 6,356,774 | 75 | B |
| Duplicated | SRR8745911_Peripatoides_novaezealandiae_onyc | Trinity | 16.6 | 5,768,550 | 75 | B |
| Duplicated | ERR1000783_Paracentrotus_lividus | Trinity | 33.1 | 6,803,316 | 75 | B |
| Duplicated | SRR10744002_Paracentrotus_lividus | Trinity | 63.4 | 13,583,857 | 75 | B |
| Duplicated | SRR1232685_Nephtys_caeca | Trinity | 2.7 | 1,576,665 | 75 | B |
| Duplicated | SRR1271607_Gnathostomula_paradoxa | Trinity | 26 | 5,954,962 | 75 | B |
| Duplicated | SRR1271706_Macrodasys_sp | Trinity | 16.1 | 3,204,609 | 75 | B |
| Duplicated | SRR1273732_Lepidodermella_squamata | Trinity | 11.7 | 4,370,938 | 75 | B |

| | | | | | | |
|---|---|---|---|---|---|---|
| Duplicated | SRR1273789_Cephalothrix_linearis | Trinity | 14 | 4,869,244 | 75 | B |
| Duplicated | SRR5569439_Acropora_palmata | Trinity | 19.9 | 10,476,071 | 75 | B |
| Duplicated | SRR8601367_Acropora_pulchra | Trinity | 31.1 | 14,037,157 | 75 | B |
| Duplicated | SRR1041944_Ephydatia_muelleri | Trinity | 61.1 | 11,425,188 | 100 | C |
| Duplicated | SRR1042012_Oscarella_pearsei | Trinity | 59.9 | 11,306,242 | 100 | C |
| Duplicated | SRR1168575_Spongilla_lacustris | Trinity | 43.3 | 5,136,881 | 100 | C |
| Duplicated | SRR1711043_Mycale_phylophylla_adult | Trinity | 29.6 | 11,408,543 | 100 | C |
| Duplicated | SRR1793376_Haliclona_tubifera | Trinity | 66.4 | 16,356,602 | 100 | C |
| Duplicated | SRR504690_Sycon_coactum | Trinity | 35.6 | 9,098,097 | 100 | C |
| Duplicated | SRR1145776_Peripatopsis_capensis | Trinity | 12.9 | 11,638,180 | 100 | C |
| Duplicated | SRR1505119_Monodonta_labio | Trinity | 25.3 | 10,388,770 | 100 | C |
| Duplicated | SRR1560310_Lyonsia_floridana | Trinity | 24.3 | 9,919,645 | 100 | C |
| Duplicated | SRR1560359_Mercenaria_campechiensis | Trinity | 6.6 | 11,935,267 | 100 | C |
| Duplicated | SRR1560432_Neotrigonia_margaritacea | Trinity | 10.4 | 11,215,767 | 100 | C |
| Duplicated | SRR1560458_Cardites_antiquatus | Trinity | 10.4 | 11,916,756 | 100 | C |
| Duplicated | SRR1561723_Sphaerium_nucleus | Trinity | 18.2 | 18,539,173 | 100 | C |
| Duplicated | SRR1611556_Hemithiris_psittacea | Trinity | 65.4 | 9,221,875 | 100 | C |
| Duplicated | SRR1611560_Malacobdella_grossa | Trinity | 73.9 | 8,307,739 | 100 | C |
| Duplicated | SRR1611565_Phoronis_psammophila | Trinity | 71 | 12,949,999 | 100 | C |
| Duplicated | SRR1796434_Catenula_lemnae | Trinity | 20.8 | 3,028,636 | 100 | C |
| Duplicated | SRR3097584_Florometra | Trinity | 56.7 | 32,710,859 | 100 | C |
| Duplicated | SRR3105702_Childia_submaculatum | Trinity | 55.9 | 6,089,955 | 100 | C |
| Duplicated | SRR331123_Ennucula_tenuis | Trinity | 40.4 | 14,420,942 | 100 | C |
| Duplicated | SRR7655554_Ircinia_fasciculata | Trinity | 35.5 | 13,420,109 | 100 | C |
| Duplicated | SRR7754744_Krohnitta_subtilis | Trinity | 56 | 15,954,007 | 100 | C |
| Duplicated | SRR10527303_Octolasmis_warwickii | Trinity | 69.9 | 15,813,391 | 150 | D |
| Duplicated | SRR3350463_Dimya_lima | Trinity | 19.1 | 5,426,850 | 150 | D |
| Duplicated | SRR3404576_Brachionus_plicatilis | Trinity | 37.8 | 7,403,847 | 150 | D |
| Duplicated | SRR4113507_Calanus_finmarchicus | Trinity | 48.1 | 10,633,606 | 150 | D |
| Duplicated | SRR4294206_Millepora_alcicornis | Trinity | 48.6 | 24,645,545 | 150 | D |
| Duplicated | SRR504694_Corticium_candelabrum | Trinity | 57.5 | 18,897,095 | 150 | D |
| Duplicated | SRR5140141_Eoleptestheria_cf_ticinensis | Trinity | 40.3 | 5,471,351 | 150 | D |
| Duplicated | SRR5626553_Nautilus_pompilius | Trinity | 27.8 | 16,461,197 | 150 | D |
| Duplicated | SRR5873556_Neocalanus_Flemingeri | Trinity | 28.6 | 4,112,626 | 150 | D |
| Duplicated | SRR8393254_Apostichopus_japonicus | Trinity | 65.4 | 8,289,770 | 150 | D |
| Duplicated | SRR8491966_Porites_pukoensis | Trinity | 75.9 | 16,448,725 | 150 | D |
| Fragmented | ERR3026433_Pinnigorgia_flava | Trinity | 4.6 | 30,545,400 | 50 | A |
| Fragmented | ERR3026434_Sinularia_cruciata | Trinity | 11.3 | 22,160,908 | 50 | A |
| Fragmented | ERR3026435_Tubipora_musica | Trinity | 5.8 | 23,006,724 | 50 | A |
| Fragmented | ERR3040053_Heliopora_coerulea | Trinity | 8.5 | 29,000,821 | 50 | A |
| Fragmented | SRR8280777_Godzilliognomus_frondosus | Trinity | 11.3 | 14,086,834 | 75 | B |
| Fragmented | SRR8745910_Galeodes_sp | Trinity | 13.9 | 6,356,774 | 75 | B |
| Fragmented | SRR8745911_Peripatoides_novaezealandiae_onyc | Trinity | 9.3 | 5,768,550 | 75 | B |
| Fragmented | ERR1000783_Paracentrotus_lividus | Trinity | 18 | 6,803,316 | 75 | B |
| Fragmented | SRR10744002_Paracentrotus_lividus | Trinity | 4.8 | 13,583,857 | 75 | B |
| Fragmented | SRR1232685_Nephtys_caeca | Trinity | 28.2 | 1,576,665 | 75 | B |
| Fragmented | SRR1271607_Gnathostomula_paradoxa | Trinity | 25.6 | 5,954,962 | 75 | B |
| Fragmented | SRR1271706_Macrodasys_sp | Trinity | 15.4 | 3,204,609 | 75 | B |
| Fragmented | SRR1273732_Lepidodermella_squamata | Trinity | 25.8 | 4,370,938 | 75 | B |
| Fragmented | SRR1273789_Cephalothrix_linearis | Trinity | 16.2 | 4,869,244 | 75 | B |
| Fragmented | SRR5569439_Acropora_palmata | Trinity | 22.5 | 10,476,071 | 75 | B |

| | | | | | | |
|---|---|---|---|---|---|---|
| Fragmented | SRR8601367_Acropora_pulchra | Trinity | 8.3 | 14,037,157 | 75 | B |
| Fragmented | SRR1041944_Ephydatia_muelleri | Trinity | 1.5 | 11,425,188 | 100 | C |
| Fragmented | SRR1042012_Oscarella_pearsei | Trinity | 2.7 | 11,306,242 | 100 | C |
| Fragmented | SRR1168575_Spongilla_lacustris | Trinity | 10.2 | 5,136,881 | 100 | C |
| Fragmented | SRR1711043_Mycale_phylophylla_adult | Trinity | 13.4 | 11,408,543 | 100 | C |
| Fragmented | SRR1793376_Haliclona_tubifera | Trinity | 1.3 | 16,356,602 | 100 | C |
| Fragmented | SRR504690_Sycon_coactum | Trinity | 8.4 | 9,098,097 | 100 | C |
| Fragmented | SRR1145776_Peripatopsis_capensis | Trinity | 30.3 | 11,638,180 | 100 | C |
| Fragmented | SRR1505119_Monodonta_labio | Trinity | 12.3 | 10,388,770 | 100 | C |
| Fragmented | SRR1560310_Lyonsia_floridana | Trinity | 20.1 | 9,919,645 | 100 | C |
| Fragmented | SRR1560359_Mercenaria_campechiensis | Trinity | 21.8 | 11,935,267 | 100 | C |
| Fragmented | SRR1560432_Neotrigonia_margaritacea | Trinity | 27.8 | 11,215,767 | 100 | C |
| Fragmented | SRR1560458_Cardites_antiquatus | Trinity | 33.1 | 11,916,756 | 100 | C |
| Fragmented | SRR1561723_Sphaerium_nucleus | Trinity | 43.3 | 18,539,173 | 100 | C |
| Fragmented | SRR1611556_Hemithiris_psittacea | Trinity | 4.2 | 9,221,875 | 100 | C |
| Fragmented | SRR1611560_Malacobdella_grossa | Trinity | 2.6 | 8,307,739 | 100 | C |
| Fragmented | SRR1611565_Phoronis_psammophila | Trinity | 6.5 | 12,949,999 | 100 | C |
| Fragmented | SRR1796434_Catenula_lemnae | Trinity | 18.6 | 3,028,636 | 100 | C |
| Fragmented | SRR3097584_Florometra | Trinity | 1.2 | 32,710,859 | 100 | C |
| Fragmented | SRR3105702_Childia_submaculatum | Trinity | 5 | 6,089,955 | 100 | C |
| Fragmented | SRR331123_Ennucula_tenuis | Trinity | 4.7 | 14,420,942 | 100 | C |
| Fragmented | SRR7655554_Ircinia_fasciculata | Trinity | 19.4 | 13,420,109 | 100 | C |
| Fragmented | SRR7754744_Krohnitta_subtilis | Trinity | 8.3 | 15,954,007 | 100 | C |
| Fragmented | SRR10527303_Octolasmis_warwickii | Trinity | 0.7 | 15,813,391 | 150 | D |
| Fragmented | SRR3350463_Dimya_lima | Trinity | 29.9 | 5,426,850 | 150 | D |
| Fragmented | SRR3404576_Brachionus_plicatilis | Trinity | 0.4 | 7,403,847 | 150 | D |
| Fragmented | SRR4113507_Calanus_finmarchicus | Trinity | 16.8 | 10,633,606 | 150 | D |
| Fragmented | SRR4294206_Millepora_alcicornis | Trinity | 1 | 24,645,545 | 150 | D |
| Fragmented | SRR504694_Corticium_candelabrum | Trinity | 5.8 | 18,897,095 | 150 | D |
| Fragmented | SRR5140141_Eoleptestheria_cf_ticinensis | Trinity | 12.9 | 5,471,351 | 150 | D |
| Fragmented | SRR5626553_Nautilus_pompilius | Trinity | 32.8 | 16,461,197 | 150 | D |
| Fragmented | SRR5873556_Neocalanus_Flemingeri | Trinity | 7.3 | 4,112,626 | 150 | D |
| Fragmented | SRR8393254_Apostichopus_japonicus | Trinity | 7.3 | 8,289,770 | 150 | D |
| Fragmented | SRR8491966_Porites_pukoensis | Trinity | 0.6 | 16,448,725 | 150 | D |
| Missing | ERR3026433_Pinnigorgia_flava | Trinity | 1.8 | 30,545,400 | 50 | A |
| Missing | ERR3026434_Sinularia_cruciata | Trinity | 3.8 | 22,160,908 | 50 | A |
| Missing | ERR3026435_Tubipora_musica | Trinity | 2.6 | 23,006,724 | 50 | A |
| Missing | ERR3040053_Heliopora_coerulea | Trinity | 2 | 29,000,821 | 50 | A |
| Missing | SRR8280777_Godzilliognomus_frondosus | Trinity | 6.5 | 14,086,834 | 75 | B |
| Missing | SRR8745910_Galeodes_sp | Trinity | 3.3 | 6,356,774 | 75 | B |
| Missing | SRR8745911_Peripatoides_novaezealandiae_onyc | Trinity | 7.4 | 5,768,550 | 75 | B |
| Missing | ERR1000783_Paracentrotus_lividus | Trinity | 8.1 | 6,803,316 | 75 | B |
| Missing | SRR10744002_Paracentrotus_lividus | Trinity | 1.8 | 13,583,857 | 75 | B |
| Missing | SRR1232685_Nephtys_caeca | Trinity | 45.2 | 1,576,665 | 75 | B |
| Missing | SRR1271607_Gnathostomula_paradoxa | Trinity | 20.3 | 5,954,962 | 75 | B |
| Missing | SRR1271706_Macrodasys_sp | Trinity | 12 | 3,204,609 | 75 | B |
| Missing | SRR1273732_Lepidodermella_squamata | Trinity | 22.1 | 4,370,938 | 75 | B |
| Missing | SRR1273789_Cephalothrix_linearis | Trinity | 4.1 | 4,869,244 | 75 | B |
| Missing | SRR5569439_Acropora_palmata | Trinity | 11.4 | 10,476,071 | 75 | B |
| Missing | SRR8601367_Acropora_pulchra | Trinity | 1.9 | 14,037,157 | 75 | B |
| Missing | SRR1041944_Ephydatia_muelleri | Trinity | 4.1 | 11,425,188 | 100 | C |

| | | | | | | |
|---|---|---|---|---|---|---|
| Missing | SRR1042012_Oscarella_pearsei | Trinity | 4.7 | 11,306,242 | 100 | C |
| Missing | SRR1168575_Spongilla_lacustris | Trinity | 4.8 | 5,136,881 | 100 | C |
| Missing | SRR1711043_Mycale_phylophylla_adult | Trinity | 9.2 | 11,408,543 | 100 | C |
| Missing | SRR1793376_Haliclona_tubifera | Trinity | 4.5 | 16,356,602 | 100 | C |
| Missing | SRR504690_Sycon_coactum | Trinity | 9 | 9,098,097 | 100 | C |
| Missing | SRR1145776_Peripatopsis_capensis | Trinity | 8.1 | 11,638,180 | 100 | C |
| Missing | SRR1505119_Monodonta_labio | Trinity | 3.5 | 10,388,770 | 100 | C |
| Missing | SRR1560310_Lyonsia_floridana | Trinity | 3.2 | 9,919,645 | 100 | C |
| Missing | SRR1560359_Mercenaria_campechiensis | Trinity | 57.9 | 11,935,267 | 100 | C |
| Missing | SRR1560432_Neotrigonia_margaritacea | Trinity | 14.5 | 11,215,767 | 100 | C |
| Missing | SRR1560458_Cardites_antiquatus | Trinity | 24.2 | 11,916,756 | 100 | C |
| Missing | SRR1561723_Sphaerium_nucleus | Trinity | 22.1 | 18,539,173 | 100 | C |
| Missing | SRR1611556_Hemithiris_psittacea | Trinity | 1.3 | 9,221,875 | 100 | C |
| Missing | SRR1611560_Malacobdella_grossa | Trinity | 3.1 | 8,307,739 | 100 | C |
| Missing | SRR1611565_Phoronis_psammophila | Trinity | 0.9 | 12,949,999 | 100 | C |
| Missing | SRR1796434_Catenula_lemnae | Trinity | 28.9 | 3,028,636 | 100 | C |
| Missing | SRR3097584_Florometra | Trinity | 0.8 | 32,710,859 | 100 | C |
| Missing | SRR3105702_Childia_submaculatum | Trinity | 13.2 | 6,089,955 | 100 | C |
| Missing | SRR331123_Ennucula_tenuis | Trinity | 1.1 | 14,420,942 | 100 | C |
| Missing | SRR7655554_Ircinia_fasciculata | Trinity | 9.2 | 13,420,109 | 100 | C |
| Missing | SRR7754744_Krohnitta_subtilis | Trinity | 3.5 | 15,954,007 | 100 | C |
| Missing | SRR10527303_Octolasmis_warwickii | Trinity | 1.8 | 15,813,391 | 150 | D |
| Missing | SRR3350463_Dimya_lima | Trinity | 17 | 5,426,850 | 150 | D |
| Missing | SRR3404576_Brachionus_plicatilis | Trinity | 4.1 | 7,403,847 | 150 | D |
| Missing | SRR4113507_Calanus_finmarchicus | Trinity | 3.4 | 10,633,606 | 150 | D |
| Missing | SRR4294206_Millepora_alcicornis | Trinity | 1.7 | 24,645,545 | 150 | D |
| Missing | SRR504694_Corticium_candelabrum | Trinity | 3 | 18,897,095 | 150 | D |
| Missing | SRR5140141_Eoleptestheria_cf_ticinensis | Trinity | 2.3 | 5,471,351 | 150 | D |
| Missing | SRR5626553_Nautilus_pompilius | Trinity | 5 | 16,461,197 | 150 | D |
| Missing | SRR5873556_Neocalanus_Flemingeri | Trinity | 3.9 | 4,112,626 | 150 | D |
| Missing | SRR8393254_Apostichopus_japonicus | Trinity | 1.1 | 8,289,770 | 150 | D |
| Missing | SRR8491966_Porites_pukoensis | Trinity | 0 | 16,448,725 | 150 | D |
