## Supplementary material for "TransPi – a comprehensive TRanscriptome ANalysiS PIpeline for *de novo* transcriptome assembly": Supplementary_File_1.html

Suppplementary File 1 - TransPi model organisms BUSCO 50bp


### Suppplementary File 1 - TransPi model organisms BUSCO 50bp

```
# BUSCO plots all kmer sets
# setwd("~/Desktop/R/ramon/TransPi/paper/")
library(reshape2)
library(plotly)
library(dplyr)
```

### busco3\_50

```
csv=read.csv("busco3_50.csv", header=TRUE)
```

##### All BUSCO (all sets)

**Complete**

```
## Complete genes comparison
```

```
## [1] "One (or more) set is not normally distributed"
## [1] "Data not significant. Skipping pairwise comparison"
```

**Single**

```
## Single genes comparison
```

```
## [1] "One (or more) set is not normally distributed"
## [1] "Kruskal-Wallis test was significant (p<.05)"
```

```
##
##  Pairwise comparisons using Wilcoxon rank sum test
##
## data:  sing$Score and sing$Program
##
##         Transpi
## Trinity 3.9e-05
##
## P value adjustment method: BH
```

**Duplicated**

```
## Duplicated genes comparison
```

```
## [1] "One (or more) set is not normally distributed"
## [1] "Kruskal-Wallis test was significant (p<.05)"
```

```
##
##  Pairwise comparisons using Wilcoxon rank sum test
##
## data:  dup$Score and dup$Program
##
##         Transpi
## Trinity 8e-07
##
## P value adjustment method: BH
```

**Fragmented**

```
## Fragmented genes comparison
```

```
## [1] "One (or more) set is not normally distributed"
## [1] "Kruskal-Wallis test was significant (p<.05)"
```

```
##
##  Pairwise comparisons using Wilcoxon rank sum test
##
## data:  frag$Score and frag$Program
##
##         Transpi
## Trinity 7.9e-06
##
## P value adjustment method: BH
```

**Missing**

```
## Missing genes comparison
```

```
## [1] "One (or more) set is not normally distributed"
## [1] "Data not significant. Skipping pairwise comparison"
```

##### All BUSCO (kmer test)

**Only TransPi**

**Complete**

```
## Complete genes comparison
```

```
## [1] "One (or more) set is not normally distributed"
## [1] "Data not significant. Skipping pairwise comparison"
```

**Single**

```
## Single genes comparison
```

```
## [1] "One (or more) set is not normally distributed"
## [1] "Data not significant. Skipping pairwise comparison"
```

**Duplicated**

```
## Duplicated genes comparison
```

```
## [1] "One (or more) set is not normally distributed"
## [1] "Data not significant. Skipping pairwise comparison"
```

**Fragmented**

```
## Fragmented genes comparison
```

```
## [1] "One (or more) set is not normally distributed"
## [1] "Data not significant. Skipping pairwise comparison"
```

**Missing**

```
## Missing genes comparison
```

```
## [1] "One (or more) set is not normally distributed"
## [1] "Data not significant. Skipping pairwise comparison"
```

##### By species (all sets)

##### By species (kmer test)

#### CE

**Complete**

```
## Complete genes comparison
```

```
## [1] "All sets are normally distributed"
## [1] "ANOVA"
##             Df Sum Sq Mean Sq F value Pr(>F)
## Kmer         2  0.022 0.01083   0.097  0.908
## Residuals   33  3.686 0.11169
## [1] "Pairwise comparison"
```

```
##   Tukey multiple comparisons of means
##     95% family-wise confidence level
##
## Fit: aov(formula = Score ~ Kmer, data = ceTra2)
##
## $Kmer
##                    diff        lwr       upr     p adj
## KmerB-KmerA -0.05833333 -0.3931241 0.2764574 0.9044460
## KmerC-KmerA -0.04166667 -0.3764574 0.2931241 0.9499673
## KmerC-KmerB  0.01666667 -0.3181241 0.3514574 0.9918091
```

**Single**

```
## Single genes comparison
```

```
## [1] "All sets are normally distributed"
## [1] "ANOVA"
##             Df Sum Sq Mean Sq F value Pr(>F)
## Kmer         2  0.052  0.0258   0.041   0.96
## Residuals   33 21.028  0.6372
## [1] "Pairwise comparison"
```

```
##   Tukey multiple comparisons of means
##     95% family-wise confidence level
##
## Fit: aov(formula = Score ~ Kmer, data = ceTra2)
##
## $Kmer
##                    diff        lwr       upr     p adj
## KmerB-KmerA -0.03333333 -0.8329982 0.7663315 0.9942497
## KmerC-KmerA  0.05833333 -0.7413315 0.8579982 0.9825010
## KmerC-KmerB  0.09166667 -0.7079982 0.8913315 0.9573793
```

**Duplicated**

```
## Duplicated genes comparison
```

```
## [1] "All sets are normally distributed"
## [1] "ANOVA"
##             Df Sum Sq Mean Sq F value Pr(>F)
## Kmer         2  0.065  0.0325   0.081  0.923
## Residuals   33 13.282  0.4025
## [1] "Pairwise comparison"
```

```
##   Tukey multiple comparisons of means
##     95% family-wise confidence level
##
## Fit: aov(formula = Score ~ Kmer, data = ceTra2)
##
## $Kmer
##               diff        lwr       upr     p adj
## KmerB-KmerA -0.025 -0.6605438 0.6105438 0.9948775
## KmerC-KmerA -0.100 -0.7355438 0.5355438 0.9213138
## KmerC-KmerB -0.075 -0.7105438 0.5605438 0.9548928
```

**Fragmented**

```
## Fragmented genes comparison
```

```
## [1] "One (or more) set is not normally distributed"
## [1] "Data not significant. Skipping pairwise comparison"
```

**Missing**

```
## Fragmented genes comparison
```

```
## [1] "All sets are normally distributed"
## [1] "ANOVA"
##             Df Sum Sq Mean Sq F value Pr(>F)
## Kmer         2 0.0117 0.00583   0.083  0.921
## Residuals   33 2.3183 0.07025
## [1] "Pairwise comparison"
```

```
##   Tukey multiple comparisons of means
##     95% family-wise confidence level
##
## Fit: aov(formula = Score ~ Kmer, data = ceTra2)
##
## $Kmer
##                     diff        lwr       upr     p adj
## KmerB-KmerA  0.041666667 -0.2238511 0.3071844 0.9217148
## KmerC-KmerA  0.008333333 -0.2571844 0.2738511 0.9967358
## KmerC-KmerB -0.033333333 -0.2988511 0.2321844 0.9491153
```

**BUSCO and reads**

```
##   comp.Program comp.Category comp.Score comp.Reads comp.Sample
## 1      Transpi      Complete       85.5 24,017,399         CE1
## 2      Transpi      Complete       85.3 24,409,279         CE2
## 3      Transpi      Complete       85.3 24,694,386         CE3
## 4      Transpi      Complete       85.7 26,159,433         CE4
## 5      Transpi      Complete       85.4 26,417,070         CE5
## 6      Transpi      Complete       85.2 24,711,402         CE6
```

#### DM

**Complete**

```
## Complete genes comparison
```

```
## [1] "One (or more) set is not normally distributed"
## [1] "Data not significant. Skipping pairwise comparison"
```

**Single**

```
## Single genes comparison
```

```
## [1] "All sets are normally distributed"
## [1] "ANOVA"
##             Df Sum Sq Mean Sq F value Pr(>F)
## Kmer         2      0    0.24   0.003  0.997
## Residuals   51   4117   80.72
## [1] "Pairwise comparison"
```

```
##   Tukey multiple comparisons of means
##     95% family-wise confidence level
##
## Fit: aov(formula = Score ~ Kmer, data = dmTra2)
##
## $Kmer
##                    diff       lwr      upr     p adj
## KmerB-KmerA -0.22222222 -7.451660 7.007215 0.9969692
## KmerC-KmerA -0.05555556 -7.284993 7.173882 0.9998103
## KmerC-KmerB  0.16666667 -7.062771 7.396104 0.9982940
```

**Duplicated**

```
## Duplicated genes comparison
```

```
## [1] "One (or more) set is not normally distributed"
## [1] "Data not significant. Skipping pairwise comparison"
```

**Fragmented**

```
## Fragmented genes comparison
```

```
## [1] "One (or more) set is not normally distributed"
## [1] "Data not significant. Skipping pairwise comparison"
```

**Missing**

```
## Missing genes comparison
```

```
## [1] "One (or more) set is not normally distributed"
## [1] "Data not significant. Skipping pairwise comparison"
```

**BUSCO and reads**

```
##   comp.Program comp.Category comp.Score comp.Reads comp.Sample
## 1      Transpi      Complete       44.7 15,648,174         DM1
## 2      Transpi      Complete       44.6 15,181,806         DM2
## 3      Transpi      Complete       37.0  9,857,308         DM3
## 4      Transpi      Complete       39.8 15,940,368         DM4
## 5      Transpi      Complete       48.5 18,874,930         DM5
## 6      Transpi      Complete       35.1 21,169,448         DM6
```

#### MM

**Complete**

```
## Complete genes comparison
```

```
## [1] "All sets are normally distributed"
## [1] "ANOVA"
##             Df Sum Sq Mean Sq F value Pr(>F)
## Kmer         2   0.02  0.0102   0.005  0.995
## Residuals   39  80.18  2.0558
## [1] "Pairwise comparison"
```

```
##   Tukey multiple comparisons of means
##     95% family-wise confidence level
##
## Fit: aov(formula = Score ~ Kmer, data = mmTra2)
##
## $Kmer
##                    diff       lwr      upr     p adj
## KmerB-KmerA 0.042857143 -1.277442 1.363156 0.9965582
## KmerC-KmerA 0.050000000 -1.270299 1.370299 0.9953184
## KmerC-KmerB 0.007142857 -1.313156 1.327442 0.9999042
```

**Single**

```
## Single genes comparison
```

```
## [1] "One (or more) set is not normally distributed"
## [1] "Data not significant. Skipping pairwise comparison"
```

**Duplicated**

```
## Duplicated genes comparison
```

```
## [1] "One (or more) set is not normally distributed"
## [1] "Data not significant. Skipping pairwise comparison"
```

**Fragmented**

```
## Fragmented genes comparison
```

```
## [1] "One (or more) set is not normally distributed"
## [1] "Data not significant. Skipping pairwise comparison"
```

**Missing**

```
## Missing genes comparison
```

```
## [1] "All sets are normally distributed"
## [1] "ANOVA"
##             Df Sum Sq Mean Sq F value Pr(>F)
## Kmer         2  0.078  0.0388   0.089  0.915
## Residuals   39 17.079  0.4379
## [1] "Pairwise comparison"
```

```
##   Tukey multiple comparisons of means
##     95% family-wise confidence level
##
## Fit: aov(formula = Score ~ Kmer, data = mmTra2)
##
## $Kmer
##                    diff        lwr       upr     p adj
## KmerB-KmerA -0.10000000 -0.7093637 0.5093637 0.9158524
## KmerC-KmerA -0.07857143 -0.6879351 0.5307923 0.9471300
## KmerC-KmerB  0.02142857 -0.5879351 0.6307923 0.9959619
```

**BUSCO and reads**

```
##   comp.Program comp.Category comp.Score comp.Reads comp.Sample
## 1      Transpi      Complete       96.9 33,540,466         MM1
## 2      Transpi      Complete       96.4 43,541,512         MM2
## 3      Transpi      Complete       97.0 43,440,008         MM3
## 4      Transpi      Complete       95.8 33,595,655         MM4
## 5      Transpi      Complete       95.0 33,174,108         MM5
## 6      Transpi      Complete       95.5 35,030,665         MM6
```

##### By sample (all sets)
