## Supplementary material for "TransPi – a comprehensive TRanscriptome ANalysiS PIpeline for *de novo* transcriptome assembly": Supplementary_File_2.html

Suppplementary File 2 - TransPi model organisms BUSCO 75bp


### Suppplementary File 2 - TransPi model organisms BUSCO 75bp

```
# BUSCO plots all kmer sets
# setwd("~/Desktop/R/ramon/TransPi/paper/")
library(reshape2)
library(plotly)
library(dplyr)
```

### busco3\_75

```
csv=read.csv("busco3_75.csv", header=TRUE)
```

##### All BUSCO (all sets)

**Complete**

```
## Complete genes comparison
```

```
## [1] "One (or more) set is not normally distributed"
## [1] "Kruskal-Wallis test was significant (p<.05)"
```

```
##
##  Pairwise comparisons using Wilcoxon rank sum test
##
## data:  comp$Score and comp$Program
##
##         Transpi
## Trinity 0.013
##
## P value adjustment method: BH
```

**Single**

```
## Single genes comparison
```

```
## [1] "One (or more) set is not normally distributed"
## [1] "Kruskal-Wallis test was significant (p<.05)"
```

```
##
##  Pairwise comparisons using Wilcoxon rank sum test
##
## data:  sing$Score and sing$Program
##
##         Transpi
## Trinity 3.2e-16
##
## P value adjustment method: BH
```

**Duplicated**

```
## Duplicated genes comparison
```

```
## [1] "One (or more) set is not normally distributed"
## [1] "Kruskal-Wallis test was significant (p<.05)"
```

```
##
##  Pairwise comparisons using Wilcoxon rank sum test
##
## data:  dup$Score and dup$Program
##
##         Transpi
## Trinity 4.3e-08
##
## P value adjustment method: BH
```

**Fragmented**

```
## Fragmented genes comparison
```

```
## [1] "One (or more) set is not normally distributed"
## [1] "Data not significant. Skipping pairwise comparison"
```

**Missing**

```
## Missing genes comparison
```

```
## [1] "One (or more) set is not normally distributed"
## [1] "Kruskal-Wallis test was significant (p<.05)"
```

```
##
##  Pairwise comparisons using Wilcoxon rank sum test
##
## data:  mis$Score and mis$Program
##
##         Transpi
## Trinity 0.0041
##
## P value adjustment method: BH
```

##### All BUSCO (kmer test)

**Only TransPi**

**Complete**

```
## Complete genes comparison
```

```
## [1] "One (or more) set is not normally distributed"
## [1] "Data not significant. Skipping pairwise comparison"
```

**Single**

```
## Single genes comparison
```

```
## [1] "All sets are normally distributed"
## [1] "ANOVA"
##              Df Sum Sq Mean Sq F value Pr(>F)
## singTra$Kmer  2    2.0   0.985   0.083  0.921
## Residuals    48  572.1  11.920
## [1] "Pairwise comparison"
```

```
##   Tukey multiple comparisons of means
##     95% family-wise confidence level
##
## Fit: aov(formula = singTra$Score ~ singTra$Kmer, data = singTra2)
##
## $`singTra$Kmer`
##                  diff       lwr      upr     p adj
## KmerB-KmerA 0.3411765 -2.522761 3.205114 0.9553164
## KmerC-KmerA 0.4647059 -2.399232 3.328644 0.9187644
## KmerC-KmerB 0.1235294 -2.740409 2.987467 0.9940194
```

**Duplicated**

```
## Duplicated genes comparison
```

```
## [1] "All sets are normally distributed"
## [1] "ANOVA"
##             Df Sum Sq Mean Sq F value Pr(>F)
## dupTra$Kmer  2    2.7   1.356   0.047  0.954
## Residuals   48 1377.0  28.687
## [1] "Pairwise comparison"
```

```
##   Tukey multiple comparisons of means
##     95% family-wise confidence level
##
## Fit: aov(formula = dupTra$Score ~ dupTra$Kmer, data = dupTra2)
##
## $`dupTra$Kmer`
##                   diff       lwr      upr     p adj
## KmerB-KmerA -0.2941176 -4.737159 4.148924 0.9859730
## KmerC-KmerA -0.5647059 -5.007748 3.878336 0.9493041
## KmerC-KmerB -0.2705882 -4.713630 4.172453 0.9881141
```

**Fragmented**

```
## Fragmented genes comparison
```

```
## [1] "All sets are normally distributed"
## [1] "ANOVA"
##              Df Sum Sq Mean Sq F value Pr(>F)
## fragTra$Kmer  2  0.184   0.092   0.191  0.827
## Residuals    48 23.136   0.482
## [1] "Pairwise comparison"
```

```
##   Tukey multiple comparisons of means
##     95% family-wise confidence level
##
## Fit: aov(formula = fragTra$Score ~ fragTra$Kmer, data = fragTra2)
##
## $`fragTra$Kmer`
##                    diff        lwr       upr     p adj
## KmerB-KmerA -0.07647059 -0.6523908 0.4994497 0.9448074
## KmerC-KmerA  0.07058824 -0.5053320 0.6465085 0.9527663
## KmerC-KmerB  0.14705882 -0.4288614 0.7229791 0.8112562
```

**Missing**

```
## Missing genes comparison
```

```
## [1] "One (or more) set is not normally distributed"
## [1] "Data not significant. Skipping pairwise comparison"
```

##### By species (all sets)

##### By species (kmer test)

#### CE

**Complete**

```
## Complete genes comparison
```

```
## [1] "All sets are normally distributed"
## [1] "ANOVA"
##             Df Sum Sq Mean Sq F value Pr(>F)
## Kmer         2  0.021  0.0103   0.011  0.989
## Residuals   27 24.953  0.9242
## [1] "Pairwise comparison"
```

```
##   Tukey multiple comparisons of means
##     95% family-wise confidence level
##
## Fit: aov(formula = Score ~ Kmer, data = ceTra2)
##
## $Kmer
##              diff       lwr      upr     p adj
## KmerB-KmerA  0.06 -1.005968 1.125968 0.9893241
## KmerC-KmerA  0.01 -1.055968 1.075968 0.9997018
## KmerC-KmerB -0.05 -1.115968 1.015968 0.9925730
```

**Single**

```
## Single genes comparison
```

```
## [1] "All sets are normally distributed"
## [1] "ANOVA"
##             Df Sum Sq Mean Sq F value Pr(>F)
## Kmer         2    0.2   0.117   0.008  0.992
## Residuals   27  381.4  14.124
## [1] "Pairwise comparison"
```

```
##   Tukey multiple comparisons of means
##     95% family-wise confidence level
##
## Fit: aov(formula = Score ~ Kmer, data = ceTra2)
##
## $Kmer
##              diff      lwr     upr     p adj
## KmerB-KmerA  0.21 -3.95722 4.37722 0.9914329
## KmerC-KmerA  0.15 -4.01722 4.31722 0.9956190
## KmerC-KmerB -0.06 -4.22722 4.10722 0.9992976
```

**Duplicated**

```
## Duplicated genes comparison
```

```
## [1] "All sets are normally distributed"
## [1] "ANOVA"
##             Df Sum Sq Mean Sq F value Pr(>F)
## Kmer         2    0.1    0.07   0.003  0.997
## Residuals   27  584.7   21.66
## [1] "Pairwise comparison"
```

```
##   Tukey multiple comparisons of means
##     95% family-wise confidence level
##
## Fit: aov(formula = Score ~ Kmer, data = ceTra2)
##
## $Kmer
##              diff       lwr      upr     p adj
## KmerB-KmerA -0.15 -5.310022 5.010022 0.9971403
## KmerC-KmerA -0.14 -5.300022 5.020022 0.9975084
## KmerC-KmerB  0.01 -5.150022 5.170022 0.9999873
```

**Fragmented**

```
## Fragmented genes comparison
```

```
## [1] "One (or more) set is not normally distributed"
## [1] "Data not significant. Skipping pairwise comparison"
```

**Missing**

```
## Fragmented genes comparison
```

```
## [1] "All sets are normally distributed"
## [1] "ANOVA"
##             Df Sum Sq Mean Sq F value Pr(>F)
## Kmer         2  0.008  0.0040   0.005  0.995
## Residuals   27 20.932  0.7753
## [1] "Pairwise comparison"
```

```
##   Tukey multiple comparisons of means
##     95% family-wise confidence level
##
## Fit: aov(formula = Score ~ Kmer, data = ceTra2)
##
## $Kmer
##              diff       lwr      upr     p adj
## KmerB-KmerA -0.02 -0.996311 0.956311 0.9985788
## KmerC-KmerA -0.04 -1.016311 0.936311 0.9943282
## KmerC-KmerB -0.02 -0.996311 0.956311 0.9985788
```

**BUSCO and reads**

```
##   comp.Program comp.Category comp.Score comp.Reads comp.Sample
## 1      Transpi      Complete       86.0 40,302,838         CE1
## 2      Transpi      Complete       86.2 50,516,835         CE2
## 3      Transpi      Complete       83.9 41,947,175         CE3
## 4      Transpi      Complete       83.8 44,969,393         CE4
## 5      Transpi      Complete       84.1 45,605,396         CE5
## 6      Transpi      Complete       86.2 40,302,838         CE1
```

#### DM

**Complete**

```
## Complete genes comparison
```

```
## [1] "All sets are normally distributed"
## [1] "ANOVA"
##             Df Sum Sq Mean Sq F value Pr(>F)
## Kmer         2  0.134  0.0669   0.148  0.863
## Residuals   33 14.923  0.4522
## [1] "Pairwise comparison"
```

```
##   Tukey multiple comparisons of means
##     95% family-wise confidence level
##
## Fit: aov(formula = Score ~ Kmer, data = dmTra2)
##
## $Kmer
##                     diff        lwr       upr     p adj
## KmerB-KmerA  0.008333333 -0.6653044 0.6819710 0.9994921
## KmerC-KmerA -0.125000000 -0.7986377 0.5486377 0.8923866
## KmerC-KmerB -0.133333333 -0.8069710 0.5403044 0.8785582
```

**Single**

```
## Single genes comparison
```

```
## [1] "One (or more) set is not normally distributed"
## [1] "Data not significant. Skipping pairwise comparison"
```

**Duplicated**

```
## Duplicated genes comparison
```

```
## [1] "One (or more) set is not normally distributed"
## [1] "Data not significant. Skipping pairwise comparison"
```

**Fragmented**

```
## Fragmented genes comparison
```

```
## [1] "One (or more) set is not normally distributed"
## [1] "Data not significant. Skipping pairwise comparison"
```

**Missing**

```
## Missing genes comparison
```

```
## [1] "All sets are normally distributed"
## [1] "ANOVA"
##             Df Sum Sq Mean Sq F value Pr(>F)
## Kmer         2  0.032 0.01583   0.055  0.947
## Residuals   33  9.498 0.28783
## [1] "Pairwise comparison"
```

```
##   Tukey multiple comparisons of means
##     95% family-wise confidence level
##
## Fit: aov(formula = Score ~ Kmer, data = dmTra2)
##
## $Kmer
##                     diff        lwr       upr     p adj
## KmerB-KmerA  0.066666667 -0.4707725 0.6041058 0.9502882
## KmerC-KmerA  0.058333333 -0.4791058 0.5957725 0.9616976
## KmerC-KmerB -0.008333333 -0.5457725 0.5291058 0.9992022
```

**BUSCO and reads**

```
##   comp.Program comp.Category comp.Score  comp.Reads comp.Sample
## 1      Transpi      Complete       97.1  87,423,452         DM1
## 2      Transpi      Complete       97.6  85,714,154         DM2
## 3      Transpi      Complete       96.6  88,252,694         DM3
## 4      Transpi      Complete       97.4  82,110,608         DM4
## 5      Transpi      Complete       96.8 102,413,880         DM5
## 6      Transpi      Complete       97.6  98,529,578         DM6
```

#### MM

**Complete**

```
## Complete genes comparison
```

```
## [1] "One (or more) set is not normally distributed"
## [1] "Data not significant. Skipping pairwise comparison"
```

**Single**

```
## Single genes comparison
```

```
## [1] "One (or more) set is not normally distributed"
## [1] "Data not significant. Skipping pairwise comparison"
```

**Duplicated**

```
## Duplicated genes comparison
```

```
## [1] "One (or more) set is not normally distributed"
## [1] "Data not significant. Skipping pairwise comparison"
```

**Fragmented**

```
## Fragmented genes comparison
```

```
## [1] "All sets are normally distributed"
## [1] "ANOVA"
##             Df Sum Sq Mean Sq F value Pr(>F)
## Kmer         2   0.01  0.0036   0.003  0.997
## Residuals   33  47.05  1.4257
## [1] "Pairwise comparison"
```

```
##   Tukey multiple comparisons of means
##     95% family-wise confidence level
##
## Fit: aov(formula = Score ~ Kmer, data = mmTra2)
##
## $Kmer
##                    diff       lwr      upr     p adj
## KmerB-KmerA 0.025000000 -1.171128 1.221128 0.9985510
## KmerC-KmerA 0.033333333 -1.162795 1.229462 0.9974255
## KmerC-KmerB 0.008333333 -1.187795 1.204462 0.9998389
```

**Missing**

```
## Missing genes comparison
```

```
## [1] "All sets are normally distributed"
## [1] "ANOVA"
##             Df Sum Sq Mean Sq F value Pr(>F)
## Kmer         2  0.004  0.0019   0.002  0.998
## Residuals   33 30.286  0.9178
## [1] "Pairwise comparison"
```

```
##   Tukey multiple comparisons of means
##     95% family-wise confidence level
##
## Fit: aov(formula = Score ~ Kmer, data = mmTra2)
##
## $Kmer
##                     diff        lwr       upr     p adj
## KmerB-KmerA  0.008333333 -0.9513443 0.9680109 0.9997497
## KmerC-KmerA -0.016666667 -0.9763443 0.9430109 0.9989993
## KmerC-KmerB -0.025000000 -0.9846776 0.9346776 0.9977499
```

**BUSCO and reads**

```
##   comp.Program comp.Category comp.Score comp.Reads comp.Sample
## 1      Transpi      Complete       98.0 33,700,156         MM1
## 2      Transpi      Complete       98.0 41,236,457         MM2
## 3      Transpi      Complete       98.0 35,598,598         MM3
## 4      Transpi      Complete       93.9 41,745,958         MM4
## 5      Transpi      Complete       96.1 45,329,544         MM5
## 6      Transpi      Complete       94.5 44,469,310         MM6
```

##### By sample (all sets)
