## Supplementary material for "TransPi – a comprehensive TRanscriptome ANalysiS PIpeline for *de novo* transcriptome assembly": Supplementary_File_3.html

Suppplementary File 3 - TransPi model organisms BUSCO 100bp


### Suppplementary File 3 - TransPi model organisms BUSCO 100bp

```
# BUSCO plots all kmer sets
# setwd("~/Desktop/R/ramon/TransPi/paper/")
library(reshape2)
library(plotly)
library(dplyr)
```

### busco3\_100

```
csv=read.csv("busco3_100.csv", header=TRUE)
```

##### All BUSCO (all sets)

**Complete**

```
## Complete genes comparison
```

```
## [1] "One (or more) set is not normally distributed"
## [1] "Data not significant. Skipping pairwise comparison"
```

**Single**

```
## Single genes comparison
```

```
## [1] "One (or more) set is not normally distributed"
## [1] "Kruskal-Wallis test was significant (p<.05)"
```

```
##
##  Pairwise comparisons using Wilcoxon rank sum test
##
## data:  sing$Score and sing$Program
##
##         Transpi
## Trinity 6.1e-14
##
## P value adjustment method: BH
```

**Duplicated**

```
## Duplicated genes comparison
```

```
## [1] "One (or more) set is not normally distributed"
## [1] "Kruskal-Wallis test was significant (p<.05)"
```

```
##
##  Pairwise comparisons using Wilcoxon rank sum test
##
## data:  dup$Score and dup$Program
##
##         Transpi
## Trinity 3.3e-08
##
## P value adjustment method: BH
```

**Fragmented**

```
## Fragmented genes comparison
```

```
## [1] "One (or more) set is not normally distributed"
## [1] "Data not significant. Skipping pairwise comparison"
```

**Missing**

```
## Missing genes comparison
```

```
## [1] "One (or more) set is not normally distributed"
## [1] "Data not significant. Skipping pairwise comparison"
```

##### All BUSCO (kmer test)

**Only TransPi**

**Complete**

```
## Complete genes comparison
```

```
## [1] "One (or more) set is not normally distributed"
## [1] "Data not significant. Skipping pairwise comparison"
```

**Single**

```
## Single genes comparison
```

```
## [1] "One (or more) set is not normally distributed"
## [1] "Data not significant. Skipping pairwise comparison"
```

**Duplicated**

```
## Duplicated genes comparison
```

```
## [1] "One (or more) set is not normally distributed"
## [1] "Data not significant. Skipping pairwise comparison"
```

**Fragmented**

```
## Fragmented genes comparison
```

```
## [1] "One (or more) set is not normally distributed"
## [1] "Data not significant. Skipping pairwise comparison"
```

**Missing**

```
## Missing genes comparison
```

```
## [1] "One (or more) set is not normally distributed"
## [1] "Data not significant. Skipping pairwise comparison"
```

##### By species (all sets)

##### By species (kmer test)

#### CE

**Complete**

```
## Complete genes comparison
```

```
## [1] "One (or more) set is not normally distributed"
## [1] "Data not significant. Skipping pairwise comparison"
```

**Single**

```
## Single genes comparison
```

```
## [1] "All sets are normally distributed"
## [1] "ANOVA"
##             Df Sum Sq Mean Sq F value Pr(>F)
## Kmer         2    0.8    0.38   0.004  0.996
## Residuals   33 2862.0   86.73
## [1] "Pairwise comparison"
```

```
##   Tukey multiple comparisons of means
##     95% family-wise confidence level
##
## Fit: aov(formula = Score ~ Kmer, data = ceTra2)
##
## $Kmer
##                    diff       lwr      upr     p adj
## KmerB-KmerA -0.33333333 -9.662397 8.995731 0.9957715
## KmerC-KmerA -0.27500000 -9.604064 9.054064 0.9971199
## KmerC-KmerB  0.05833333 -9.270731 9.387397 0.9998702
```

**Duplicated**

```
## Duplicated genes comparison
```

```
## [1] "All sets are normally distributed"
## [1] "ANOVA"
##             Df Sum Sq Mean Sq F value Pr(>F)
## Kmer         2    0.6    0.31   0.009  0.991
## Residuals   33 1138.3   34.49
## [1] "Pairwise comparison"
```

```
##   Tukey multiple comparisons of means
##     95% family-wise confidence level
##
## Fit: aov(formula = Score ~ Kmer, data = ceTra2)
##
## $Kmer
##                   diff       lwr      upr     p adj
## KmerB-KmerA  0.3166667 -5.566869 6.200203 0.9904328
## KmerC-KmerA  0.2000000 -5.683536 6.083536 0.9961719
## KmerC-KmerB -0.1166667 -6.000203 5.766869 0.9986956
```

**Fragmented**

```
## Fragmented genes comparison
```

```
## [1] "One (or more) set is not normally distributed"
## [1] "Data not significant. Skipping pairwise comparison"
```

**Missing**

```
## Fragmented genes comparison
```

```
## [1] "All sets are normally distributed"
## [1] "ANOVA"
##             Df Sum Sq Mean Sq F value Pr(>F)
## Kmer         2      0   0.002       0      1
## Residuals   33    246   7.455
## [1] "Pairwise comparison"
```

```
##   Tukey multiple comparisons of means
##     95% family-wise confidence level
##
## Fit: aov(formula = Score ~ Kmer, data = ceTra2)
##
## $Kmer
##                     diff       lwr      upr     p adj
## KmerB-KmerA  0.025000000 -2.710109 2.760109 0.9997227
## KmerC-KmerA  0.016666667 -2.718442 2.751776 0.9998767
## KmerC-KmerB -0.008333333 -2.743442 2.726776 0.9999692
```

**BUSCO and reads**

```
##   comp.Program comp.Category comp.Score comp.Reads comp.Sample
## 1      Transpi      Complete       82.3 49,743,412         CE1
## 2      Transpi      Complete       82.3 38,836,876         CE2
## 3      Transpi      Complete       67.1 11,166,310         CE3
## 4      Transpi      Complete       80.4 11,626,153         CE4
## 5      Transpi      Complete       85.3 12,567,755         CE5
## 6      Transpi      Complete       69.6 14,372,593         CE6
```

#### DM

**Complete**

```
## Complete genes comparison
```

```
## [1] "All sets are normally distributed"
## [1] "ANOVA"
##             Df Sum Sq Mean Sq F value Pr(>F)
## Kmer         2  0.112  0.0558   0.084   0.92
## Residuals   33 21.968  0.6657
## [1] "Pairwise comparison"
```

```
##   Tukey multiple comparisons of means
##     95% family-wise confidence level
##
## Fit: aov(formula = Score ~ Kmer, data = dmTra2)
##
## $Kmer
##                   diff        lwr       upr     p adj
## KmerB-KmerA 0.04166667 -0.7756760 0.8590093 0.9914127
## KmerC-KmerA 0.13333333 -0.6840093 0.9506760 0.9156942
## KmerC-KmerB 0.09166667 -0.7256760 0.9090093 0.9591627
```

**Single**

```
## Single genes comparison
```

```
## [1] "One (or more) set is not normally distributed"
## [1] "Data not significant. Skipping pairwise comparison"
```

**Duplicated**

```
## Duplicated genes comparison
```

```
## [1] "One (or more) set is not normally distributed"
## [1] "Data not significant. Skipping pairwise comparison"
```

**Fragmented**

```
## Fragmented genes comparison
```

```
## [1] "All sets are normally distributed"
## [1] "ANOVA"
##             Df Sum Sq Mean Sq F value Pr(>F)
## Kmer         2  0.037 0.01861   0.179  0.837
## Residuals   33  3.439 0.10422
## [1] "Pairwise comparison"
```

```
##   Tukey multiple comparisons of means
##     95% family-wise confidence level
##
## Fit: aov(formula = Score ~ Kmer, data = dmTra2)
##
## $Kmer
##                    diff        lwr       upr     p adj
## KmerB-KmerA  0.01666667 -0.3067275 0.3400609 0.9912244
## KmerC-KmerA -0.05833333 -0.3817275 0.2650609 0.8979782
## KmerC-KmerB -0.07500000 -0.3983942 0.2483942 0.8373956
```

**Missing**

```
## Missing genes comparison
```

```
## [1] "All sets are normally distributed"
## [1] "ANOVA"
##             Df Sum Sq Mean Sq F value Pr(>F)
## Kmer         2  0.037  0.0186    0.05  0.952
## Residuals   33 12.386  0.3753
## [1] "Pairwise comparison"
```

```
##   Tukey multiple comparisons of means
##     95% family-wise confidence level
##
## Fit: aov(formula = Score ~ Kmer, data = dmTra2)
##
## $Kmer
##                    diff        lwr       upr     p adj
## KmerB-KmerA -0.05833333 -0.6720504 0.5553838 0.9704847
## KmerC-KmerA -0.07500000 -0.6887171 0.5387171 0.9517128
## KmerC-KmerB -0.01666667 -0.6303838 0.5970504 0.9975550
```

**BUSCO and reads**

```
##   comp.Program comp.Category comp.Score comp.Reads comp.Sample
## 1      Transpi      Complete       95.4 30,515,068         DM1
## 2      Transpi      Complete       95.8 24,284,630         DM2
## 3      Transpi      Complete       95.0 24,773,404         DM3
## 4      Transpi      Complete       95.6 30,140,704         DM4
## 5      Transpi      Complete       94.6 25,828,680         DM5
## 6      Transpi      Complete       95.2 19,049,236         DM6
```

#### MM

**Complete**

```
## Complete genes comparison
```

```
## [1] "One (or more) set is not normally distributed"
## [1] "Data not significant. Skipping pairwise comparison"
```

**Single**

```
## Single genes comparison
```

```
## [1] "One (or more) set is not normally distributed"
## [1] "Data not significant. Skipping pairwise comparison"
```

**Duplicated**

```
## Duplicated genes comparison
```

```
## [1] "One (or more) set is not normally distributed"
## [1] "Data not significant. Skipping pairwise comparison"
```

**Fragmented**

```
## Fragmented genes comparison
```

```
## [1] "One (or more) set is not normally distributed"
## [1] "Data not significant. Skipping pairwise comparison"
```

**Missing**

```
## Missing genes comparison
```

```
## [1] "One (or more) set is not normally distributed"
## [1] "Data not significant. Skipping pairwise comparison"
```

**BUSCO and reads**

```
##   comp.Program comp.Category comp.Score  comp.Reads comp.Sample
## 1      Transpi      Complete       96.3 182,180,123         MM1
## 2      Transpi      Complete       97.0 107,280,600         MM2
## 3      Transpi      Complete       97.3 127,017,374         MM3
## 4      Transpi      Complete       97.3 102,909,165         MM4
## 5      Transpi      Complete       97.3  65,372,078         MM5
## 6      Transpi      Complete       53.9  27,016,597         MM6
```

##### By sample (all sets)
