## Supplementary material for "TransPi – a comprehensive TRanscriptome ANalysiS PIpeline for *de novo* transcriptome assembly": Supplementary_File_4.html

Suppplementary File 4 - TransPi model organisms BUSCO 150bp


### Suppplementary File 4 - TransPi model organisms BUSCO 150bp

```
# BUSCO plots all kmer sets
# setwd("~/Desktop/R/ramon/TransPi/paper/")
library(reshape2)
library(plotly)
library(dplyr)
```

### busco3\_150

```
csv=read.csv("busco3_150.csv", header=TRUE)
```

##### All BUSCO (all sets)

**Complete**

```
## Complete genes comparison
```

```
## [1] "One (or more) set is not normally distributed"
## [1] "Kruskal-Wallis test was significant (p<.05)"
```

```
##
##  Pairwise comparisons using Wilcoxon rank sum test
##
## data:  comp$Score and comp$Program
##
##         Transpi
## Trinity 0.02
##
## P value adjustment method: BH
```

**Single**

```
## Single genes comparison
```

```
## [1] "One (or more) set is not normally distributed"
## [1] "Kruskal-Wallis test was significant (p<.05)"
```

```
##
##  Pairwise comparisons using Wilcoxon rank sum test
##
## data:  sing$Score and sing$Program
##
##         Transpi
## Trinity <2e-16
##
## P value adjustment method: BH
```

**Duplicated**

```
## Duplicated genes comparison
```

```
## [1] "One (or more) set is not normally distributed"
## [1] "Kruskal-Wallis test was significant (p<.05)"
```

```
##
##  Pairwise comparisons using Wilcoxon rank sum test
##
## data:  dup$Score and dup$Program
##
##         Transpi
## Trinity 8.4e-09
##
## P value adjustment method: BH
```

**Fragmented**

```
## Fragmented genes comparison
```

```
## [1] "One (or more) set is not normally distributed"
## [1] "Kruskal-Wallis test was significant (p<.05)"
```

```
##
##  Pairwise comparisons using Wilcoxon rank sum test
##
## data:  frag$Score and frag$Program
##
##         Transpi
## Trinity 0.031
##
## P value adjustment method: BH
```

**Missing**

```
## Missing genes comparison
```

```
## [1] "One (or more) set is not normally distributed"
## [1] "Kruskal-Wallis test was significant (p<.05)"
```

```
##
##  Pairwise comparisons using Wilcoxon rank sum test
##
## data:  mis$Score and mis$Program
##
##         Transpi
## Trinity 0.021
##
## P value adjustment method: BH
```

##### All BUSCO (kmer test)

**Only TransPi**

**Complete**

```
## Complete genes comparison
```

```
## [1] "One (or more) set is not normally distributed"
## [1] "Data not significant. Skipping pairwise comparison"
```

**Single**

```
## Single genes comparison
```

```
## [1] "One (or more) set is not normally distributed"
## [1] "Data not significant. Skipping pairwise comparison"
```

**Duplicated**

```
## Duplicated genes comparison
```

```
## [1] "All sets are normally distributed"
## [1] "ANOVA"
##             Df Sum Sq Mean Sq F value Pr(>F)
## dupTra$Kmer  2    1.2   0.596   0.037  0.964
## Residuals   45  725.6  16.125
## [1] "Pairwise comparison"
```

```
##   Tukey multiple comparisons of means
##     95% family-wise confidence level
##
## Fit: aov(formula = dupTra$Score ~ dupTra$Kmer, data = dupTra2)
##
## $`dupTra$Kmer`
##                 diff      lwr     upr     p adj
## KmerB-KmerA -0.13750 -3.57842 3.30342 0.9948428
## KmerC-KmerA -0.38125 -3.82217 3.05967 0.9610632
## KmerC-KmerB -0.24375 -3.68467 3.19717 0.9838872
```

**Fragmented**

```
## Fragmented genes comparison
```

```
## [1] "All sets are normally distributed"
## [1] "ANOVA"
##              Df Sum Sq Mean Sq F value Pr(>F)
## fragTra$Kmer  2  0.101  0.0506   0.085  0.918
## Residuals    45 26.671  0.5927
## [1] "Pairwise comparison"
```

```
##   Tukey multiple comparisons of means
##     95% family-wise confidence level
##
## Fit: aov(formula = fragTra$Score ~ fragTra$Kmer, data = fragTra2)
##
## $`fragTra$Kmer`
##                 diff        lwr       upr     p adj
## KmerB-KmerA  0.05625 -0.6034308 0.7159308 0.9767433
## KmerC-KmerA -0.05625 -0.7159308 0.6034308 0.9767433
## KmerC-KmerB -0.11250 -0.7721808 0.5471808 0.9103266
```

**Missing**

```
## Missing genes comparison
```

```
## [1] "One (or more) set is not normally distributed"
## [1] "Data not significant. Skipping pairwise comparison"
```

##### By species (all sets)

##### By species (kmer test)

#### CE

**Complete**

```
## Complete genes comparison
```

```
## [1] "All sets are normally distributed"
## [1] "ANOVA"
##             Df Sum Sq Mean Sq F value Pr(>F)
## Kmer         2  0.155  0.0775   0.186  0.831
## Residuals   33 13.735  0.4162
## [1] "Pairwise comparison"
```

```
##   Tukey multiple comparisons of means
##     95% family-wise confidence level
##
## Fit: aov(formula = Score ~ Kmer, data = ceTra2)
##
## $Kmer
##               diff        lwr       upr     p adj
## KmerB-KmerA -0.125 -0.7712788 0.5212788 0.8836821
## KmerC-KmerA  0.025 -0.6212788 0.6712788 0.9950458
## KmerC-KmerB  0.150 -0.4962788 0.7962788 0.8371627
```

**Single**

```
## Single genes comparison
```

```
## [1] "All sets are normally distributed"
## [1] "ANOVA"
##             Df Sum Sq Mean Sq F value Pr(>F)
## Kmer         2   0.26   0.130   0.033  0.967
## Residuals   33 128.44   3.892
## [1] "Pairwise comparison"
```

```
##   Tukey multiple comparisons of means
##     95% family-wise confidence level
##
## Fit: aov(formula = Score ~ Kmer, data = ceTra2)
##
## $Kmer
##                   diff       lwr      upr     p adj
## KmerB-KmerA -0.1666667 -2.143018 1.809685 0.9766873
## KmerC-KmerA  0.0250000 -1.951352 2.001352 0.9994690
## KmerC-KmerB  0.1916667 -1.784685 2.168018 0.9692934
```

**Duplicated**

```
## Duplicated genes comparison
```

```
## [1] "One (or more) set is not normally distributed"
## [1] "Data not significant. Skipping pairwise comparison"
```

**Fragmented**

```
## Fragmented genes comparison
```

```
## [1] "All sets are normally distributed"
## [1] "ANOVA"
##             Df Sum Sq Mean Sq F value Pr(>F)
## Kmer         2  0.020 0.01000   0.102  0.904
## Residuals   33  3.247 0.09841
## [1] "Pairwise comparison"
```

```
##   Tukey multiple comparisons of means
##     95% family-wise confidence level
##
## Fit: aov(formula = Score ~ Kmer, data = ceTra2)
##
## $Kmer
##                      diff        lwr       upr     p adj
## KmerB-KmerA -2.220446e-16 -0.3142535 0.3142535 1.0000000
## KmerC-KmerA -5.000000e-02 -0.3642535 0.2642535 0.9196201
## KmerC-KmerB -5.000000e-02 -0.3642535 0.2642535 0.9196201
```

**Missing**

```
## Fragmented genes comparison
```

```
## [1] "All sets are normally distributed"
## [1] "ANOVA"
##             Df Sum Sq Mean Sq F value Pr(>F)
## Kmer         2  0.105  0.0525   0.239  0.789
## Residuals   33  7.263  0.2201
## [1] "Pairwise comparison"
```

```
##   Tukey multiple comparisons of means
##     95% family-wise confidence level
##
## Fit: aov(formula = Score ~ Kmer, data = ceTra2)
##
## $Kmer
##               diff        lwr       upr     p adj
## KmerB-KmerA  0.125 -0.3449468 0.5949468 0.7921632
## KmerC-KmerA  0.025 -0.4449468 0.4949468 0.9906526
## KmerC-KmerB -0.100 -0.5699468 0.3699468 0.8611037
```

**BUSCO and reads**

```
##   comp.Program comp.Category comp.Score comp.Reads comp.Sample
## 1      Transpi      Complete       85.9 20,345,696         CE1
## 2      Transpi      Complete       86.4 19,948,493         CE2
## 3      Transpi      Complete       85.4 21,276,093         CE3
## 4      Transpi      Complete       85.7 23,993,704         CE4
## 5      Transpi      Complete       85.7 25,110,365         CE5
## 6      Transpi      Complete       84.7 22,967,010         CE6
```

#### DM

**Complete**

```
## Complete genes comparison
```

```
## [1] "All sets are normally distributed"
## [1] "ANOVA"
##             Df Sum Sq Mean Sq F value Pr(>F)
## Kmer         2    0.2   0.086   0.004  0.996
## Residuals   27  655.1  24.262
## [1] "Pairwise comparison"
```

```
##   Tukey multiple comparisons of means
##     95% family-wise confidence level
##
## Fit: aov(formula = Score ~ Kmer, data = dmTra2)
##
## $Kmer
##              diff       lwr      upr     p adj
## KmerB-KmerA -0.13 -5.591714 5.331714 0.9980818
## KmerC-KmerA  0.05 -5.411714 5.511714 0.9997160
## KmerC-KmerB  0.18 -5.281714 5.641714 0.9963261
```

**Single**

```
## Single genes comparison
```

```
## [1] "One (or more) set is not normally distributed"
## [1] "Data not significant. Skipping pairwise comparison"
```

**Duplicated**

```
## Duplicated genes comparison
```

```
## [1] "All sets are normally distributed"
## [1] "ANOVA"
##             Df Sum Sq Mean Sq F value Pr(>F)
## Kmer         2      2     0.8   0.005  0.995
## Residuals   27   4018   148.8
## [1] "Pairwise comparison"
```

```
##   Tukey multiple comparisons of means
##     95% family-wise confidence level
##
## Fit: aov(formula = Score ~ Kmer, data = dmTra2)
##
## $Kmer
##              diff       lwr      upr     p adj
## KmerB-KmerA  0.12 -13.40735 13.64735 0.9997333
## KmerC-KmerA -0.42 -13.94735 13.10735 0.9967385
## KmerC-KmerB -0.54 -14.06735 12.98735 0.9946148
```

**Fragmented**

```
## Fragmented genes comparison
```

```
## [1] "All sets are normally distributed"
## [1] "ANOVA"
##             Df Sum Sq Mean Sq F value Pr(>F)
## Kmer         2   0.06  0.0303   0.021   0.98
## Residuals   27  39.53  1.4639
## [1] "Pairwise comparison"
```

```
##   Tukey multiple comparisons of means
##     95% family-wise confidence level
##
## Fit: aov(formula = Score ~ Kmer, data = dmTra2)
##
## $Kmer
##              diff       lwr      upr     p adj
## KmerB-KmerA  0.06 -1.281604 1.401604 0.9932458
## KmerC-KmerA -0.05 -1.391604 1.291604 0.9953043
## KmerC-KmerB -0.11 -1.451604 1.231604 0.9774936
```

**Missing**

```
## Missing genes comparison
```

```
## [1] "One (or more) set is not normally distributed"
## [1] "Data not significant. Skipping pairwise comparison"
```

**BUSCO and reads**

```
##   comp.Program comp.Category comp.Score comp.Reads comp.Sample
## 1      Transpi      Complete       83.0  8,276,228         DM1
## 2      Transpi      Complete       96.2 47,552,414         DM2
## 3      Transpi      Complete       95.2 45,807,064         DM3
## 4      Transpi      Complete       91.1 29,221,598         DM4
## 5      Transpi      Complete       93.4 46,936,912         DM5
## 6      Transpi      Complete       83.1  8,276,228         DM1
```

#### MM

**Complete**

```
## Complete genes comparison
```

```
## [1] "All sets are normally distributed"
## [1] "ANOVA"
##             Df Sum Sq Mean Sq F value Pr(>F)
## Kmer         2   0.02  0.0103   0.006  0.994
## Residuals   27  50.51  1.8709
## [1] "Pairwise comparison"
```

```
##   Tukey multiple comparisons of means
##     95% family-wise confidence level
##
## Fit: aov(formula = Score ~ Kmer, data = mmTra2)
##
## $Kmer
##              diff       lwr      upr     p adj
## KmerB-KmerA -0.01 -1.526662 1.506662 0.9998527
## KmerC-KmerA  0.05 -1.466662 1.566662 0.9963237
## KmerC-KmerB  0.06 -1.456662 1.576662 0.9947108
```

**Single**

```
## Single genes comparison
```

```
## [1] "One (or more) set is not normally distributed"
## [1] "Data not significant. Skipping pairwise comparison"
```

**Duplicated**

```
## Duplicated genes comparison
```

```
## [1] "One (or more) set is not normally distributed"
## [1] "Data not significant. Skipping pairwise comparison"
```

**Fragmented**

```
## Fragmented genes comparison
```

```
## [1] "All sets are normally distributed"
## [1] "ANOVA"
##             Df Sum Sq Mean Sq F value Pr(>F)
## Kmer         2  0.041  0.0203   0.023  0.978
## Residuals   27 24.329  0.9011
## [1] "Pairwise comparison"
```

```
##   Tukey multiple comparisons of means
##     95% family-wise confidence level
##
## Fit: aov(formula = Score ~ Kmer, data = mmTra2)
##
## $Kmer
##              diff       lwr       upr     p adj
## KmerB-KmerA  0.05 -1.002555 1.1025553 0.9923833
## KmerC-KmerA -0.04 -1.092555 1.0125553 0.9951181
## KmerC-KmerB -0.09 -1.142555 0.9625553 0.9755489
```

**Missing**

```
## Missing genes comparison
```

```
## [1] "One (or more) set is not normally distributed"
## [1] "Data not significant. Skipping pairwise comparison"
```

**BUSCO and reads**

```
##   comp.Program comp.Category comp.Score comp.Reads comp.Sample
## 1      Transpi      Complete       97.1 21,173,335         MM1
## 2      Transpi      Complete       96.7 22,731,971         MM2
## 3      Transpi      Complete       94.6 68,712,046         MM3
## 4      Transpi      Complete       95.9 59,012,049         MM4
## 5      Transpi      Complete       95.7 21,930,527         MM5
## 6      Transpi      Complete       97.2 21,173,335         MM1
```

##### By sample (all sets)
