## Supplementary material for "TransPi – a comprehensive TRanscriptome ANalysiS PIpeline for *de novo* transcriptome assembly": Supplementary_File_5.html

Suppplementary File 5 - TransPi non-model organisms BUSCO


### Suppplementary File 5 - TransPi non-model organisms BUSCO

```
library(reshape2)
library(plotly)
library(dplyr)
```

### busco3\_real

```
csv=read.csv("busco3_real.csv", header=TRUE)
```

##### All BUSCO (all sets)

**Complete**

```
## Complete genes comparison
```

```
## [1] "One (or more) set is not normally distributed"
## [1] "Data not significant. Skipping pairwise comparison"
```

**Single**

```
## Single genes comparison
```

```
## [1] "One (or more) set is not normally distributed"
## [1] "Kruskal-Wallis test was significant (p<.05)"
## [1] "P value"
## [1] 5.665218e-10
## [1] "Pairwise wilcox test"
```

```
##
##  Pairwise comparisons using Wilcoxon rank sum test with continuity correction
##
## data:  sing$Score and sing$Program
##
##         Transpi
## Trinity 5.8e-10
##
## P value adjustment method: BH
```

**Duplicated**

```
## Duplicated genes comparison
```

```
## [1] "One (or more) set is not normally distributed"
## [1] "Kruskal-Wallis test was significant (p<.05)"
## [1] "P value"
## [1] 9.602908e-11
## [1] "Pairwise wilcox test"
```

```
##
##  Pairwise comparisons using Wilcoxon rank sum test with continuity correction
##
## data:  dup$Score and dup$Program
##
##         Transpi
## Trinity 9.8e-11
##
## P value adjustment method: BH
```

**Fragmented**

```
## Fragmented genes comparison
```

```
## [1] "One (or more) set is not normally distributed"
## [1] "Data not significant. Skipping pairwise comparison"
```

**Missing**

```
## Missing genes comparison
```

```
## [1] "One (or more) set is not normally distributed"
## [1] "Data not significant. Skipping pairwise comparison"
```

##### All BUSCO (by reads)

**Only TransPi**

**TEST STATS**

**Complete**

```
## Complete genes comparison
```

**Singe-Copy**

```
## Singe-Copy genes comparison
```

**Duplicated**

```
## Duplicated genes comparison
```

**Fragmented**

```
## Fragmented genes comparison
```

**Missing**

```
## Missing genes comparison
```

##### By species (all sets)

**BUSCO and reads**
