## Supplementary material for "TransPi – a comprehensive TRanscriptome ANalysiS PIpeline for *de novo* transcriptome assembly": Supplementary_File_6.html

Suppplementary File 6 - TransPi non-model organisms QUAST


### Suppplementary File 6 - TransPi non-model organisms QUAST

```
library(reshape2)
library(plotly)
library(dplyr)
```

```
csv=read.csv("real_data_quast.csv", header=TRUE)
csvt=read.csv("real_data_quast_trinity.csv", header=TRUE)
```

##### Number of trancripts per sample

##### Number of trancripts >500bp per sample

##### Number of trancripts >1000bp per sample

##### Average length of trancripts per sample

##### Longest trancripts per sample

##### Total length per sample

##### N50 per sample

##### Number of Genes per sample

##### Boxplot of all samples

##### Boxplot of number of transcripts and longest transcripts

##### Boxplot of n500,n1k,genes

##### Boxplot of Transpi vs Trinity

##### Boxplot of Transpi vs Trinity: number of transcripts, n>500, n>1k, genes
