## Supplementary material for "TransPi – a comprehensive TRanscriptome ANalysiS PIpeline for *de novo* transcriptome assembly": Supplementary_File_7.html

Suppplementary File 7 - TransPi non-model organisms Mapping


### Suppplementary File 7 - TransPi non-model organisms Mapping

```
library(reshape2)
library(plotly)
library(dplyr)
```

```
csv=read.csv("sum_map.csv", header=TRUE)
csvb=read.csv("sum_map2.csv", header=TRUE)
```

##### Mapping and number of transcripts by sample

##### Mapping, number of transcripts and BUSCO (complete and single) by sample

##### Bar plot of mapping % and BUSCO (complete and single) by sample

##### Histograms of mapping % and BUSCO (complete and single)
