## Supplementary material for "TransPi – a comprehensive TRanscriptome ANalysiS PIpeline for *de novo* transcriptome assembly": Supplementary_File_8.html

Supplementary File 8 - TransPi non-model organisms assemblers distribution


### Supplementary File 8 - TransPi non-model organisms assemblers distribution

```
library(reshape2)
library(plotly)
library(dplyr)
```

```
csv_preEG=read.csv("real_preEG.csv", header=TRUE)
csv_EG=read.csv("real_EG.csv", header=TRUE)
```

##### Pre EG (by sample)

##### After EG (by sample)

##### Pre EG (combined)

##### After EG (combined)
