## Supplementary material for "TransPi – a comprehensive TRanscriptome ANalysiS PIpeline for *de novo* transcriptome assembly": Supplementary_FIle_9.html

[golden\_dalembert] Nextflow Workflow Report


Nextflow Report


- Summary
- Resources
- Tasks

[golden\_dalembert]

### Nextflow workflow report

#### `[golden_dalembert]` *(resumed run)*

Workflow execution completed successfully!

Run times
:   09-Jun-2021 12:14:07 - 09-Jun-2021 13:04:09
    (duration: **50m 2s**)

35 succeeded

1 cached

0 ignored

0 failed

Nextflow command
:   ```
    nextflow run TransPi.nf --all --outdir /data/ubuntu/all_results -w /data/ubuntu/all_work -profile docker,test -resume
    ```

CPU-Hours
:   `9.0 (0% cached)`

Launch directory
:   `/home/ubuntu/test/TransPi/ALL`

Work directory
:   `/data/ubuntu/all_work`

Project directory
:   `/home/ubuntu/test/TransPi`

Script name
:   `TransPi.nf`

Script ID
:   `6acd9a7ecd53e0f96dbb95815cae36d0`

Workflow session
:   `a0b26b21-fbfb-40da-8855-70152940fb14`

Workflow profile
:   docker,test

Nextflow version
:   version 20.10.0, build 5430 (01-11-2020 15:14 UTC)

## Resource Usage

These plots give an overview of the distribution of resource usage for each process.

#### CPU

- Raw Usage
- % Allocated

#### Memory

- Physical (RAM)
- Virtual (RAM + Disk swap)
- % RAM Allocated

#### Job Duration

- Raw Usage
- % Allocated

#### I/O

- Read
- Write

## Tasks

This table shows information about each task in the workflow. Use the search box on the right
to filter rows for specific values. Clicking headers will sort the table by that value and
scrolling side to side will reveal more columns.

Values shown as:

Human readable
Raw values

(tasks table omitted because the dataset is too big)

Generated by Nextflow, version 20.10.0
